# Prominent members of the human gut microbiota express endo-acting O-glycanases to initiate mucin breakdown

**DOI:** 10.1101/835843

**Authors:** Lucy I. Crouch, Marcelo V. Liberato, Paulina A. Urbanowicz, Arnaud Baslé, Christopher A. Lamb, Christopher J. Stewart, Katie Cooke, Mary Doona, Stephanie Needham, Richard R. Brady, Janet E. Berrington, Katarina Madunic, Manfred Wuhrer, Peter Chater, Jeffery P. Pearson, Robert Glowacki, Eric C. Martens, Fuming Zhang, Robert J. Linhardt, Daniel I. R. Spencer, David N. Bolam

## Abstract

The human gut microbiota (HGM) are closely associated with health, development and disease. The thick intestinal mucus layer, especially in the colon, is the key barrier between the contents of the lumen and the epithelial cells, providing protection against infiltration by the microbiota as well potential pathogens. The upper layer of the colonic mucus is a niche for a subset of the microbiota which utilise the mucin glycoproteins as a nutrient source and mucin grazing by the microbiota appears to play a key role in maintaining barrier function as well as community stability. Despite the importance of mucin breakdown for gut health, the mechanisms by which gut bacteria access this complex glycoprotein are not well understood. The current model for mucin degradation involves exclusively exo-acting glycosidases that sequentially trim monosaccharides from the termini of the glycan chains to eventually allow access to the mucin peptide backbone by proteases. However, this model is in direct contrast to the Sus paradigm of glycan breakdown used by the Bacteroidetes which involves extracellular cleavage of glycans by surface located endo-acting enzymes prior to import of the oligosaccharide products. Here we describe the discovery and characterisation of endo-acting family 16 glycoside hydrolases (GH16s) from prominent mucin degrading gut bacteria that specifically target the oligosaccharide side chains of intestinal mucins from both animals and humans. These endo-acting O-glycanases display β1,4-glactosidase activity and in several cases are surface located indicating they are involved in the initial step in mucin breakdown. The data suggest a new paradigm for mucin breakdown by the microbiota and the endo-mucinases provide a potential tool to explore changes that occur in mucin structure in intestinal disorders such as inflammatory bowel disease and colon cancer.

## Introduction

The human gastrointestinal (GI) tract is home to a large and complex community of microbes known as the gut microbiota, which in the large intestine there is an estimated between 100-1000 trillion bacteria^1^. The mucous layer of the GI tract protects the underlying epithelia from the huge microbial load of mutualists, environmental insults and enteric pathogens.

The GI mucus layer is predominantly composed of gel-forming mucins, which are complex glycoproteins secreted by the epithelial cells^2^. Different mucins are expressed in different mucosal surfaces throughout the body and a complete mucin glycoprotein is at least 50 % O-glycan by mass^3^. In the colon, MUC2 is the most abundant gel-forming mucin and is composed of ~80 % glycan^1^. While the number of different monosaccharides making up mucin are relatively limited, the order in which they can be assembled is hugely variable. This huge heterogeneity between individual O-glycan chains and very complex macromolecules. It is this glycan complexity that provides mucin’s resistance to microbial degradation and contributes to the mucus layers’ protective role^4^. Notably, however, some prominent bacterial members of the microbiota, including certain Bacteroidetes spp. and *Akkermanisa muciniphila*^*5*^, have developed the capacity to graze on mucins^6–9^. This trait is thought to be critical to initial colonisation by the microbiota in a new-born and therefore to the development of a healthy adult microbiota^10^. Mucin grazing also enables survival during the absence of diet-derived glycans^11^ and non-mucin degrading species have been shown to be cross-fed by mucin degraders, contributing to the long-term survival and stability of the microbiota^12,13^. By contrast, aberrant or excess degradation of the mucosal layer by the normal microbiota has recently been linked to enhanced pathogen susceptibility, inflammatory bowel disease and even colorectal cancer^14,11^. Despite the importance to gut health of mucin breakdown by the microbiota, little is known about the molecular details of this process. Current models of mucin degradation propose extracellular sequential trimming of terminal sugars from the O-glycan side chains by exo-acting glycosidases to eventually expose the peptide backbone for proteolysis^15^. However, this extracellular ‘exo-trimming’ model is based only on the activity of currently characterised mucin active enzymes and notably is in direct contrast with the Gram-negative Bacteroidetes Sus-like paradigm for glycan breakdown, which requires a surface endo-acting glycanase to cleave the substrate (polysaccharide or glycoconjugate) into smaller oligosaccharides for uptake by the SusC/D OM complex^16–18^. Here we describe the discovery and biochemical and structural characterisation of enzymes expressed by mucin degrading members of the gut microbiota that are able to specifically cleave the O-glycan chains of a range of different animal and human mucins in an endo-like manner. Many of these endo-O-glycanases are surface located and thus support a model where the initial step of mucin degradation by gut bacteria is the extracellular removal of oligosaccharide chains from the glycoprotein, prior to import of these oligosaccharides for intracellular processing. This model fits the Sus paradigm and significantly enhances our understating of the mechanism of mucin breakdown by the microbiota. Furthermore, the activity displayed by these enzymes suggest they could be exploited as tools to explore changes that occur in mucin glycosylation in intestinal disorders such as IBD and colon cancer.

## Results

### Identification of GH16 enzymes expressed during growth on mucin

Inspection of previously published transcriptomic and proteomic data from four Gram-negative prominent mucin degrading HGM species (*B. thetaiotaomicron, B. fragilis, B. caccae* and *A. mucinphila*) identified genes and proteins which are likely involved in mucin breakdown^8,11,19–22^. These included many exo-acting enzymes from CAZy families (carbohydrate active enzymes; CAZymes) previously identified as involved degradation of O-glycans, and in the case of *Bacteroides* spp., SusCD glycan import apparatus and putative surface glycan binding proteins (pSGBPs; Supplementary Fig. 1-3). Surprisingly, some of the most upregulated CAZymes in all species analysed were from glycoside hydrolase family 16 (GH16). This is unexpected as GH16 enzymes are to date almost exclusively endo-acting and have been predominantly characterised as targeting a variety of marine or terrestrial plant polysaccharides, specifically β1,3 or 1,4 glycosidic bonds of glucans and galactans (Supplementary Fig 4). More specifically, these enzymes are a part of subfamily 3 of this family, which is a large and sequence-diverse subfamily characterised solely as β1,3/4-glucosidases found in Metazoa, Fungi, Archaea and Bacteria^23^. In total nine GH16 enzymes were identified from the four species analysed (Supplementary Fig. 1-3 and Supplementary Table 1). Sequence comparison of these enzymes with GH16 family members from other *Bacteroides* spp. indicates that there may be similar enzymes present in species other than the ones characterised in this report (Supplementary Fig. 5). Furthermore, five of them are predicted lipoproteins and therefore likely cell surface associated (Supplementary Table 2). For details on the genomic context of these genes and phylogenetic analysis see Supplementary Discussion and Supplementary Figs. 5-7.

### The GH16 enzymes are endo-acting mucinases

To explore the activity of the O-glycan-upregulated GH16 enzymes, the recombinant forms of the enzymes were screened against mucin from porcine small intestinal (SI) mucin and porcine stomach mucins (PGM type II and III; Supplementary Fig. 8). Initial analysis by TLC suggested that all nine enzymes were active against both SI and gastric mucins and released a range of products that are larger than monosaccharides from these glycoproteins, suggesting endo-like cleavage of the O-glycan chains. To investigate the identity of these products in more detail, the products were labelled with the fluorophore procainamide and analysed by liquid chromatography-fluorescence-detection-electrospray-mass spectrometry (LC-FLD-ESI-MS) and the glycan structures determined by MS/MS (Fig. 2a). The data show that oligosaccharides are produced by the GH16 enzymes with between 2 and 6 alternating hexose and HexNAc sugars that are likely to be sections of the polyLacNAc repeats that form the repeating unit of O-glycan chains (Fig. 1b). The reducing ends were all hexoses, indicating hydrolysis occurred at β-galactose (α-galactose only occurs in mucins as a terminal sugar in blood group B structures) and the products also had a range of fucose and sulfate decoration, revealing these can be accommodated by the GH16 enzymes. Overall these data suggest the nine GH16 enzymes are all endo-acting β-galactosidases that are active on the O-glycan side chains of mucin. Notably, sialic acid (SA) was never observed as a decoration on any products released by the enzymes, even though SA is present on mucin glycans. These data suggest this terminal sugar decoration cannot be accommodated by the GH16 enzymes and, as a result, the broad acting sialidase BT0455^GH33^ was included in all assays to maximise access of the GH16 enzymes to the mucin chains.

**Figure 1.**
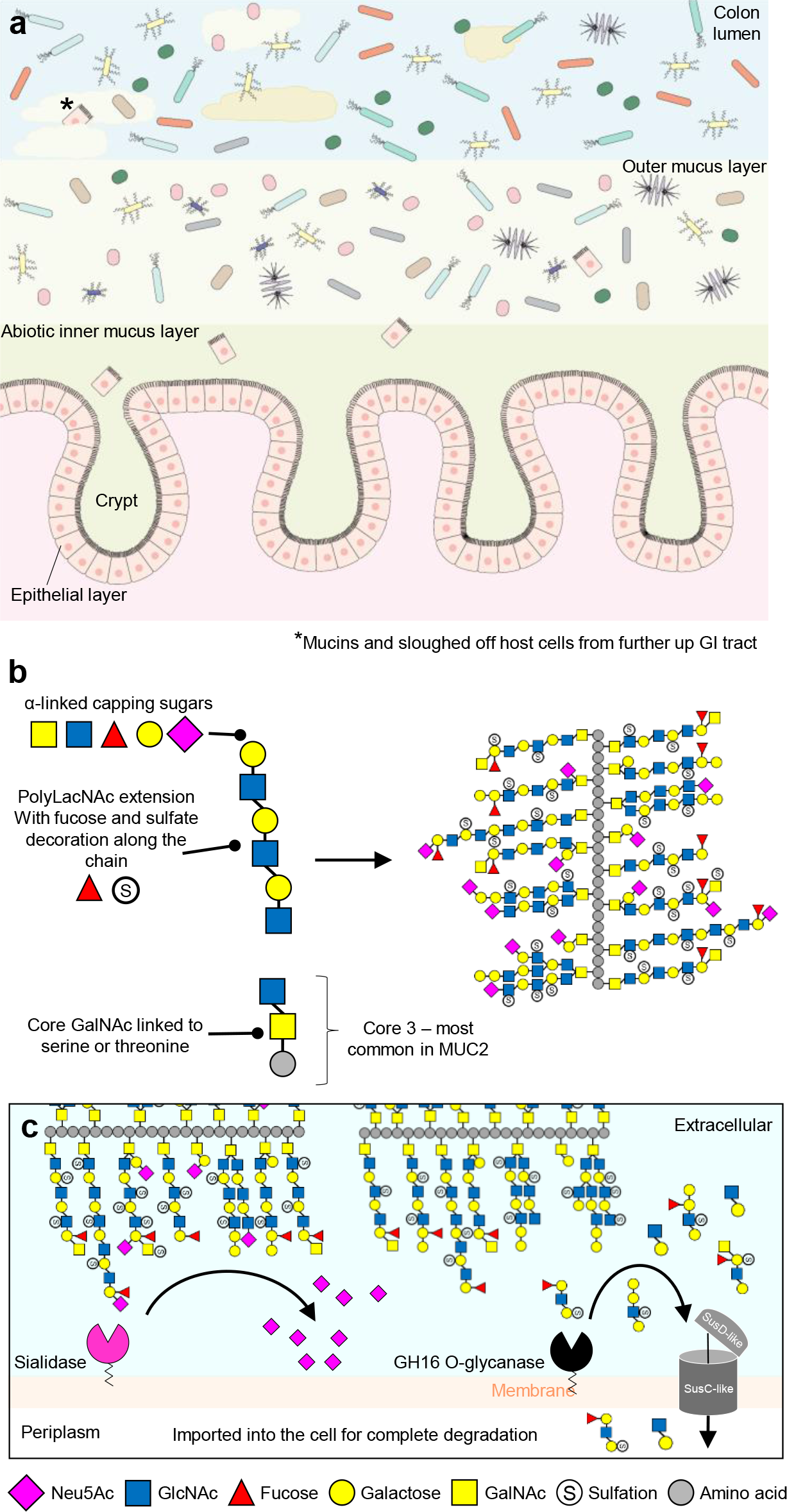
Overview of the mucosal layer in the human colon and the glycan structures in mucins from the GI tract. **a**, A cross section through the colon showing the two layers of the major mucin MUC2. The lower layer is highly viscous and impenetrable to bacteria and protects the epithelial cells. The upper layer thicker, but less viscous, facilitating lubrication of the colon contents and colonisation by a subset of the normal microbiota. **b**, The left hand side shows the core structures that compose a model mucin O-glycan chain. All mucin oligosaccharides are linked via an α-GalNAc to serine and threonine residues in the peptide backbone. The cores are then often extended with polyLacNAc repeats of varying lengths which are decorated along their length by sulfation and fucosylation and capped at the non-reducing end by a variety of α-linked monosaccharides. On the right is a model of an intestinal mucin showing complexity and variability of glycan chains attached to peptide backbone. **c**, A model of the initial steps of O-glycan degradation on the surface of *Bacteroides spp*. Sialic acid is removed by surface-localised sialidase and the GH16 enzyme removes sections of O-glycan for import into the cell for further degradation.

**Figure 2.**
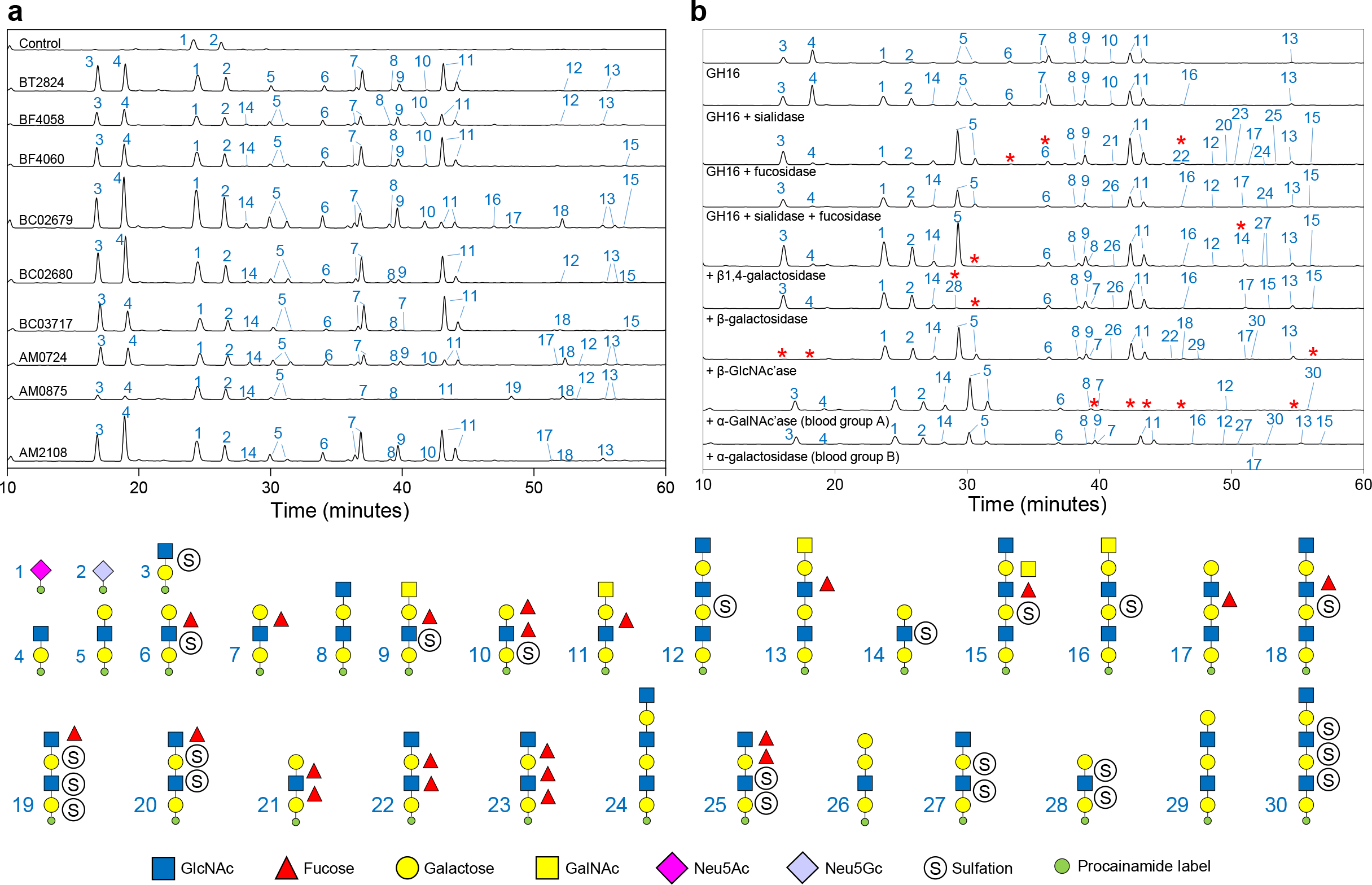
Products of porcine small intestinal mucin digestion by the mucin associated GH16 enzymes. **a**, Products of mucin digestion were labelled with procainamide at the reducing end and analysed by LC-FLD-ESI-MS. From top to bottom the chromatograms correspond to control, BT2824^GH16^, BF4058^GH16^, BF4060^GH16^, Baccac_02679^GH16^, Baccac_02680^GH16^, Baccac_03717^GH16^, Amuc_0724^GH16^, Amuc_0875^GH16^ and Amuc_2108^GH16^, respectively. The locus tags on the figure has been shortened for clarity and labels are shown above each chromatogram. All samples were pre-treated with a broad-acting sialidase (BT0455^GH33^) to maximise access of the GH16 enzymes to the mucin chains. **b**, BF4058^GH16^ products were treated with a series of exo-acting enzymes to provide further insight into the products released. Labels are shown below each chromatogram. Red asterisks highlight peaks that disappear with the addition of an exo-enzyme. See Supplementary Figs. 9, 10 and 19 and Briliūtė *et al.* 2019 for activities of exo-acting enzymes against defined oligosaccharides.

The O-glycan products from the nine GH16 family members from SI mucin varied somewhat, but could be split into two main groups (Fig. 2a). The first group comprised six of the enzymes (BT2824^GH16^, BF4058^GH16^, BF4060^GH16^, Baccac_02680^GH16^, Baccac_03717^GH16^, and Amuc_2108^GH16^) whose products were mainly made up of glycans no more than four sugars in length, although very small amounts of longer oligosaccharides could be detected. The second group composed of Baccac_02679^GH16^, Amuc_0724^GH16^ and Amuc_0875^GH16^ produced a mix of short and longer chain glycans (up to 6 sugars long). Amuc_0875^GH16^ consistently had lowest relative activity against all mucins.

To investigate the structures of the oligosaccharide released by the GH16 enzymes in more detail, the products of BF4058^GH16^ degradation of SI mucin were treated with a series of exo-acting glycosidases of known specificity (Fig. 2b and Supplementary Figs. 9 and 10). With the inclusion of a broad acting α-1,2 fucosidase^24^, there is a different proportion of oligosaccharides, notably, *glycans 7* and *10* disappear and an increase in a relative abundance of *glycans 5* and *6* complements this. A bigger array of larger oligosaccharides are now also observed suggesting that the fucosidase is allowing the GH16 enzyme access to more complex structures. G*lycan 6* shifts in position and these two different resolving times indicate different isomers - for instance this could simply mean a different linkage between two sugars or could be a completely different re-ordering of the monosaccharides within a glycan. The fucosylations that are still present on the glycans produced in the presence of fucosidase are likely to either be inaccessible to this particular enzyme or have either α-1,3 or 4 linkages which are also present in mucins.

Inclusion of further exo-glycosidases with the BF4058^GH16^, sialidase and fucosidase digests reveal further insight into the oligosaccharide structures released by the GH16. The addition of a β1,4-galactosidase (BT0461^GH2^)^25^ removes one of the *glycan 5* peaks, indicating this saccharide is capped with a β1,4-galactose, while both *glycan 5* peaks disappear with the addition of a β1,3/4-galactosidase (BF4061^GH35^; see Supplementary discussion and supplementary Fig. 10), indicating the other *glycan 5* peak is capped with a β1,3-galactose. Interestingly, the β-GlcNAc’ase (BT0459^GH20^)^25^ could degrade the sulfated GlcNAc-Gal disaccharide, although the position of the sulfation in not known so it is unclear at which position the enzyme can accommodate this modification (Fig. 2b, *glycan 3*).

Exo-acting enzymes specific to either the α-GalNAc and α-galactose found on blood group A or B structures, respectively, were also included (Supplementary Fig. 9). No difference in glycans was observed when an α-galactosidase was added, but inclusion of an α-GalNAc’ase revealed several of the larger oligosaccharides could be further degraded, indicating these glycans have a capping α-GalNAc (Fig. 2b, *glycans 9, 11, 13 and 16*). Keratan sulfate is structurally similar to mucin O-glycans in having a repeating polyLacNAc structure with 6S sulfation possible on both the galactose and GlcNAc, but less fucosylation and sialylation than most mucins. The nine GH16 family members were active against egg and bovine corneal keratan sulfate (Supplementary Fig. 11) and the released products indicate that a number of sulfate groups can be tolerated by the enzymes (Supplementary discussion).

A range of defined oligosaccharides were used to further probe the specificity of the O-glycan active GH16 enzymes. TriLacNAc is hydrolysed by nine GH16 enzymes initially produce two trisaccharides, one of which is hydrolysed further to produce GlcNAc and GlcNAcβ1,3Gal. The identity of the products was confirmed using specific exo-acting enzymes (Supplementary Fig. 12). The data revealed that all nine are endo β1,4-galactosidases with a requirement for a β1,3-linked sugar at the −2 position (Supplementary Fig. 12-15). Furthermore, the GH16 enzymes all displayed a preference for GlcNAc over Glc at the +1 site, revealing a discrimination for O-glycans over milk oligosaccharides which are built on a lactose core. Activity on these defined oligosaccharides showed that sulfation and fucosylation are not required for activity. Blood group sugars in the −3’ (fucose) and −4 (GalNAc or Gal) sub-sites are tolerated in most cases but reduce the rate of activity (See Supplementary discussion and Supplementary Table 4).

The activity of the nine recombinant GH16 enzymes was also tested against polysaccharides previously shown to be GH16 substrates (Supplementary Fig. 16). No activity could be detected for agarose, κ-carrageenan, porphyran, pectic galactan, xyloglucan or chitin. However, Amuc_0724^GH16^ displayed significant endo activity against laminarin and weak activity against barley β-glucan and lichenan. BF4060^GH16^, Baccac_02680^GH16^ and Baccac_03717^GH16^ also displayed some very weak activity against laminarin. The possible structural rationale for the activity of Amuc_0724^GH16^ against Glc configured substrates is discussed in the light of structural information and in the Supplementary Discussion. Other non-mucin host polysaccharides are also present in significant amounts in mucosal surfaces, including chondroitin sulfate (CS), heparan sulfate (HS) and hyaluronic acid (HA). The O-glycan active GH16 enzymes were also tested against these and no significant activity could be found, except for small amounts of product released from HS and this is explored in the Supplementary Discussion.

### Crystal structures of O-glycan active GH16 family members

to investigate the basis for O-glycan specificity, structures were obtained for four out of the nine O-glycan active GH16 family members. The apo structures of Baccac 02680^GH16^, Baccac 02680^GH16E143Q^, BACCAC_03717^GH16^ and Amuc0724^GH16^ were obtained to 2.0, 2.1, 2.1 and 2.7 Å, respectively. Structures of BF4060^GH16^ and Baccac_02680^GH16E143Q^ were also obtained with the Galβ1,4GlcNAcβ1,3Gal product present in the negative subsites (despite the latter enzyme being a catalytic mutant) to 3.3 and 2.0 Å (Fig. 3, Supplementary Tables 5 and 6 and Supplementary Figs. 15-17). The electron density of the product allowed us to model the trisaccharide with confidence, even though the occupancies are less than 100 %. All of the GH16 enzymes comprise a β-jellyroll fold, characteristic of the family, consisting of two β-sheets composed of β-strands that form the core fold, which were superimposable with other GH16 structures previously published. A cleft running along the concave surface of the enzymes contains the active site and where the trisaccharide product was bound in the cases of BF4060^GH16^ and Baccac_02680^GH16E143Q^ (Fig. 3a). While the location of the substrate binding site is conserved in the GH16 family, the structures of these clefts vary depending on substrate specificity (Supplementary Fig. 18a). Some form a tight tunnel for linear undecorated glycans (e.g. the agarase from *Zobellia galactanivorans*^26^), others are much more open to accommodate decorations (e.g. the xyloglucanase from *Tropaeolum majus*^27^), while some GH16 enzymes have substrate binding clefts that are curved to optimise binding to highly curved glycans such as laminarin^28^. There is also a single example of a GH16 family member that has evolved a pocket-like active site to recognise a specific disaccharide^29^ (Supplementary Fig. 18b). Substrate specificity in the GH16 family appears to be dictated by the relative size and positon of the loops and short α-helices extending from the β-strands surrounding the substrate binding cleft. These extensions have been likened to fingers that interact with substrate, therefore modulating specificity, and that nomenclature is used herein^30^. For the O-glycanase GH16 enzymes, BF4060^GH16^ and Baccac_02680^GH16E143Q^ have four fingers and BACCAC_03717^GH16^ and Amuc0724^GH16^ have five out of six possible fingers that have been observed previously in other GH16 structures.

**Figure 3.**
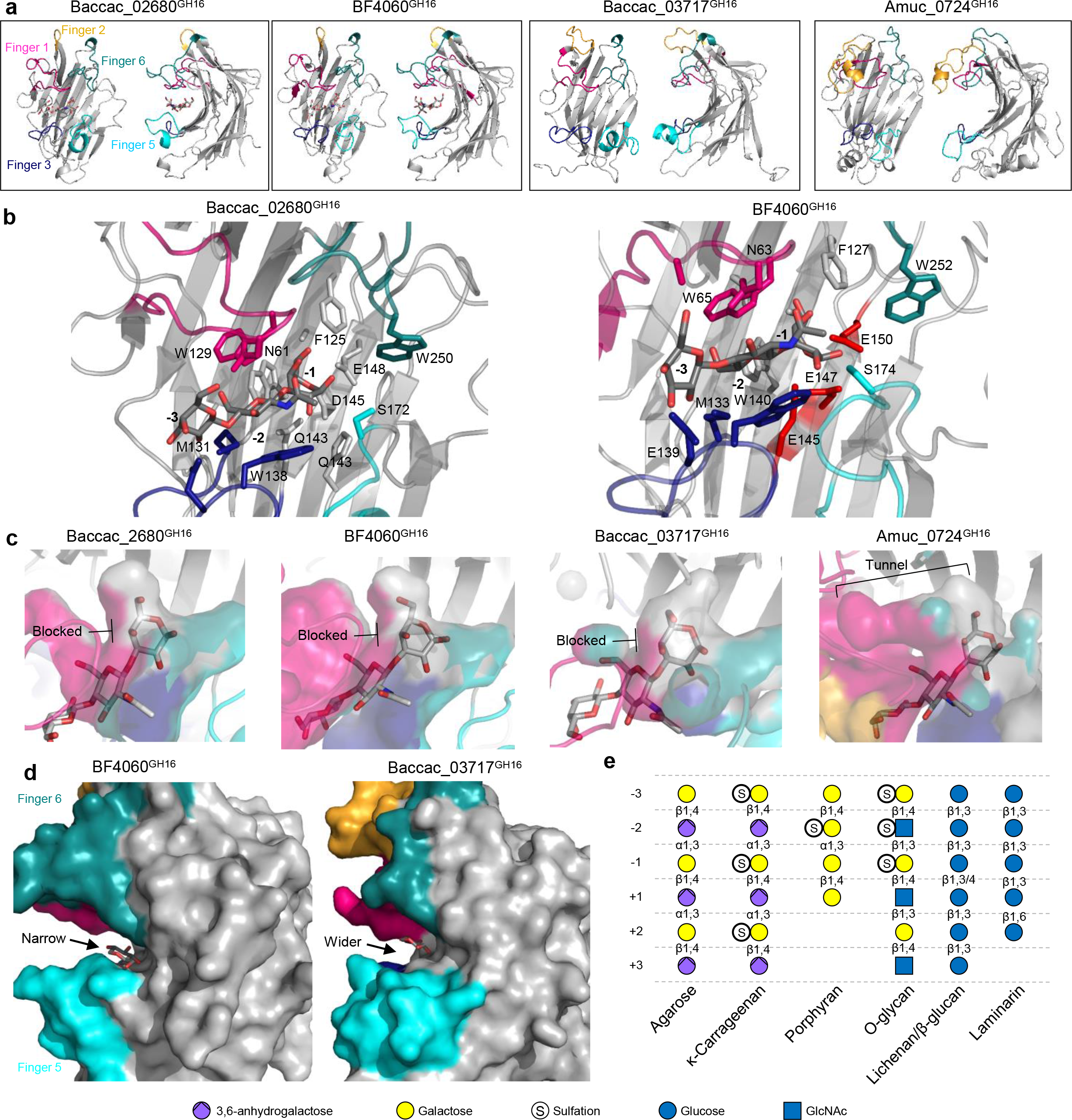
Structures of four of the O-glycan active GH16 family members characterised in this study. **a**, Crystal structures of Baccac_02680^GH16E143Q^, BF4060^GH16^, Baccac_03717^GH16^, and Amuc_0724^GH16^. The loops extending from the active site that are proposed to be involved in substrate specificity in GH16 enzymes are termed ‘fingers’ and are colour coded. **b**, Subsites −1 to −3 of Baccac_02680^GH16E143Q^ and BF4060^GH16^ have the product of TriLacNAc cleavage bound (Galβ1,4GlcNAcβ1,3Gal). The residues interacting directly with sugar are shown as sticks. The aromatic residues shared with β-glucanase GH16 family members that drive specificity for a β1,3 between the −1 and −2 sugars are shown (insert numbers here; See Supplementary Fig. 17 for active sites of Baccac_03717^GH16^, and Amuc_0724^GH16^). **c**, A surface representation of the regions surrounding the −1 subsite showing the selection for the axial O4 of Gal in the three Bacteroides enzymes, while Amuc_0724 has a more open ‘tunnel’ like space that appears to also allow accommodation of the equatorial O4 of Glc. The product from Baccac_02680^GH16E143Q^ was overlaid in the Baccac_03717^GH16^, and Amuc_0724^GH16^ structures. Colours represent the different ‘fingers’. **d**, A view of the predicted positive subsites of BF4060^GH16^ and Baccac_03717^GH16^ overlaid with the glucose from the +1 subsite of a laminarinase from *Phaenerochaete chrysosporium*. The positive subsites are much more closed for BF4060^GH16^ and Baccac_02680^GH16E143Q^ compared to Baccac_03717^GH16^, and Amuc_0724^GH16^ (Supplementary Fig. 17). **e**, An overview of the monosaccharides occupying the different subsites in GH16 family members with different activities. Linkages also shown.

Inspection of the Baccac_02680^GH16E143Q^ and BF4060^GH16^ structures with product reveal most of the interactions between enzyme and ligand are with the Gal at −1 and GlcNAc at −2. The −1 subsite in Baccac_02680^GH16E143Q^ is composed of a number of aromatics, which are also a common feature of the GH16 structures available (Fig. 3b). This enzyme possesses four fingers (numbers 1, 3, 5 and 6) that extend towards the cleft, with fingers 1 and 3 sandwiching the negative subsites and fingers 5 and 6 sandwiching the positive subsites.

Finger 3 contains the sequence motif for GH16 subfamily 3, which consists of three tryptophans interspaced by other residues^23^. BF4060^GH16^ and Baccac_02680^GH16^ are 79 % identical and the structures of these two enzymes are almost identical. In contrast, BACCAC_03717^GH16^ and Amuc0724^GH16^ both have a finger 2 (in addition to 1, 3, 5 and 6) and this has more variable topology of the other fingers (Fig. 3a). For Amuc_0724^GH16^, finger 2 sits over the top of finger 1, but in the BACCAC_03717^GH16^ structure it points away from the cleft. This could reflect the flexibility of finger 2 in this enzyme and could potentially come down over loop 1 in solution like in the Amuc_0724^GH16^ structure or have another role in BACCAC_03717^GH16^. The B-factor putty projections of these structures show finger 2 is dynamic in the BACCAC_03717^GH16^ structure (Supplementary Fig. 17) and alternative conformations of fingers from the crystal structures of other GH16 family members has been observed previously, a finding which is indicative of flexibility^31^.

GH16 family enzymes target a variety of β-glucan and galactan substrates (Supplementary Fig. 4). Glucose and galactose differ only in the hydroxyl group at C4 being equatorial or axial, respectively. For those galactan substrates hydrolysed by the GH16 family, there is also an anhydrogalactose to accommodate in agarose and carrageenan and sulfation for porphyran and carrageenan. Porphyran and carrageenan are 6S and 4S sulfated, respectively, and these decorations would therefore point into the GH16 binding cleft at subsite −2 and −1, respectively. Structural features characteristic to O-glycans include alternating Glc and Gal configured sugars and additionally the presence of GlcNAc, which is not found in other GH16 substrates. In addition, 6S is found on both Gal and GlcNAc and 3S is possible on the galactose at the non-reducing ends of O-glycan chains

There are a number of structural features of the mucin active GH16 family members that indicate a tailoring towards O-glycans as substrates, which include the polyLacNAc chains and also fucose and sulfate decoration. Firstly, in the −1 subsites of the structures from *Bacteroides* spp., the closed space around the O4 hydroxyl explains why only Gal configured sugars can be recognised as the equatorial O4 of glucose would not be accommodated (Fig. 3c). The structure of Amuc_0724^GH16^ in this area is much more open and is a likely explanation for this enzymes additional activity against laminarin (Supplementary Fig. 18). Furthermore, the open space at the O4 in Amuc_0724^GH16^ is a potential pocket for sulfation that the *Bacteroides* spp. enzymes would not be able to accommodate (Supplementary Discussion). Phylogenetic analysis reveals the mucin active GH16 enzymes are likely to have derived from β-glucanases rather than β-galactanases (Supplementary Fig. 6), but in the −1 subsite the selection is for galactose rather than glucose, which shows specificity for O-glycan structures. For the GH16 family there is no conserved way of selecting between glucose and galactose and specificity for galactans arises in distinct branches of β-glucanases the phylogenetic tree (Supplementary Fig. 6, for example the endo-β1,3-galactanases), so this is an example of convergent evolution and the side activity seen in Amuc_0724^GH16^ is linked to its evolutionary origin.

Secondly specificity for polyLacNAcs in O-glycans requires a β1,3 linkage between the −1 and −2 sugars and the structural features driving this specificity in the O-glycan active GH16 enzymes are identical to those in the GH16 enzymes specific for mixed linkage β-glucanases. An aromatic residue in the −2 subsite (also a part of the sequence motif from the subfamily) acts as a hydrophobic platform for the GlcNAc at this position and is at 90° relative to an aromatic residue carrying out the same function in the −1. This feature is conserved amongst β-glucanases (not in GH16 enzymes with other activities, see Supplementary Discussion) and is also required also for the degradation of O-glycans. Thirdly, at the −2 subsite where the GlcNAc is accommodated, the N-acetyl group of the sugar faces the solvent. Other non-mucinase GH16 enzymes with tighter clefts would not be able to accommodate this structure. Also in terms of the −2 subsite, overlay of a porphyran product (originally from a porphyranase GH16 structure^32^) into the clefts of the mucinase GH16 enzymes indicates that sulfation on the GlcNAc at C3 could be accommodated within the cleft at the −2 subsite (Supplementary Fig. 19f).

Fourthly, substrate depletion assays support a preference for polyLacNAc chains in the positive subsites (compared to milk oligosaccharides, Supplementary Fig. 13). Although the product complexes reported here do not have sugars in the positive subsites, comparison with previously published GH16-substrates complexes could be used to propose a structural rationale for the preference of GlcNAc at the +1 in the mucin active enzymes (Fig. 3d and Supplementary Discussion). BF4060^GH16^ has a significant preference for triLacNAc over milk oligosaccharides and analysis of the +1 subsite shows a narrow slot where a GlcNAc would insert with the N-acetyl pointing away from the cleft and S174 from Finger 5 would pincer the N-acetyl against finger 6, thus generating specificity for GlcNAc over glucose. The other structures for O-glycan active GH16 enzymes presented here are more accommodating of milk oligosaccharides and have more open +1 subsites (Fig. 3d and Supplementary Discussion).

### The O-glycan active GH16 enzymes target human mucins from health and diseased samples

We examined the activity of these GH16 family members on human-derived O-glycans from three different disease states (Fig. 4). Tissues from two adults suffering from ulcerative colitis (UC) were obtained and preterm tissue samples from infants with NEC were from 4 infants of gestations 26, 27, 28 and 35 weeks. The three most preterm had terminal ileal NEC and the 35-week infant had colonic NEC. We also tested a number of cultured colorectal cancer (CRC) cell lines.

**Figure 4.**
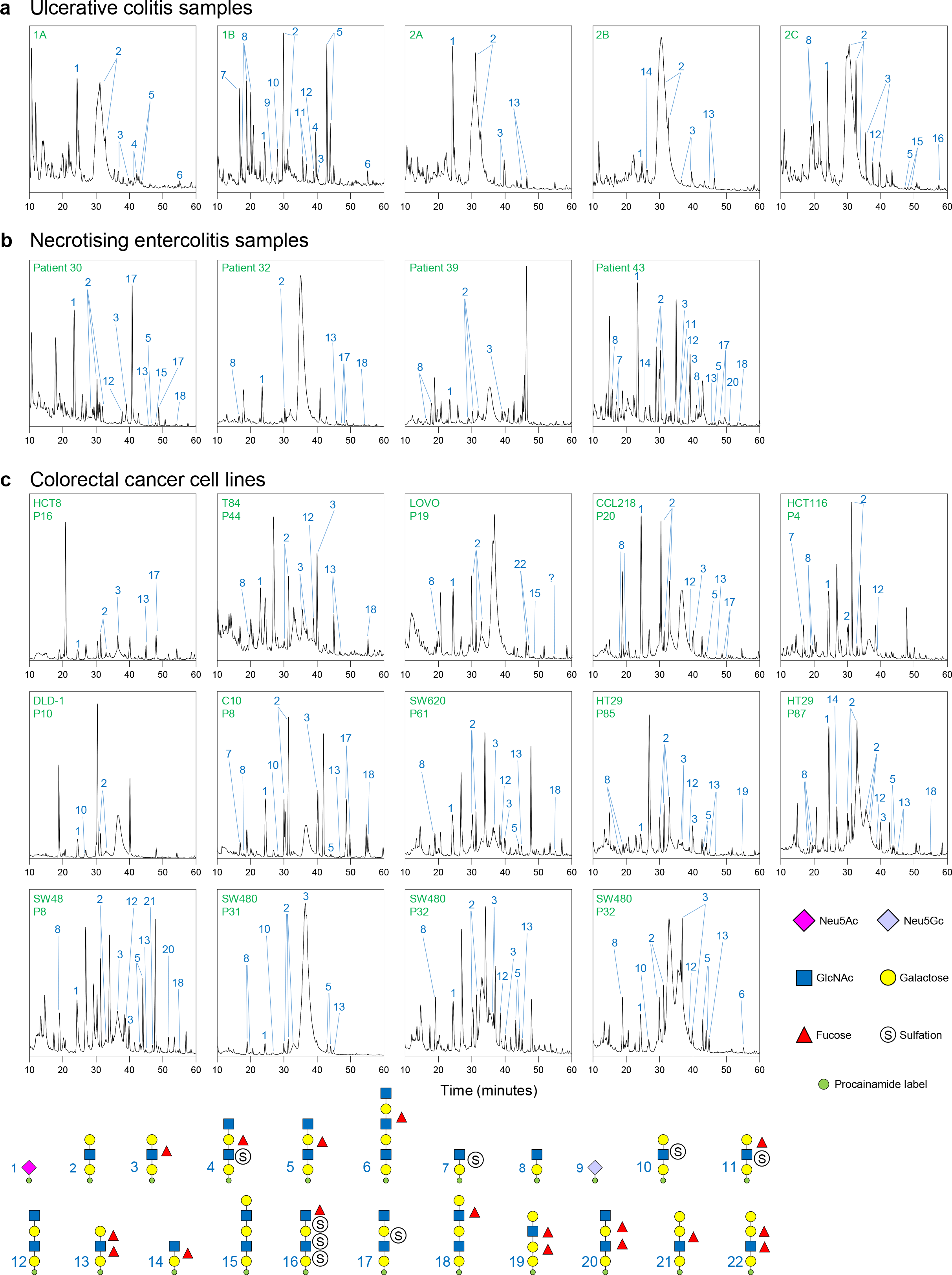
Products released from human mucins by Amuc_0724^GH16^ O-glycanase. Products of mucin digestion were labelled with procainamide at the reducing end and analysed by LC-FLD-ESI-MS. All samples were pre-treated with the broad acting sialidase BT0455^GH33^ **a**, Products of GH16 digestion of mucin samples from two patients with ulcerative colitis. Patient 1 had a laproascopic panproctocolectomy and all removed colon was inflamed. Sample 1A is uninflamed ileum and 1B from inflamed colon. Patient 2 had a laprascopic ileocaecal resection. Sample 2A inflamed ileum (bowel terminal ileum), 2B from uninflamed ileum (near small bowel staple line) and 2C from uninflamed colon (ascending). **b**, Mucin from neonates with necrotising enterocolitis. Each sample is a different patient **c**, Mucin from colorectal cancer cell lines. Small amounts of Neu5Gc are seen in some of the ulcerative colitis samples (e.g. 1A) suggesting either the presence of contaminating dietary animal O-glycans remaining in the mucus layer or that this xenobiotic sugar has been incorporated into human mucins from dietary sources.

Ulcerative colitis (UC) is one form of IDB characterised by an erosion of the mucosal layer^14^. This allows the bacteria to contact the epithelial layer and induce an inflammatory response. Necrotising enterocolitis (NEC) is a condition developed by premature infants where a section of bowel dies, likely linked to an underdeveloped mucosal surface and parts of the innate immune system being not yet active^33^. A complete understanding of all the factors driving these diseases has yet to be full determined. Finally, CRC is the second and fourth deadliest type of cancer in western countries and globally, respectively, and is exacerbated by a ‘Western’ lifestyle, so is likely to become an increasing problem^34,35^. Alterations in the synthesis, secretion and composition of O-glycans in the mucosal surface of the colon have been linked to causation and exasperation of UC and CRC^1,36–41^.

Human O-glycan samples were incubated with Amuc_0724^GH16^ and sialidase BT0455^GH33^, labelled with procainamide and analysed using the same methods as described above. Profiles of GH16 O-glycan products could be produced from all the samples. This work demonstrates that the O-glycan active GH16 family members provide another avenue for researchers to analyse O-glycans in different disease states.

## Discussion

Mucin turnover by the microbiota appears to play a key role in maintaining the normal barrier function of the intestinal mucus layer^11^. Despite the importance of this process, our understanding of the mechanism of mucin breakdown by the microbiota is fragmentary at best. The current model, which is based mainly on the biochemical characterisation of individual enzymes, proposes exo-acting glycosidases trim terminal sugars from the oligosaccharides until the peptide backbone is exposed and can be cleaved by peptidases^15,42^. While this exo-trimming undoubtedly does occur, based on the large size of the intact mucin molecule, this would have to be exclusively outside the cell, thus risking loss of valuable resources to competitors. Furthermore, this extracellular exo-trimming does not fit with the Sus-paradigm for glycan use by Bacteroidetes, which usually involves endo-cleavage of large substrates on the cell surface oligosaccharides that are small enough for import across the outer membrane and further deconstruction in the periplasm^18^. Sus-like systems in Bacteroidetes are encoded within clusters of co-regulated genes known as PULs, with the products of each PUL all being involved in breakdown of a specific glycan. Several mucin-using *Bacteroides* spp. are known to upregulate discrete PULs during growth on mucins, indicating these complex glycoproteins are degraded using Sus-like systems. Here we provide evidence that prominent mucin-using members of the gut microbiota express PUL encoded endo-acting glycanases which target the decorated polyLacNAc structures. Significantly, in all of the *Bacteroides* species studied here, at least one of the GH16 endo-mucinases was a lipoprotein and therefore most likely localised to the cell surface (see Supplementary Table 2). These data support a model in which the endo-mucinases are cleaving oligosaccharide chains from intact mucin at the cell surface and are therefore likely one of the initial steps in mucin breakdown by these bacteria. The oligosaccharide products are then imported via the outer membrane SusC/D apparatus and further degraded in the periplasm by a range of exo-acting enzymes, some of which we also characterise here (e.g. α-fucosidases, β-galactosidases, β-hexosaminidases), but also in some cases by periplasmic GH16 endo-mucinases (e.g. BF4060^GH16^, BACCAC_02680^GH16^). The presence of endo-acting enzymes in the periplasm has been described previously for other glycan breakdown pathways^16^ and may increase the efficiency of exo-cleavage by rapidly generating substrates (i.e. chain ends) for the exo-acting enzymes.

While this model applies to *Bacteroides* spp, it is currently unknown how *Akkermansia* cells access complex glycans, as while they are also Gram negative, there is no evidence of SusC/D genes in the Verrucomicrobium phyla^6^. However, there is direct experimental evidence that at least one of the GH16 mucinases expressed by *A. muciniphila* (Amuc_2108) is localised to the outer membrane during growth on mucin, supporting a similar role to the *Bacteroides* enzymes for the *Akkermansia* GH16 in initiating mucin breakdown. Furthermore, sequence analysis of most of the exo-acting CAZymes expressed by *A. muciniphila* reveal that these enzymes are likely periplasmic, similar to those observed in *Bacteroides* spp. and supporting exo-degradation of the surface GH16 released oligosaccharides in the periplasm of *Akkermasia*.

Although targeting of polyLacNAc structures by the GH16 enzymes is likely an initial step in mucin breakdown, further processing of the remaining mucin would be required. The polyLacNAc side chains are attached directly to different core glycan structures, which are in turn linked to the peptide backbone. One possibility for processing the remaining glycopeptide is that extracellular or surface exo-acting glycosidases remove these core structures, prior to peptidase action on the naked backbone, or that proteases are able to act on the core glycosylated backbone to remove glycopeptides for uptake and further processing. This latter model is supported by recent the recent discovery of a class of glyco-peptidases expressed by gut microbes, including *B. thetaiotaomicron*, that specifically target O-glycosylated mucins^43,44^.

While much of the O-glycan that colonic microbiota will be exposed to will be from MUC2^45^, as the major mucin expressed in the distal intestine, it is worth noting that these bacteria will also come into contact with significant amounts of MUC5AC, MUC5B and MUC6 mucins that have moved down the digestive tract from the saliva, oesophagus and stomach where they originated. In addition to gel-forming mucins, gut microbes will be exposed to membrane-associated mucins that are a part of the apical surface glycocalyx of epithelial cells, especially when dead cells are sloughed off the epithelium throughout the GI tract and these include MUC3, MUC12 and MUC17^46,47^. Furthermore, greater than 80 % of secreted proteins are O-glycosylated and the gut microbiota will come into contact with these from both host and dietary sources^48^. The PGM and SI used in this study indicate theses microbes can access the different types of O-glycans moving through the GI tract.

Overall, the findings reported here contribute significantly towards the understanding about the associations between the host and prominent members of the human gut microbiota. Significantly, we were also able to use these enzymes to produce glycan fragments from mucins derived from patients and cell lines with different disease states. These findings open up the exciting possibility of exploiting this activity for characterisation and detection of biomarkers to allow more effective and earlier diagnosis of diseases such as IBD and CRC, with the possibility of extending the applications to other mucosal surfaces.

## Materials and Methods

### Sources of glycans and glycoproteins

TriLacNAc was purchased from Elicityl and the rest of the defined oligosaccharides were from Carbosynth. PGM II and III (Sigma) was produced by dissolving in DI water at 50 mg/ml and the precipitate removed by centrifugation before assays were carried out (leaving 35-40 mg/ml). Porcine small intestinal mucin was prepared as previously described with the only modification being a double CsCl gradient without Sepharose separation or SDS Page in between^49^. Keratan was prepared as described previously^50^.

### Bacterial strains

The *Bacteroides* strains used were: *B. thetaiotaomicron* VPI-5482, *B. fragilis* NCTC9342, *B. caccae* ATCC43185, *B. cellulosilyticus* DSM14838, *B. finegoldii* DSM17565, *B. vulgatus* ATCC8483, *B. ovatus* ATCC8482, *B. xylanisolvans* XBA1, *B. intestinalis* DSM17393, and *Akkermansia muciniphila* ATCC BAA835/DSM22959.

### Cloning, expression and purification of recombinant proteins

This was carried out as described in^25^.

### Crystallization

The GH16 enzymes were initially screened using commercial kits (Molecular Dimensions and Qiagen). Protein concentrations, crystallisation conditions and cryo-protectant used are given in supplementary table 6. The drops, composed of 0.1 ul of protein solution plus 0.1 or 0.2 ul of reservoir solution, were set up in sitting drop vapour diffusion plates by a Mosquito crystallization robot and incubated at 20 °C. BACCAC_02680^GH16E143Q^ was incubated with 5 mM of ligand for one hour and co-crystallised. BF4060 crystals were soaked in solution containing cryo-protectant and 3.5 mM TriLacNAc for 5 minutes prior to flash cooling in liquid nitrogen.

### Data collection, structure solution, model building, refinement and validation

Data sets were integrated with XDS^51^ or DIALS ^52^ or XIA2 ^53^ and scaled with Aimless^54^. Initial phases were obtained for Amuc_0724^GH16^ by molecular replacement with Molrep ^55^ (REF) using 3WUT and Phaser^56^ using a GH16 laminarinase from *Rhodothermus marinus* as search model (PDB 3ILN) for all the other proteins. Models were refined with refmac^57^ and manual model building with Coot^58^. Final models were validated with MolProbity^59^. The statistics from data processing and refinement can be found in Supplementary Table X. Other software used were from CCP4 suite^60^ of Phenix suite^61^. Figures were made with pymol^62^.

### Growth of bacterial species

All growths were carried out in an anaerobic cabinet (Whitley A35 Workstation; Don Whitley). Glycerol stocks of bacterial were revived overnight in tryptone-yeast-extract-glycerol medium plus haematin^63^. *A. muciniphila* and *B. xylanisolvans* required chopped meat broth (CMB) at this stage instead^11,64^. Monitoring growth against different substrates was done in minimal media for all *Bacteroides* spp.^8^, however for *A. muciniphila* CMB was used without the addition of monosaccharides. For plate growths, a 96-well plate was monitored at 600 nm for 48 h by a Biotek Epoch plate reader. Growth against monosaccharides and PGM II and III (precipitate removed) was carried out at 35 and 40 mg/ml, respectively. Growth against heparan sulfate and chondroitin sulfate was carried out at 20 mg/ml and hyaluronic acid at 10 mg/ml for viscosity reasons.

### Recombinant enzyme assays

For overnight assays, defined oligosaccharides were incubated at a final concentration of 1 mM in the presence of 3 μM of enzyme. For substrate depletion assays, 1 mM oligosaccharides were incubated with 0.1 μM enzyme and samples removed at various times. Some enzymes required increasing to 1 μM to assess the activity against substrates. The concentrations of different substrates are indicated to throughout the figures.

### Thin-layer chromatography

For defined oligosaccharides and other polysaccharides, 3 μl of an assay containing 1 mM substrate was spotted on to silica plates. For assays against mucin, this was increased to 9 μl. The plates were resolved in running buffer containing butanol/acetic acid/water (2:1:1) and stained using a diphenylamine-aniline-phosphoric acid stain^65^.

### Colorectal cancer cell line growth

Human CRC cell lines were obtained from the Department of Surgery of the Leiden University Medical Center (LUMC), Leiden, The Netherlands. The cell lines cultured at the LUMC were kept in Hepes-buffered RPMI 1640 culture medium containing L-glutamine and supplemented with penicillin (5000 IU per mL), streptomycin (5 mg ml^−1^), and 10% (*v/v*) fetal calf serum (FCS). Cells were incubated at 37°C with 5% CO_2_ in humidified air. The cells were harvested after reaching approximately 80% of confluence. To detach the cells from the culture flask a trypsin/EDTA solution in 1X PBS was used. Enzyme activity was stopped using the medium in a ratio 2:5 (trypsin:medium *v/v*). The cells were counted using TC20 automated cell counter from Bio-Rad technologies (California, USA) based on trypan blue staining. The cells were washed twice with 5 mL of 1x PBS, aliquoted to 2.0 × 10^6^ cells per mL of 1x PBS and pelleted by centrifuging 3 min at 1500 x g. Finally, the supernatant was removed, and the cell pellets were stored at −20°C. 2 million cells were used per reaction.

### Human sample collection

IBD tissue samples from two subjects were collected as part of the Newcastle Biobank following written consent according to approval from Newcastle and North Tyneside Research Ethics Committee 1 (REC:17/NE/0361). Matched ileal and colonic samples were obtained from one panproctocolectomy for ulcerative colitis and one ileocaecal resection of Crohn’s disease. Samples were transferred on wet ice directly to the laboratory for mechanical isolation of the mucous layer by gently scraping using a pipette tip. Necrotising enterocolitis samples were collected as part of the ethically approved SERVIS study and Great North Neonatal Biobank (approvals 10/H0908/39 and 15-NE-0334). Fresh tissue was collected from surgically resected specimens when a clinically necessary procedure was taking part, stored briefly in sterile phosphate buffered saline and transported to the laboratory on ice.

### High-performance liquid chromatography with pulsed amphoteric detection (HPAEC-PAD)

To analyse the substrate depletion assays, they were separated using a CARBOPAC PA-100 anion exchange column with a CARBOPAC PA-100 guard. Flow was 1 ml min^−1^ and elution conditions were 0-10 min, 100 mM NaOH; 10-35 min 100 mM NaOH with a 0-166 mM sodium acetate gradient.

### Procainamide labelling

Reducing ends of GH16 products were labelled by reductive amination using a procainamide labelling kit containing sodium cyanoborohydride as reductant (Ludger). Before and after labelling the O-glycan samples were cleaned up using PBM plates and S-cartridges, respectively (Ludger).

### Liquid chromatography-fluorescence detection-electrospray-mass spectrometry analysis of procainamide labelled glycans

Procainamide-labelled samples were analysed by LC-FLD-ESI-MS. 25 μl of sample was injected to a Waters ACQUITY UPLC Glycan BEH Amide column (2.1 × 150 mm, 1.7 μm particle size, 130 Å pore size) at 40 °C on a Dionex Ultimate 3000 UHPLC instrument with a fluorescent detector (λ_ex_= 310 nm, λ_em_= 370 nm) attached to a Bruker Amazon Speed ETD. Mobile phase A was a 50 mM ammonium formate solution (pH 4.4) and mobile phase B was neat acetonitrile. Analyte separation was accomplished by gradients running at a flow rate of 0.4 ml.min^−1^ from 85 to 57 % mobile phase B over 115 min and from 85 to 62 % over 95 min for mucin and keratan samples, respectively. The Amazon speed was operated in the positive sensitivity mode using the following settings: source temperature, 180 °C; gas flow. 41 min^−1^; capillary voltage, 4,500 V; ICC target, 200,000; maximum accumulation time, 50.00 ms; rolling average, 2; number of precursor ions selected, 3; scan mode, enhanced resolution; mass range scanned, 400 to 1,700.

### Analysis of mass spectrometry data

Mass spectrometry of procainamide-labelled glycans was analysed using Bruker Compass Data Analysis Software and GlycoWorkbench^66^. Glycan compositions were elucidated on the basis of MS^2^ fragmentation and previously published data.

### Bioinformatics

Putative signal sequences were identified using SignalP 5.0^67^. Sequence identities were determined using Clustal Omega using full sequences^68^. The IMG database was used to analyse synteny between different species^69^. The CAZy database (www.cazy.org) was used as the main reference for CAZymes^70^. Alignments and phylogenetic trees were completed in SeaView^71^. To determine the boundaries between different modules in a protein Pfam^72^ and SMART^73,74^ were used.

## Supporting information

Supplementary Results

## Acknowledgements

We thank Carl Morland (Newcastle University, UK) for his expert technical assistance. We thank Dr Mirjam Czjzek for her advice on our structural data and interpretation of it and Prof Robert Hirt for his insightful conversations about phylogenetics. We would like to thank Diamond Light Source (Oxfordshire, UK) for beamtime (proposal mx18598) and staff of beamline I03, I04-1 and I24. We are grateful to Newcastle Biobank and NIHR Newcastle Biomedical Research Centre. Dr Jose Muñoz-Muñoz kingly gifted the arabinogalactan substrates used. The NEC samples were collected as part of the ethically approved SERVIS study (REC 10/H0908/39). The colorectal cancer cell lines were from the Department of Surgery, Leiden University Medical Centre (LUMC), Leiden. The work was funded by BBSRC/Innovate UK IB catalyst award ‘Glycoenzymes for Bioindustries’ (BB/M029018/1).

## Contributions

L.I.C. sorted through previously published gene upregulation and protein expression data. L.I.C. and M.V.L. carried out reactions on defined oligosaccharides and commercial polysaccharides available. M.V.L. produced catalytic mutants. L.I.C. and M.V.L. purified proteins for crystallography. M.V.L. and A.B. obtained and harvested crystals, collected data and solved crystal structures. L.I.C. and M.V.L. carried out comparisons of crystal structures with ones already present in the database. P.A.U. carried out the LC-MS. L.I.C. analysed the LC-MS data. P.C. and J.P.P. collected and purified porcine small intestinal mucin. F.Z. and R.J.L. purified keratan sulfate. R.G and E.C.M supplied the *Bt* PUL knockout strains. L.I.C. carried out substrate depletion assays, carried out growth experiments with different bacterial species, completed the phylogenetic analysis, carried out assays against human samples and assays of with other enzymes used in the report against defined oligosaccharides. C.A.L., R.R.B., M.D., and S.N. were responsible for ethical approval, governance, patient identification and sample collection for IBD tissues. R.R.B. performed surgery where adult intestinal samples were collected. K.C. and C.A.L. were responsible for lab preparation of IBD tissue. C.J.S. and J.E.B were responsible for ethical approval and provision of samples for the preterm neonate NEC samples. J.E.B performed the neonate surgery. K.M. prepared the colorectal cancer cells. L.I.C. and D.N.B designed the experiments, analysed the data and wrote the manuscript.

## Data availability

The crystal structures are deposited in the Protein Data Bank under the accession numbers 6T2N, 6T2O, 6T2P, 6T2Q, 6T2R and 6T2S. The other data that supports the findings in this paper are available upon request from the corresponding authors.

## References

1 Johansson, M. E., Sjovall, H. & Hansson, G. C. The gastrointestinal mucus system in health and disease. Nature reviews. Gastroenterology & hepatology 10, 352–361, doi:10.1038/nrgastro.2013.35 (2013).

2 Lang, T., Hansson, G. C. & Samuelsson, T. Gel-forming mucins appeared early in metazoan evolution. Proceedings of the National Academy of Sciences of the United States of America 104, 16209–16214, doi:10.1073/pnas.0705984104 (2007).

3 Johansson, M. E. & Hansson, G. C. Immunological aspects of intestinal mucus and mucins. Nature reviews. Immunology 16, 639–649, doi:10.1038/nri.2016.88 (2016).

4 Larsson, J. M., Karlsson, H., Sjovall, H. & Hansson, G. C. A complex, but uniform O-glycosylation of the human MUC2 mucin from colonic biopsies analyzed by nanoLC/MSn. Glycobiology 19, 756–766, doi:10.1093/glycob/cwp048 (2009).

5 Forster, S. C. et al. A human gut bacterial genome and culture collection for improved metagenomic analyses. Nature biotechnology 37, 186–192, doi:10.1038/s41587-018-0009-7 (2019).

6 Derrien, M., Vaughan, E. E., Plugge, C. M. & de Vos, W. M. Akkermansia muciniphila gen. nov., sp. nov., a human intestinal mucin-degrading bacterium. Int J Syst Evol Microbiol 54, 1469–1476, doi:10.1099/ijs.0.02873-0 (2004).

7 Martens, E. C., Roth, R., Heuser, J. E. & Gordon, J. I. Coordinate regulation of glycan degradation and polysaccharide capsule biosynthesis by a prominent human gut symbiont. The Journal of biological chemistry 284, 18445–18457, doi:10.1074/jbc.M109.008094 (2009).

8 Martens, E. C., Chiang, H. C. & Gordon, J. I. Mucosal glycan foraging enhances fitness and transmission of a saccharolytic human gut bacterial symbiont. Cell host & microbe 4, 447–457, doi:10.1016/j.chom.2008.09.007 (2008).

9 Martens, E. C. et al. Recognition and degradation of plant cell wall polysaccharides by two human gut symbionts. PLoS Biol 9, e1001221, doi:10.1371/journal.pbio.1001221 (2011).

10 Marcobal, A. et al. Bacteroides in the infant gut consume milk oligosaccharides via mucus-utilization pathways. Cell host & microbe 10, 507–514, doi:10.1016/j.chom.2011.10.007 (2011).

11 Desai, M. S. et al. A Dietary Fiber-Deprived Gut Microbiota Degrades the Colonic Mucus Barrier and Enhances Pathogen Susceptibility. Cell 167, 1339–1353.e1321, doi:10.1016/j.cell.2016.10.043 (2016).

12 Egan, M. et al. Cross-feeding by Bifidobacterium breve UCC2003 during co-cultivation with Bifidobacterium bifidum PRL2010 in a mucin-based medium. BMC microbiology 14, 282, doi:10.1186/s12866-014-0282-7 (2014).

13 Schroeder, B. O. et al. Bifidobacteria or Fiber Protects against Diet-Induced Microbiota-Mediated Colonic Mucus Deterioration. Cell host & microbe 23, 27–40.e27, doi:10.1016/j.chom.2017.11.004 (2018).

14 Corfield, A. P. The Interaction of the Gut Microbiota with the Mucus Barrier in Health and Disease in Human. Microorganisms 6, doi:10.3390/microorganisms6030078 (2018).

15 Marcobal, A., Southwick, A. M., Earle, K. A. & Sonnenburg, J. L. A refined palate: bacterial consumption of host glycans in the gut. Glycobiology 23, 1038–1046, doi:10.1093/glycob/cwt040 (2013).

16 Cuskin, F. et al. Human gut Bacteroidetes can utilize yeast mannan through a selfish mechanism. Nature 517, 165–169, doi:10.1038/nature13995 (2015).

17 Koropatkin, N. M., Martens, E. C., Gordon, J. I. & Smith, T. J. Starch catabolism by a prominent human gut symbiont is directed by the recognition of amylose helices. Structure (London, England : 1993) 16, 1105–1115, doi:10.1016/j.str.2008.03.017 (2008).

18 Rogowski, A. et al. Glycan complexity dictates microbial resource allocation in the large intestine. Nature communications 6, 7481, doi:10.1038/ncomms8481 (2015).

19 Pudlo, N. A. et al. Symbiotic Human Gut Bacteria with Variable Metabolic Priorities for Host Mucosal Glycans. MBio 6, e01282–01215, doi:10.1128/mBio.01282-15 (2015).

20 Ottman, N. et al. Genome-Scale Model and Omics Analysis of Metabolic Capacities of Akkermansia muciniphila Reveal a Preferential Mucin-Degrading Lifestyle. Applied and environmental microbiology 83, doi:10.1128/aem.01014-17 (2017).

21 Ottman, N. et al. Characterization of Outer Membrane Proteome of Akkermansia muciniphila Reveals Sets of Novel Proteins Exposed to the Human Intestine. Frontiers in microbiology 7, 1157, doi:10.3389/fmicb.2016.01157 (2016).

22 Shin, J. et al. Elucidation of Akkermansia muciniphila Probiotic Traits Driven by Mucin Depletion. Frontiers in microbiology 10, 1137, doi:10.3389/fmicb.2019.01137 (2019).

23 Viborg, A. H. et al. A subfamily roadmap for functional glycogenomics of the evolutionarily diverse Glycoside Hydrolase Family 16 (GH16). The Journal of biological chemistry, doi:10.1074/jbc.RA119.010619 (2019).

24 Katayama, T. et al. Molecular cloning and characterization of Bifidobacterium bifidum 1,2-alpha-L-fucosidase (AfcA), a novel inverting glycosidase (glycoside hydrolase family 95). Journal of bacteriology 186, 4885–4893, doi:10.1128/jb.186.15.4885-4893.2004 (2004).

25 Briliute, J. et al. Complex N-glycan breakdown by gut Bacteroides involves an extensive enzymatic apparatus encoded by multiple co-regulated genetic loci. Nature microbiology 4, 1571–1581, doi:10.1038/s41564-019-0466-x (2019).

26 Hehemann, J. H. et al. Biochemical and structural characterization of the complex agarolytic enzyme system from the marine bacterium Zobellia galactanivorans. The Journal of biological chemistry 287, 30571–30584, doi:10.1074/jbc.M112.377184 (2012).

27 Mark, P. et al. Analysis of nasturtium TmNXG1 complexes by crystallography and molecular dynamics provides detailed insight into substrate recognition by family GH16 xyloglucan endo-transglycosylases and endo-hydrolases. Proteins 75, 820–836, doi:10.1002/prot.22291 (2009).

28 Labourel, A. et al. The beta-glucanase ZgLamA from Zobellia galactanivorans evolved a bent active site adapted for efficient degradation of algal laminarin. The Journal of biological chemistry 289, 2027–2042, doi:10.1074/jbc.M113.538843 (2014).

29 Tempel, W. et al. Three-dimensional structure of GlcNAcalpha1-4Gal releasing endo-beta-galactosidase from Clostridium perfringens. Proteins 59, 141–144, doi:10.1002/prot.20363 (2005).

30 Matard-Mann, M. et al. Structural insights into marine carbohydrate degradation by family GH16 kappa-carrageenases. The Journal of biological chemistry 292, 19919–19934, doi:10.1074/jbc.M117.808279 (2017).

31 Jeng, W. Y., Wang, N. C., Lin, C. T., Shyur, L. F. & Wang, A. H. Crystal structures of the laminarinase catalytic domain from Thermotoga maritima MSB8 in complex with inhibitors: essential residues for beta-1,3-and beta-1,4-glucan selection. The Journal of biological chemistry 286, 45030–45040, doi:10.1074/jbc.M111.271213 (2011).

32 Hehemann, J. H. et al. Transfer of carbohydrate-active enzymes from marine bacteria to Japanese gut microbiota. Nature 464, 908–912, doi:10.1038/nature08937 (2010).

33 Hsueh, W. et al. Neonatal necrotizing enterocolitis: clinical considerations and pathogenetic concepts. Pediatric and developmental pathology : the official journal of the Society for Pediatric Pathology and the Paediatric Pathology Society 6, 6–23, doi:10.1007/s10024-002-0602-z (2003).

34 Brody, H. Colorectal cancer. Nature 521, S1, doi:10.1038/521S1a (2015).

35 Zamani, M., Hosseini, S. V. & Mokarram, P. Epigenetic biomarkers in colorectal cancer: premises and prospects. Biomarkers : biochemical indicators of exposure, response, and susceptibility to chemicals 23, 105–114, doi:10.1080/1354750x.2016.1252961 (2018).

36 Gao, N. et al. Loss of intestinal O-glycans promotes spontaneous duodenal tumors. American journal of physiology. Gastrointestinal and liver physiology 311, G74–83, doi:10.1152/ajpgi.00060.2016 (2016).

37 Bergstrom, K. S. & Xia, L. Mucin-type O-glycans and their roles in intestinal homeostasis. Glycobiology 23, 1026–1037, doi:10.1093/glycob/cwt045 (2013).

38 Velcich, A. et al. Colorectal cancer in mice genetically deficient in the mucin Muc2. Science (New York, N.Y.) 295, 1726–1729, doi:10.1126/science.1069094 (2002).

39 Van der Sluis, M. et al. Muc2-deficient mice spontaneously develop colitis, indicating that MUC2 is critical for colonic protection. Gastroenterology 131, 117–129, doi:10.1053/j.gastro.2006.04.020 (2006).

40 Fu, J. et al. Loss of intestinal core 1-derived O-glycans causes spontaneous colitis in mice. The Journal of clinical investigation 121, 1657–1666, doi:10.1172/jci45538 (2011).

41 An, G. et al. Increased susceptibility to colitis and colorectal tumors in mice lacking core 3-derived O-glycans. The Journal of experimental medicine 204, 1417–1429, doi:10.1084/jem.20061929 (2007).

42 Tailford, L. E., Crost, E. H., Kavanaugh, D. & Juge, N. Mucin glycan foraging in the human gut microbiome. Frontiers in genetics 6, 81, doi:10.3389/fgene.2015.00081 (2015).

43 Noach, I. et al. Recognition of protein-linked glycans as a determinant of peptidase activity. Proceedings of the National Academy of Sciences of the United States of America 114, E679–e688, doi:10.1073/pnas.1615141114 (2017).

44 Nakjang, S., Ndeh, D. A., Wipat, A., Bolam, D. N. & Hirt, R. P. A novel extracellular metallopeptidase domain shared by animal host-associated mutualistic and pathogenic microbes. PloS one 7, e30287, doi:10.1371/journal.pone.0030287 (2012).

45 Gum, J. R., Jr., Hicks, J. W., Toribara, N. W., Siddiki, B. & Kim, Y. S. Molecular cloning of human intestinal mucin (MUC2) cDNA. Identification of the amino terminus and overall sequence similarity to prepro-von Willebrand factor. The Journal of biological chemistry 269, 2440–2446 (1994).

46 Schneider, H. et al. The human transmembrane mucin MUC17 responds to TNFalpha by increased presentation at the plasma membrane. The Biochemical journal, doi:10.1042/bcj20190180 (2019).

47 Hansson, G. C. Mucus and mucins in diseases of the intestinal and respiratory tracts. J Intern Med, doi:10.1111/joim.12910 (2019).

48 Steentoft, C. et al. Precision mapping of the human O-GalNAc glycoproteome through SimpleCell technology. The EMBO journal 32, 1478–1488, doi:10.1038/emboj.2013.79 (2013).

49 Fogg, F. J. et al. Characterization of pig colonic mucins. The Biochemical journal 316 (Pt 3), 937–942, doi:10.1042/bj3160937 (1996).

50 Fu, L. et al. Keratan sulfate glycosaminoglycan from chicken egg white. Glycobiology 26, 693–700, doi:10.1093/glycob/cww017 (2016).

51 Kabsch, W. XDS. Acta crystallographica. Section D, Biological crystallography 66, 125–132, doi:10.1107/s0907444909047337 (2010).

52 Winter, G. et al. DIALS: implementation and evaluation of a new integration package. Acta crystallographica. Section D, Structural biology 74, 85–97, doi:10.1107/s2059798317017235 (2018).

53 Winter, G., Lobley, C. M. & Prince, S. M. Decision making in xia2. Acta crystallographica. Section D, Biological crystallography 69, 1260–1273, doi:10.1107/s0907444913015308 (2013).

54 Evans, P. Scaling and assessment of data quality. Acta crystallographica. Section D, Biological crystallography 62, 72–82, doi:10.1107/s0907444905036693 (2006).

55 Vagin, A. & Teplyakov, A. Molecular replacement with MOLREP. Acta crystallographica. Section D, Biological crystallography 66, 22–25, doi:10.1107/s0907444909042589 (2010).

56 McCoy, A. J. et al. Phaser crystallographic software. J Appl Crystallogr 40, 658–674, doi:10.1107/s0021889807021206 (2007).

57 Murshudov, G. N. et al. REFMAC5 for the refinement of macromolecular crystal structures. Acta crystallographica. Section D, Biological crystallography 67, 355–367, doi:10.1107/s0907444911001314 (2011).

58 Emsley, P., Lohkamp, B., Scott, W. G. & Cowtan, K. Features and development of Coot. Acta crystallographica. Section D, Biological crystallography 66, 486–501, doi:10.1107/s0907444910007493 (2010).

59 Chen, V. B. et al. MolProbity: all-atom structure validation for macromolecular crystallography. Acta crystallographica. Section D, Biological crystallography 66, 12–21, doi:10.1107/s0907444909042073 (2010).

60 The CCP4 suite: programs for protein crystallography. Acta crystallographica. Section D, Biological crystallography 50, 760–763, doi:10.1107/s0907444994003112 (1994).

61 Adams, P. D. et al. PHENIX: a comprehensive Python-based system for macromolecular structure solution. Acta crystallographica. Section D, Biological crystallography 66, 213–221, doi:10.1107/s0907444909052925 (2010).

62 The PyMOL Molecular Graphics System, Version 2.0 Schrödinger, LLC. .

63 Larsbrink, J. et al. A discrete genetic locus confers xyloglucan metabolism in select human gut Bacteroidetes. Nature 506, 498–502, doi:10.1038/nature12907 (2014).

64 Hehemann, J. H., Kelly, A. G., Pudlo, N. A., Martens, E. C. & Boraston, A. B. Bacteria of the human gut microbiome catabolize red seaweed glycans with carbohydrate-active enzyme updates from extrinsic microbes. Proceedings of the National Academy of Sciences of the United States of America 109, 19786–19791, doi:10.1073/pnas.1211002109 (2012).

65 Zhang, Z., Xie, J., Zhang, F. & Linhardt, R. J. Thin-layer chromatography for the analysis of glycosaminoglycan oligosaccharides. Anal Biochem 371, 118–120, doi:10.1016/j.ab.2007.07.003 (2007).

66 Ceroni, A. et al. GlycoWorkbench: a tool for the computer-assisted annotation of mass spectra of glycans. J Proteome Res 7, 1650–1659, doi:10.1021/pr7008252 (2008).

67 Almagro Armenteros, J. J. et al. SignalP 5.0 improves signal peptide predictions using deep neural networks. Nature biotechnology 37, 420–423, doi:10.1038/s41587-019-0036-z (2019).

68 Sievers, F. et al. Fast, scalable generation of high-quality protein multiple sequence alignments using Clustal Omega. Mol Syst Biol 7, 539, doi:10.1038/msb.2011.75 (2011).

69 Markowitz, V. M. et al. IMG: the Integrated Microbial Genomes database and comparative analysis system. Nucleic acids research 40, D115–122, doi:10.1093/nar/gkr1044 (2012).

70 Lombard, V., Golaconda Ramulu, H., Drula, E., Coutinho, P. M. & Henrissat, B. The carbohydrate-active enzymes database (CAZy) in 2013. Nucleic acids research 42, D490–495, doi:10.1093/nar/gkt1178 (2014).

71 Gouy, M., Guindon, S. & Gascuel, O. SeaView version 4: A multiplatform graphical user interface for sequence alignment and phylogenetic tree building. Molecular biology and evolution 27, 221–224, doi:10.1093/molbev/msp259 (2010).

72 El-Gebali, S. et al. The Pfam protein families database in 2019. Nucleic acids research, doi:10.1093/nar/gky995 (2018).

73 Letunic, I. & Bork, P. 20 years of the SMART protein domain annotation resource. Nucleic acids research 46, D493–d496, doi:10.1093/nar/gkx922 (2018).

74 Letunic, I., Doerks, T. & Bork, P. SMART: recent updates, new developments and status in 2015. Nucleic acids research 43, D257–260, doi:10.1093/nar/gku949 (2015).

