## Supplementary Results for "Prominent members of the human gut microbiota express endo-acting O-glycanases to initiate mucin breakdown"

**Supplementary Information**

**Mucin substrates and structures**

Commercially available PGM substrates dissolved in pure water were too opaque to monitor growth through optical density. However, centrifugation to remove precipitate produced a substrate that could be used in for growth experiments. 70 and 80 % of the original dry mass remained in the soluble portion of PGM type II and III, respectively. Incubations of an O-glycan active enzyme against the precipitate and soluble fractions showed that the majority of accessible substrate was in the soluble fractions (Supplementary Fig. 20). The soluble fraction and precipitate were also incubated with polysaccharide lyases previously characterised or predicted to have activity against HS, CS and HA[1-3]. The results indicate that these polysaccharides also remain in the soluble fraction (Supplementary Fig. 20)

The mucous surface of the colon is composed of two layers. The dense inner mucus layer of the colon is an abiotic environment formed by MUC2 still attached to the luminal epithelial cells to protect them from any contact with the HGM. The upper layer is formed from MUC2 released from the inner layer and is niche to some species of the HGM that can access host glycans as a food source[4]. The upper layer is renewed from the inner layer every 1-2 hours[5,6] and there is a close association between this process and commensal microbes, which has been demonstrated in germ-free mice where the mucin was observed to remain attached to the goblet cell rather than being released[7]. This indicates that the HGM promotes the production of healthy mucus barrier and highlights the mutualistic relationship between host and HGM[8].

Previous characterisation of the O-glycans of MUC2 from human sigmoid colon samples showed over 100 different structures, but general trends included mono- to tri-sialyation, predominantly core 3 structures, sulfation predominantly on galactose, fucose predominantly on GlcNAc (both Lewis a and x structures, decorating the chains and not capping), low occurrence of blood group sugars and elongation of up to three LacNAcs[9]. Blood group epitopes have been found to be more common in other mucins, such as salivary, respiratory and cervical[10-12]. This means that the variability would be relatively low between individuals and allow for selection of particular mutualists. The stomach mucin is predominantly Muc5AC and Muc6 and both are characterised as having a capping α1,4-GlcNAc[13], but only Muc5AC has Lewis b structures[14]. A large diversity in chain length and composition of gastric O-glycans has also been noted, with many structures seemingly specific to an individual[15].

**Gene upregulation and protein expression data sets used in the literature**

Two gene upregulation data sets were used for *B. thetaiotamicron* from two different points of growth (early and late phase) on mucin O-glycans from PGM type III relative to glucose (Supplementary Fig. 1)[3]. Single gene upregulation data sets were included for *B. fragilis* and *B. caccae* also grown on mucin O-glycans from PGM III relative to glucose[16,17] (Supplementary Fig. 2). Five different datasets were used to look at upregulated genes from *A. muciniphila* (Supplementary Fig. 3). The different reports containing the datasets state that either strain ATCC BAA835 or DSM22959 were used but these are the same strains from different banks. There were two data sets originated from the same report, one recorded gene fold upregulation between *A. muciniphila* growth on GlcNAc and purified mucin O-glycans from PGM III and also the gene fold upregulation in *A. muciniphila* between a fibre-rich diet and a fibre-free diet in mice[17]. Two data sets used purified PGM III, but with the O-glycans still attached to the protein (one against glucose and one GlcNAc)[18,19]. The final two data sets were proteomics of cells grown on PGM III versus glucose grown cells and the proteomics of the outer-membrane of these cells[20].

**Genomic context of mucin upregulated GH16 family members**

The upregulated GH16 family member from *Bt* is in a PUL with two SusCD pairs, two predicted SGBPs and a GH from family 18 (Supplementary Fig. 5). The two mucin upregulated GH16 family members from *B. fragilis* ($BF4058^{GH16}$ and $BF4060^{GH16}$) are in a PUL that included a SusCD pair and two more predicted CAZymes, BF4059 and BF4061, belonging to the GH20 and GH35 families, respectively. There are three GH16 family members upregulated on O-glycans from *B. caccae* and two of these ($BACCAC\_02679^{GH16}$ and $Baccac\_02680^{GH16}$) are adjacent to each other and are close homologues to those found in *B. fragilis*. $Baccac\_03717^{GH16}$ is the third mucin upregulated GH16 from *B. caccac*. All *B. caccae* GH16 family members are in PULs that contain SusCD pairs. In the *A. muciniphila* genome the glycan degrading apparatus is not organised in to PULs and none of the three GH16 genes upregulated during growth on mucin ($Amuc\_0724^{GH16}$, $Amuc\_0875^{GH16}$ and $Amuc2108^{GH16}$) are near other predicted CAZyme genes[17,18,20].

**Sequence identity and molecular architecture of the GH16 enzymes**

The sequence identity between the mucin upregulated GH16 family members was relatively low with values of between 24-34 % (Supplementary Table 1). The exceptions to this were two pairs of *B. fragilis* and *B. caccae* enzymes in homologous PULs: $BF4058^{GH16}$ and $BC02679^{GH16}$ and $BF4060^{GH16}$ and $BC2680^{GH16}$ with 87 and 79 % identity, respectively. All enzymes were predicted to be composed of a single catalytic module only, with the exception of $BT2824^{GH16}$, which also possesses an N-terminal module of uncharacterised function (DUF4971; Supplementary Fig. 5). Signal peptide predictions using SignalP 5.0 revealed that most of the *Bacteroides* spp. GH16 family members were predicted to have a Type II signal peptide (SP), suggesting surface localisation, except for the closely related BF4060$^{GH16}$ and Baccac_02680$^{GH16}$, which had Type I SP predictions and are thus likely periplasmic. All three *A. muciniphila* GH16 enzymes are predicted to have a Type I SP, suggesting they are periplasmic, although intriguingly Amuc_2108$^{GH16}$ also has a C-terminal hydrophobic region followed by a highly positively charged sequence after this, indicative of a potential TM anchor. Notably, a previous proteomics study has shown that Amuc_2108$^{GH16}$ is present in the outer membrane in cells grown on mucin, supporting the TM anchor prediction[20]. It is possible that SignalP 5.0 is not a very accurate tool for *A. muciniphila* proteins yet (Supplementary Table 2).

**Growth of human gut bacteria on O-glycan substrates**

Human gut *Bacteroides* spp. and *A. muciniphila* were tested for their ability to grow on mucin as the sole carbon source (Supplementary Fig. 21). Of the 8 *Bacteroides* spp. tested, all could grow to some extent on PGM types II and III. *A. muciniphila* consistently grew to a higher OD than the *Bacteroides* spp. indicating that the former can access a greater proportion of the substrate. Growth of the *Bacteroides* spp. on PGM type III was commonly biphasic, indicating that the species utilises more than one substrate during a single growth curve in order of preference. Mucin preparations are likely contaminated with other host polysaccharides from glycocalyx sources that are difficult to completely remove, including chondroitin sulfate (CS), heparan sulfate (HS) and hyaluronic acid (HA). Incubation of the polysaccharide lyases from the *B. thetaiotaomicron* PULs known to degrade these host glycans [1,2] with both types of PGM show these polysaccharides are present (Supplementary Fig. 20). However, growth on PGM with a *B. thetaiotaomicron* strain where the PULs required for the degradation of these other host glycans have been knocked out shows little difference relative to the wild-type strain. There is slightly less initial growth, but the biphasic trend remains, suggesting the mucin is responsible for this. Furthermore, some species show very little growth on CS, HS or HA, but do grow on PGM (e.g. *B. fragilis* and *B. vulgatus*). Showing priority for utilisation of other host glycans over O-glycans has been observed previously in *B. thetaiotaomicron*[3].

**Phylogenetic analysis**

Phylogenetic analysis was carried out to explore the relationship between these nine mucin upregulated GH16 family members and those previously characterised (Supplementary Fig. 6). The data show that the newly identified mucin upregulated GH16 family members stem out of β-glucanase enzymes and not with any of the GH16 family members with activities on β-galactans, xyloglucan or chitinβ1,6gluconatransferases. There was significant clustering of the GH16 family members active on O-glycans.

Further phylogenetic analysis was carried out with GH16 family sequences from a number of *Bacteroides* spp. alongside the O-glycanase GH16 enzymes described in this report (Supplementary Fig. 7). In this tree, the O-glycanase enzymes cluster away from GH16 family characterised to have agarase, porphyranase, glucanase and glucosidase activities, however, they do not all cluster together. This likely represents an example of convergent evolution in the O-glycanase activity evolving multiple times from different sources. From the tree, however, the enzymes used by other species to degrade O-glycans can start to be predicted. Notably, Amuc_0875$^{GH16}$ clusters closely with GH16 sequences characterised as endo-β1,3-galactosidases shown to degrade arabinogalactan[21] and also has comparable activity to one of these enzymes on β1,3-galactan (Supplementary Fig. 16m). This raises the question of whether Amuc_0875$^{GH16}$ is a true O-glycanase, as is has relatively poor activity, however, the gene is consistently upregulated in the numerous data sets available and *A. muciniphila* also cannot grow on arabinogalactan[17]. We consider this activity to be coincidental and it much more likely that the true target for this enzyme may be a type of O-glycan structure not able to be tested in the set of assays reported here.

Interestingly, the genomic contexts of these genes are highly variable in terms of the PUL structures and adjacent genes. The GH16 family members that might be predicted to be O-glycanases are often just with a SusCD pair, with proteases or orphan genes (Supplementary Fig. 5). Furthermore, there are other GH16 enzymes in *B. thetaiotaomicron*, *B. fragilis* and *B. caccae* that have quite high sequence homology to the O-glycanases that are not upregulated with growth on O-glycans. This could be due to them being obsolete or alternatively they could have a different substrate. GH16 family members from pathogenic bacteria also from the Bacteroidetes phylum were included also in the analysis and they cluster quite well with the O-glycan active GH16 enzymes found in mutualists. A GH16 from *Tannerella forsythia* clusters well with BACCAC_03717$^{GH16}$, for example (Supplementary Fig. 7).

**Assays against classic GH16 substrates**

Recombinant forms of these GH16 family members were tested for activity against a range of β-glucan and β-galactan polysaccharides that have previously been shown to be substrates for GH16 family members. The data revealed little activity for most of the enzymes, except Amuc_0724$^{GH16}$, which displayed endo-like activity against marine laminarin and weak activity against barley β-glucan and lichenan (Supplementary Fig. 16).

The reason for this is discussed in the Main and Supplementary Discussions on the structures of these enzymes. BF4060$^{GH16}$, Baccac_02680$^{GH16}$ and Baccac_03717$^{GH16}$ also displayed some very weak activity against laminarin, but not any of the other plant polysaccharides tested. We also tested other host polysaccharides that are likely usually present in the gut in the mucus layer, including CS, HS and HA. No products were observed for CS and HA by TLC, but for HS there was a single product observed for BT2824$^{GH16}$, BF4060$^{GH16}$, Baccac_03717$^{GH16}$, Amuc_0724$^{GH16}$ and Amuc_2108$^{GH16}$, and two different products for Amuc_0875$^{GH16}$. The linker between heparan sulfate polysaccharides and protein is GlcAβ1,3Galβ1,3Galβ1,4Xyl, which has a potential GH16 cut site. We explored the potential for the O-glycan active enzymes to cleave this linker, but no activity was observed against human syndecan (Supplementary Fig. 16q).

**Substrate specificity of the GH16 family members**

A variety of defined O-glycan and human milk-derived oligosaccharides were used to assess the specificity of the GH16 substrates further (Supplementary Figs. 12-15 and Supplementary Table 4). These data indicate that the enzymes are endo β1,4-galactosidases with a requirement for a β1,3-linked sugar at the -2 position. Notably, none of the GH16 enzymes require sulfation or fucosylation decorations for activity. The Amuc_0875$^{GH16}$ enzyme could not completely degrade TriLacNAc overnight. Most of the GH16 enzymes hydrolysed the TriLacNAc rapidly under the conditions tested (Supplementary Fig. 13), however, BF4058$^{GH16}$ and Baccac_02679$^{GH16}$ displayed significantly lower activity against the hexasaccharide and degradation of triLacNAc by Amuc_0875$^{GH16}$ was only detectable after overnight incubation.

All enzymes, bar Amuc_0875 $^{GH16}$, could also degrade Lacto-N-neotetraose (LNnT) and Lacto-N-tetraose (LNT) overnight, suggesting plasticity in the positive subsites of these GH16 family members (Supplementary Fig. 14). However, substrate depletion assays indicate differences between the GH16 enzymes in terms of the importance of these subsites for activity (Supplementary Fig. 13). The activity against LNnT and LNT is comparable to TriLacNAc in the case of Baccac_03717, for example, but is much lower for BF4060$^{GH16}$. BT2824 $^{GH16}$, Baccac_02680$^{GH16}$, Amuc_0724 $^{GH16}$ and Amuc_2108 $^{GH16}$, but not dramatic. Furthermore, three of the enzymes showed preferences between the two milk oligosaccharides. Baccac_02680$^{GH16}$ degrades LNT preferentially, whereas BT2824 and Amuc0724 prefer LNnT. This may indicate a difference in preference for a β1,3 and a β1,4 linkages between the -2 and -3 subsites. Interestingly, when these GH16 family members were tested against a form of LNT where the GlcNAc is replaced with a GalNAc no activity was detected apart from trace activity for Amuc_0875 $^{GH16}$. The difference between these sugars is the position of the hydroxyl at C4 being equatorial and axial in GlcNAc and GalNAc, respectively. An axial bond at this position means the β1,4-linked Gal at this position would be at an angle relative to the rest of the oligosaccharide and no longer forming a linear chain. The lack of activity suggests that the axial hydroxyl in a GalNAc would generate a significant steric clash with the protein enough for it not to be accommodated. Sensitivity to the sugar in the -2 subsite has been documented previously[22,23]. There is also evidence to suggest that a sugar in the -2 subsite is required for activity for most of the GH16 enzymes tested here. For example, the Galβ1,4GlcNAcβ1,4Gal product that remains after TriLacNAc and LNnT degradation is not further degraded, although BT2824$^{GH16}$ and Baccac_03717$^{GH16}$ do appear to have trace activity against this (Supplementary Fig. 14).

Activity against blood group hexasaccharides type II was then used to probe the flexibility the enzymes have in the negative subsites (Supplementary Fig. 13 & 14). These glycans have a LNnT core with an α1,2-fucose and a α1,3-GalNAc or galactose (blood group A and B, respectively) on the galactose at the non-reducing end. Again different enzymes showed different preferences, but the activity of most of the enzymes were affected by these decorations relative to LNnT, with Amuc_0724$^{GH16}$ being the exception. Only BT2824$^{GH16}$ seemed to show a preference for the hydrolysis of blood group B over A. Blood group H was also tested to look at the effect of the α1,3-GalNAc or galactose on activity. BF4060$^{GH16}$, Baccac_02680$^{GH16}$, Baccac_03717$^{GH16}$ and Amuc0725 showed an increase in activity with blood group H relative to A and B, suggesting the α1,3 decorations are not well accommodated by these enzymes. In contrast, BT2824$^{GH16}$, BF4058$^{GH16}$ and Amuc_2108$^{GH16}$ activities were unaffected with the removal of the α1,3-GalNAc or galactose, which suggests that the α1,2-fucose is predominantly causing the reduced activity relative to LNnT. During prolonged incubation most of the enzymes had some activity against the blood group milk oligosaccharides and, furthermore, the products from porcine small intestinal mucin included those where the α1,3-GalNAc would be in the -4 position plus a variety of fucose and sulfate decoration (Fig. 2b, *glycans 9, 11, 13* and *16*).

All nine GH16 family members were tested against a series of disaccharides, but no activity could be observed (Supplementary Fig. 15). This is to be expected with endo-acting enzymes, as more than two subsites to be occupied to provide the binding energy for catalysis. Activity was possible for most enzymes against Lacto-N-triose, reiterating the requirement for a sugar in the -2 position (Supplementary Fig. 15g). A number substrates were tested that had an α-linked sugar at this position, but no activity was seen

(Supplementary Fig. 15). This is most likely due to the α-linkage causing a more kinked chain that cannot be accommodated by the enzyme similar to when a GalNAc is in the -2.

**Reports of O-glycan degradation by members of the GH16 family**

Although most of the GH16 family members have been characterised to have activity against terrestrial or marine plant polysaccharides, there have been two previous reports of GH16 enzymes with activity against host glycans. One of these was the activity of a GH16 family member from *Sphingobacterium multivorum* against keratan sulfate[24,25]. This is usually an environmental bacterial species, but can cause human infections, usually in immunocompromised patients[26]. Phylogenetic analysis of this GH16 shows the *S. multivorum* sequence clusters with the *Bacteroides* spp. sequences investigated in this study (Supplemental Fig. 6). This enzyme was determined to hydrolyse the Galβ1,4GlcNAc bond to release predominantly 6-O-sulfo-GlcNAcβ1,3Gal and GlcNAcβ1,3Gal products from a variety of keratan sulfate sources. Interestingly, the *S. multivorum* enzyme was found to also cleave milk oligosaccharides capped with sialic acid, which is different from what we observe in the O-glycan GH16 enzymes characterised here.

The second example of a GH16 with activity against O-glycans is from *Clostridium perfringens* and capable of removing the GlcNAcα1,4Gal disaccharide at the non-reducing end of the O-glycan[27,28], which is an epitope associated only with stomach mucin[29,30]. The crystal structure revealed a unique binding site to other GH16 family members with a pocket in the negative subsite area tailored to fit the disaccharide substrate[31] (Supplementary Fig. 18b).

**Keratanases**

Keratan sulfate chains are anchored to the protein through N-linkages, O-linkages and O-mannosylation are termed KS-I, KS-II and KS-III, respectively[32](Supplementary Fig. 4). Examples of areas of the body enriched in KS-I, KS-II and KS-III include the cornea, skeletal and brain, respectively. There are three classes of enzymes that have endo-activity against keratan - the β-galactosidases described above and keratanases type I and II. Type I enzymes are also β-galactosidases, but require sulfation for activity and cannot hydrolyse unsulfated glycans[25]. Type II enzymes are endo-β-N-acetylglucosaminidases, so hydrolyse a different linkage, and releases disaccharides and tetrasacchairdes with varying amounts of sulfation and also tolerate fucose[33,34]. There have been other reports of endo-β-galactosidases active on keratan sulfate and milk oligosaccharides from *Citrobacter freundii*, *Coccobacillus spp*. and *B. fragilis*, but insufficient information in the literature prevents us from confirming if these are from the GH16 family[25,35-37].

The GH16 enzymes reported here produced products from egg and bovine corneal keratan sulfate visible by TLC and were analysed in the same way as O-glycan products with procainamide labelling and LC-FLD-ESI-MS (Supplementary Fig. 11). The results indicate some sulfate groups can be tolerated by the GH16 family members, but heavily sulfated O-glycan fragments could not be degraded further (for example, *glycans 13-15*). This activity is distinct from Keratanase type I and II activities.

**Further discussion on the crystal structures of the O-glycan active GH16 family members**

*The characteristics of the ligand in complexed O-glycan active GH16 structures*

The non-reducing end Gal and GlcNAc modelled in the most stable $^4C_1$ chair conformation, whereas the reducing end Gal, occupying the -1 subsite, presented a $^1S_3$ skew boat conformation. The occurrence of a less stable conformation at this position has been described previously as a structural adaptation during hydrolysis in a GH16 1,3-1,4-β-glucanase[38]. The resolution of the ligand in the BF4060$^{GH16}$ structure was insufficient to define the conformation of the monosaccharides, so the ligand was modelled based on the trisaccharide from the Baccac_02680$^{GH16E143Q}$ structure.

*Accommodation of a glucose or galactose at the -1 subsite*

At the -1 subsite, the selection of glucose or galactose is very significant as the C4 points in towards the binding cleft. Analysis of the different GH16 structures available, however, reveals no strict rule for how space is created for the equatorial glucose hydroxyl or tightened up for the axial galactose hydroxyl. However, the residues on finger 1 always play a key role in controlling this space. In the *Bacteroides* structures described in this study, a tryptophan from finger 1 blocks off the possibility of an equatorial positioning in this area to produce a galactose-tailored pocket (Fig. 3b and Supplementary Fig. 19a). In contrast, a surface representation of the Amuc_0724$^{GH16}$ structure shows a relatively open space around the C4 and C6 formed by finger 1 being further away, with an arginine replacing the tryptophan and relatively small residues lining the β-strands in this area (Supplementary Fig. 19a). This is a possible structural explanation for the activity of this enzyme towards glucose polymers.

A closer look at this region in Amuc_0724$^{GH16}$ structure, shows that this open space is actually a short tunnel and possibly accommodates decoration of the galactose (Fig. 2c). Pockets and tunnels above the -1 subsite have previously been observed in a number of

GH16 structures, for example, the ZgLamC$_{GH16-E142S}$ structure from *Zobellia galactanivorans* (Supplementary Fig. 19b). This enzyme is active on laminarin (predominantly β1,3-glucan with occasional β1,6-branching), but also mixed-linked glucan. It was crystallised with a glycerol inside the -1 subsite pocket and it was suggested that it may be able to accommodate a branching β1,6 monosaccharide[39]. To test this idea out for the O-glycan active GH16 family members, especially Amuc_0724$^{GH16}$, we took lacto-N-hexaose and enzymatically removed the capping galactose to leave a lactose with two GlcNAc linked to the galactose through β1,3 and β1,6 bonds. Most of the enzymes were active on the same structure with just the β1,3-linked GlcNAc (Supplementary Fig. 15g), however, no significant activity could be seen against the branched substrate. This pocket, therefore, most likely cannot accommodate a β1,6-GlcNAc branch.

GH16 family members that have to accommodate galactose at this position include the agarases, porphyranases and pectic galactanases. In the agarase structures currently available that are complexed with substrate or product, a glutamic acid from the β-sheet of the cleft coordinates with and fills the space around the C4 hydroxyl. A proline from finger 1 sits adjacent to this (Supplementary Fig. 18c)[40]. In the porphyranase structures, there is a glutamic acid in approximately the same position, but an arginine from finger 1 also coordinates with C4 and fills this space (Supplementary Fig. 18c)[22]. From these observations about the -1 subsite in all GH16 structures, we can conclude that selection of glucose of galactose at the -1 position is achieved in multiple ways by GH16 family members. There are currently no pectic β1,4-galactanase structures to allow comparison.

*Specificity for a 1,3-linkage between the -1 and -2 sugars*

The linkage between the monosaccharides in the -1 and -2 subsites is a 1,3 for some GH16 family members, but this is not the case for xyloglucan hydrolases, xyloglucan endo transferases, pectic galactosidases and chitinβ1,6gluconatransferases. This specificity is commonly selected for in β-glucanases through the geometry of the two hydrophobic platforms that the monosaccharides at these positions sit on, which are at 90 ° relative to each other[41]. In the structures available for β-glucan-active enzymes, where a glucose would be in the -2 position, an aromatic residue from finger 3 or the local β-strand acts as a platform for the α-face of this sugar to stack parallel to. The endo-mucinase GH16 structures presented in this study also have this aromatic platform, so are most comparable to glucanases in this respect (Fig. 3b). This structural observation corresponds with the biochemical characterisation of the O-glycan active GH16 family members described above as requiring a β1,3-linked GlcNAc in the -2 subsite for activity.

Interactions with the β-face of this sugar is usually through residues coming from finger 1 and from the β-sheet in the cleft, but these are variable even within enzymes with the same activities. The structures available for the enzymes degrading marine polysaccharides have to deal with an alpha linkage at the -2 position, an anhydrogalactose in the case of agarases and carrageenanases, and sulfation at C6 for porphyranases. This does not involve aromatic stacking but coordination through polar interactions and non-parallel π-stacking with aromatic residues (Supplementary Fig. 18c).

The N-acetyl coming off the C2 of the GlcNAc for the O-glycanases points out into solution away from the cleft, so accommodating this in a specific cleft pocket is not an issue for the enzymes reported here. The cleft of the GH16 does have to be fairly open to allow the N-acetyl and would be too bulky for GH16 enzymes with more closed clefts, so this is likely one criterion for O-glycan active GH16 enzymes. The C6 of the GlcNAc in the -2 subsite of the O-glycanases is bonded to a methionine in Baccac_02680$^{GH16}$ and BF4060 $^{GH16}$ and likely a leucine in Amuc0724$^{GH16}$ and Baccac_03717$^{GH16}$ (Fig. 3b and Supplementary Fig. 19a).

In addition to β-glucanases, those GH16 family members accommodating glucose-type sugars at the -1 include the xyloglucan hydrolases, xyloglucan endo transferases and chitinβ1,6gluconatransferases. These are repeating β1,4-linkages, so require an open structure around the C4 allowed by the absence of a Finger 1. These enzymes also therefore do not have the characteristic of a 1,3 linkage between the -1 and -2 subsites. Structures of the xyloglucan active enzymes show a significantly different cleft to accommodate the β1,4-linkages (Supplementary Fig. 18a). There are currently no structures available of chitinβ1,6gluconatransferases for comparison.

*The -3 subsite and beyond*

The area around the -3 subsite galactose is fairly open in the Baccac_02680$^{GH16E143Q}$ and BF4060$^{GH16}$ structures and the same is true when the product is overlaid into Baccac_03717$^{GH16}$. The extended Finger 2 in the Amuc0724$^{GH16}$ structure, however, could theoretically interact with a longer substrate. An extensive loop structure reaching down to the binding cleft has been seen before in other GH16 structures, such as the κ-Carrageenases from *Zobellia galactanivorans* and *Pseudoalteromonas carrageenovora* (Supplementary Fig. 18d), which possess a similar Finger 2 and it interacts with the -4 sugar in the case of the *P. carrageenovora* structure[42].

In the O-glycan structures presented here, there is space at the -3 for a β1,4-linked galactose, but it is easy to see that if the sugar in the -3 did not continue the chain in a linear fashion then that may not be accommodated. We were able to see this when we tested for activity of the nine O-glycan active GH16 enzymes against Galβ1,4GalNAcβ1,3Galβ1,4Glc and saw negative results, except in the case of Amuc_0875$^{GH16}$ that had trace activity. The axial position of the hydroxyl at C4 in a GalNAc would mean the galactose in the -3 would now be kinked, therefore showing specificity for a GlcNAc at the -2 position. Agarases, carrageenases and porphyranases (galactans) have fairly linear glycans due to alpha bonds when a 1,3-linkage is used, which alternates with the β1,4 linkage (Fig. 3e).

*The positive subsites*

The positive subsites of retaining GH enzyme structures are rarely occupied with product due to the low binding affinity of the positive subsites to facilitate rapid leaving of the product after the initial glycosylation step[43,44]. However, structures of a GH16 from *Phanerochaete chrysosporium* with different glucan products bound allow us to hypothesise about how the positive subsite sugars would be accommodated in the O-glycan-active GH16 enzymes. Orientation of the glucose in the +1 subsite of three structures (PDB codes 2W52, 2WLQ and 2WN) has the β-face fronting a tryptophan from finger 6. The O-glycan active enzymes have an aromatic that overlays well with this tryptophan. Analysis of the positive subsites of GH16 structures show that the majority of them have an aromatic in this area and it could straddle both the +1 and +2 subsites or just one depending on the structure in question. Aromatic residues are sometimes not present in the +1 subsite of GH16 enzymes active on marine polysaccharides. The position of the +1 sugar in the *P. chrysosporium* GH16 suggests the hydrolysis of a β1,3 bond. For the O-glycan active GH16 enzymes, however, this would be a β1,4 bond and rotation of the sugar to reflect this would place the C6 pointing into the cleft of the enzyme. Overlay of the +1 glucose from 2WLQ indicate that this would be spatially difficult to accommodate, but it is possible the sugar sits slightly further away from the cleft than seen in glucanases (Fig. 3d).

The importance of the positive subsite residues was analysed by comparing the rates against triLacNAc versus milk oligosaccharides LNnT and LNT (Supplementary Fig. 13). The activity of BF4060$^{GH16}$ on the milk oligosaccharides was significantly decreased compared to triLacNAc, but this was not the case for Baccac_03717$^{GH16}$. The structures of Baccac_03717$^{GH16}$ is much more open at this position, whereas BF4060$^{GH16}$ has a much more closed slot for the +1 sugar to sit in and a serine (S174) coming from Finger 5 could potentially interact with the N-acetyl and pincher it against Finger 6 (Fig. 3d). Therefore, BF4060$^{GH16}$ could be showing specificity for GlcNAc in the +1, but it is also possible that the enzymes are sensitive to the number of positive subsites that are filled.

For the Amuc_0724$^{GH16}$ structure, the tryptophan at this position (W279) had dual occupancy so could be modelled to sit in the cleft or flipped out of this position away from the cleft. The flipped out position of the tyrosines from the two molecules in the unit cell may be an artefact of crystallographic dimer rather than biologically relevant (Supplementary Fig. 19c). A dynamic tyrosine has also been seen in the positive subsites of a GH16 from *B. ovatus* with β-glucanase activity[45] (Supplementary Fig. 17d).

*Accommodating sulfate, fucose and blood group decorations along the polyLacNAc chain by O-glycan active GH16 family members*

O-glycans can have 3S (capping) and 6S sulfation on galactose and 6S also on GlcNAc. Overlay of the product from the 5OCQ κ-carrageenases from *Pseudoalteromonas carrageenovora* with the Amuc_0725$^{GH16}$ structure shows how the tunnel structure in the -1 subsite could accommodate a sulfate group on the Gal at 6S, however this is most likely not a possibility for the *Bacteroides* spp. structures (Supplementary Fig. 19e)[42]. Overlay of 3ILF porphyranase from *Zobellia galactanivorans* with the O-glycan active GH16 structures mimics a 6S sulfation of the GlcNAc in the -2 subsite and all enzymes show potential in accommodating this decoration (Supplementary Fig. 19f)[22]. At the -3 subsite, C6 of the galactose points into solution, so could possibly accommodate sulfate here also.

Fucose can be linked through α1,2/3/4 bonds in O-glycans. It is most likely not possible for a Gal to be in the -1 position if there is a fucose attached. However, C3 of the GlcNAc in the -2 position looks open to fucose decoration at this position as the clefts have a fairly open structure. We characterised if the nine O-glycan active enzymes could accommodate blood group sugars appended to the -3 Gal. The α1,2-linked fucose would point into the cleft and the α1,3-linked GalNAc or Gal would be at approximately 90° to the rest of the linear substrate. As a general trend, the blood group sugars reduce activity of the O-glycan active GH16 enzymes and the removal of the α1,3-linked GalNAc or Gal attenuates this effect (Supplementary Fig. 13). There does look like there are pockets for a α1,2-linked fucose in the -3' position in the crystal structures presented in this study.

*Minimum substrate requirements*

For all the structures, there are three well-defined subsites from -2 to +1 with some possible interactions at the -3 and +2 positions. In contrast, it is not unusual for other GH16 family members to have longer binding clefts, which generates a more specific substrate specificity and requirement for the subsites to be filled for catalytic activity, for example AgaD, which requires a minimum product size of DP8[46]. The relatively small number of defined subsites in the O-glycan active GH16 family members described in this study and their minimum substrate length being DP3 likely reflects the variable nature of O-glycan as a substrate, particularly in terms of the sulfate and fucose decoration.

**Activities of the two other CAZymes from the *B. fragilis* PUL**

There are two genes predicted to be from other CAZy families in the *B. fragilis* PUL where the O-glycan active GH16 genes are encoded. They are from families GH20 and GH35 and both predicted to have SPI type signal peptides (Supplementary Table 2). Recombinant versions of these enzymes were expressed as described for the GH16 enzymes in this report and tested against a variety of defined oligosaccharides to determine specificity (Supplementary Fig. 10 and 22).

BF4061$^{GH35}$ was found to be active against LacNAc, Lacto-N-biose and Galβ1,3Glu, partially active against lactose and inactive against Galβ1,4Gal. This indicates a preference for GlcNAc>Glc>Gal in the +1 position, thus a specificity for mucin-type oligosaccharides over milk and plant-type saccarides (Supplementary Fig 10). There was no activity found against fucosylated versions of these oligosaccharides, indicating that fucose needs to be removed prior to this enzyme carrying out its function. This has been seen previously for host glycans, where fucose needed to be removed from the antenna of complex N-glycans from human IgA colostrum before the galactosidase (BT0461$^{GH2}$) could act[47]. This galactosidase comes from a different CAZy family, which illustrates how different bacteria evolve different ways of dealing with similar substrates. Interestingly, BF4061$^{GH35}$ described here can accommodate both β1,3 and β1,4-linkages, whereas BT0461$^{GH2}$ can only accommodate β1,4-linkages and these specificities represent the linkages present in their respective substrates. BF4061$^{GH35}$ can also remove β1,3-galactose when there is a GalNAc in the +1 subsite and both capping galactose from lacto-N-hexaose, where these substrates represent O-glycan core and branched structures, respectively (Supplementary Fig. 10).

The second CAZyme, BF4059$^{GH20}$, was very broad acting in accommodating a variety of linkages and monosaccharides in the +1 position (Supplementary Fig 22). Chitooligosacchrides of a degree of polymerisation between 2 and 5 could be degraded to GlcNAc and non-reducing end GlcNAc could be removed from O-glycan-type structures (GlcNAc β1,3Gal, Lacto-N-triose and Lacto-N-hexaose). BF4059$^{GH20}$ can also degrade GalNAcβ1,3Gal, but not GalNAcβ1,3Galβ1,4Glu. N-glycan type structure GlcNAcβ1,2Man could also be degraded, reflecting the broad activity of this enzyme.

The predicted localisation of these enzymes and their specificities most likely places them in the periplasm of *B. fragilis* acting on the GH16 products that have been imported into the cell. The action of these enzymes plus fucosidases and sulfatases would allow degradation of the GH16 products down to monosaccharides.

1 Cartmell, A. *et al.* How members of the human gut microbiota overcome the sulfation problem posed by glycosaminoglycans. *Proceedings of the National Academy of Sciences of the United States of America* **114**, 7037-7042, doi:10.1073/pnas.1704367114 (2017).

2 Ndeh, D. *et al.* The human gut microbe Bacteroides thetaiotaomicron encodes the founding member of a novel glycosaminoglycan-degrading polysaccharide lyase family PL29. *The Journal of biological chemistry* **293**, 17906-17916, doi:10.1074/jbc.RA118.004510 (2018).

3 Martens, E. C., Chiang, H. C. & Gordon, J. I. Mucosal glycan foraging enhances fitness and transmission of a saccharolytic human gut bacterial symbiont. *Cell host & microbe* **4**, 447-457, doi:10.1016/j.chom.2008.09.007 (2008).

4 Johansson, M. E. *et al.* The inner of the two Muc2 mucin-dependent mucus layers in colon is devoid of bacteria. *Proceedings of the National Academy of Sciences of the United States of America* **105**, 15064-15069, doi:10.1073/pnas.0803124105 (2008).

5 Ambort, D. *et al.* Calcium and pH-dependent packing and release of the gel-forming MUC2 mucin. *Proceedings of the National Academy of Sciences of the United States of America* **109**, 5645-5650, doi:10.1073/pnas.1120269109 (2012).

6 Johansson, M. E. Fast renewal of the distal colonic mucus layers by the surface goblet cells as measured by in vivo labeling of mucin glycoproteins. *PloS one* **7**, e41009, doi:10.1371/journal.pone.0041009 (2012).

7 Johansson, M. E. *et al.* Normalization of Host Intestinal Mucus Layers Requires Long-Term Microbial Colonization. *Cell host & microbe* **18**, 582-592, doi:10.1016/j.chom.2015.10.007 (2015).

8 Hansson, G. C. Mucus and mucins in diseases of the intestinal and respiratory tracts. *J Intern Med*, doi:10.1111/joim.12910 (2019).

9 Larsson, J. M., Karlsson, H., Sjovall, H. & Hansson, G. C. A complex, but uniform O-glycosylation of the human MUC2 mucin from colonic biopsies analyzed by nanoLC/MSn. *Glycobiology* **19**, 756-766, doi:10.1093/glycob/cwp048 (2009).

10 Andersch-Bjorkman, Y., Thomsson, K. A., Holmen Larsson, J. M., Ekerhovd, E. & Hansson, G. C. Large scale identification of proteins, mucins, and their O-glycosylation in the endocervical mucus during the menstrual cycle. *Molecular & cellular proteomics : MCP* **6**, 708-716, doi:10.1074/mcp.M600439-MCP200 (2007).

11 Thomsson, K. A., Schulz, B. L., Packer, N. H. & Karlsson, N. G. MUC5B glycosylation in human saliva reflects blood group and secretor status. *Glycobiology* **15**, 791-804, doi:10.1093/glycob/cwi059 (2005).

12 Schulz, B. L. *et al.* Mucin glycosylation changes in cystic fibrosis lung disease are not manifest in submucosal gland secretions. *The Biochemical journal* **387**, 911-919, doi:10.1042/bj20041641 (2005).

13 Fujita, M. *et al.* Glycoside hydrolase family 89 alpha-N-acetylglucosaminidase from Clostridium perfringens specifically acts on GlcNAc alpha1,4Gal beta1R at the non-reducing terminus of O-glycans in gastric mucin. *The Journal of biological chemistry* **286**, 6479-6489, doi:10.1074/jbc.M110.206722 (2011).

14 Nordman, H. *et al.* Gastric MUC5AC and MUC6 are large oligomeric mucins that differ in size, glycosylation and tissue distribution. *The Biochemical journal* **364**, 191-200, doi:10.1042/bj3640191 (2002).

15 Jin, C. *et al.* Structural Diversity of Human Gastric Mucin Glycans. *Molecular & cellular proteomics : MCP* **16**, 743-758, doi:10.1074/mcp.M116.067983 (2017).

16 Pudlo, N. A. *et al.* Symbiotic Human Gut Bacteria with Variable Metabolic Priorities for Host Mucosal Glycans. *MBio* **6**, e01282-01215, doi:10.1128/mBio.01282-15 (2015).

17 Desai, M. S. *et al.* A Dietary Fiber-Deprived Gut Microbiota Degrades the Colonic Mucus Barrier and Enhances Pathogen Susceptibility. *Cell* **167**, 1339-1353.e1321, doi:10.1016/j.cell.2016.10.043 (2016).

18 Ottman, N. *et al.* Genome-Scale Model and Omics Analysis of Metabolic Capacities of Akkermansia muciniphila Reveal a Preferential Mucin-Degrading Lifestyle. *Applied and environmental microbiology* **83**, doi:10.1128/aem.01014-17 (2017).

19 Shin, J. *et al.* Elucidation of Akkermansia muciniphila Probiotic Traits Driven by Mucin Depletion. *Frontiers in microbiology* **10**, 1137, doi:10.3389/fmicb.2019.01137 (2019).

20 Ottman, N. *et al.* Characterization of Outer Membrane Proteome of Akkermansia muciniphila Reveals Sets of Novel Proteins Exposed to the Human Intestine. *Frontiers in microbiology* **7**, 1157, doi:10.3389/fmicb.2016.01157 (2016).

21 Cartmell, A. *et al.* A surface endogalactanase in Bacteroides thetaiotaomicron confers keystone status for arabinogalactan degradation. *Nature microbiology* **3**, 1314-1326, doi:10.1038/s41564-018-0258-8 (2018).

22 Hehemann, J. H. *et al.* Transfer of carbohydrate-active enzymes from marine bacteria to Japanese gut microbiota. *Nature* **464**, 908-912, doi:10.1038/nature08937 (2010).

23 Hehemann, J. H., Kelly, A. G., Pudlo, N. A., Martens, E. C. & Boraston, A. B. Bacteria of the human gut microbiome catabolize red seaweed glycans with carbohydrate-active enzyme updates from extrinsic microbes. *Proceedings of the National Academy of Sciences of the United States of America* **109**, 19786-19791, doi:10.1073/pnas.1211002109 (2012).

24 Kitamikado, M., Ito, M. & Li, Y. T. Isolation and characterization of a keratan sulfate-degrading endo-beta-galactosidase from Flavobacterium keratolyticus. *The Journal of biological chemistry* **256**, 3906-3909 (1981).

25 Ito, M., Hirabayashi, Y. & Yamagata, T. Substrate specificity of endo-beta-galactosidases from Flavobacterium keratolyticus and Escherichia freundii is different from that of Pseudomonas sp. *Journal of biochemistry* **100**, 773-780, doi:10.1093/oxfordjournals.jbchem.a121770 (1986).

26 Abro, A. H., Rahimi Shahmirzadi, M. R., Jasim, L. M., Badreddine, S. & Al Deesi, Z. Sphingobacterium multivorum Bacteremia and Acute Meningitis in an Immunocompetent Adult Patient: A Case Report. *Iranian Red Crescent medical journal* **18**, e38750, doi:10.5812/ircmj.38750 (2016).

27 Ashida, H. *et al.* A novel endo-beta-galactosidase from Clostridium perfringens that liberates the disaccharide GlcNAcalpha 1-->Gal from glycans specifically expressed in the gastric gland mucous cell-type mucin. *The Journal of biological chemistry* **276**, 28226-28232, doi:10.1074/jbc.M103589200 (2001).

28 Ashida, H., Maskos, K., Li, S. C. & Li, Y. T. Characterization of a novel endo-beta-galactosidase specific for releasing the disaccharide GlcNAc alpha 1-->4Gal from glycoconjugates. *Biochemistry* **41**, 2388-2395, doi:10.1021/bi011940e (2002).

29 Nakayama, J. *et al.* Expression cloning of a human alpha1, 4-N-acetylglucosaminyltransferase that forms GlcNAcalpha1-->4Galbeta-->R, a glycan specifically expressed in the gastric gland mucous cell-type mucin. *Proceedings of the National Academy of Sciences of the United States of America* **96**, 8991-8996, doi:10.1073/pnas.96.16.8991 (1999).

30 Ishihara, K. *et al.* Peripheral alpha-linked N-acetylglucosamine on the carbohydrate moiety of mucin derived from mammalian gastric gland mucous cells: epitope recognized by a newly characterized monoclonal antibody. *The Biochemical journal* **318 ( Pt 2)**, 409-416, doi:10.1042/bj3180409 (1996).

31 Tempel, W. *et al.* Three-dimensional structure of GlcNAcalpha1-4Gal releasing endo-beta-galactosidase from Clostridium perfringens. *Proteins* **59**, 141-144, doi:10.1002/prot.20363 (2005).

32 Caterson, B. & Melrose, J. Keratan sulfate, a complex glycosaminoglycan with unique functional capability. *Glycobiology* **28**, 182-206, doi:10.1093/glycob/cwy003 (2018).

33 Yamagishi, K. *et al.* Purification, characterization, and molecular cloning of a novel keratan sulfate hydrolase, endo-beta-N-acetylglucosaminidase, from Bacillus circulans. *The Journal of biological chemistry* **278**, 25766-25772, doi:10.1074/jbc.M212183200 (2003).

34 Wang, H. *et al.* Construction and functional characterization of truncated versions of recombinant keratanase II from Bacillus circulans. *Glycoconjugate journal* **34**, 643-649, doi:10.1007/s10719-017-9786-3 (2017).

35 Fukuda, M. N. & Matsumura, G. Endo-beta-galactosidase of Escherichia freundii. Purification and endoglycosidic action on keratan sulfates, oligosaccharides, and blood group active glycoprotein. *The Journal of biological chemistry* **251**, 6218-6225 (1976).

36 Scudder, P., Uemura, K., Dolby, J., Fukuda, M. N. & Feizi, T. Isolation and characterization of an endo-beta-galactosidase from Bacteroides fragilis. *The Biochemical journal* **213**, 485-494, doi:10.1042/bj2130485 (1983).

37 Hirano, S. & Meyer, K. Enzymatic degradation of corneal and cartilaginous keratosulfates. *Biochemical and biophysical research communications* **44**, 1371-1375, doi:10.1016/s0006-291x(71)80237-1 (1971).

38 Biarnes, X. *et al.* The conformational free energy landscape of beta-D-glucopyranose. Implications for substrate preactivation in beta-glucoside hydrolases. *Journal of the American Chemical Society* **129**, 10686-10693, doi:10.1021/ja068411o (2007).

39 Labourel, A. *et al.* Structural and biochemical characterization of the laminarinase ZgLamCGH16 from Zobellia galactanivorans suggests preferred recognition of branched laminarin. *Acta crystallographica. Section D, Biological crystallography* **71**, 173-184, doi:10.1107/s139900471402450x (2015).

40 Allouch, J., Helbert, W., Henrissat, B. & Czjzek, M. Parallel substrate binding sites in a beta-agarase suggest a novel mode of action on double-helical agarose. *Structure (London, England : 1993)* **12**, 623-632, doi:10.1016/j.str.2004.02.020 (2004).

41 Vasur, J. *et al.* X-ray crystal structures of Phanerochaete chrysosporium Laminarinase 16A in complex with products from lichenin and laminarin hydrolysis. *The FEBS journal* **276**, 3858-3869, doi:10.1111/j.1742-4658.2009.07099.x (2009).

42 Matard-Mann, M. *et al.* Structural insights into marine carbohydrate degradation by family GH16 kappa-carrageenases. *The Journal of biological chemistry* **292**, 19919-19934, doi:10.1074/jbc.M117.808279 (2017).

43 Davies, G. J. *et al.* Snapshots along an enzymatic reaction coordinate: analysis of a retaining beta-glycoside hydrolase. *Biochemistry* **37**, 11707-11713, doi:10.1021/bi981315i (1998).

44 Davies, G. J., Tolley, S. P., Henrissat, B., Hjort, C. & Schulein, M. Structures of oligosaccharide-bound forms of the endoglucanase V from Humicola insolens at 1.9 A resolution. *Biochemistry* **34**, 16210-16220, doi:10.1021/bi00049a037 (1995).

45 Tamura, K. *et al.* Molecular Mechanism by which Prominent Human Gut Bacteroidetes Utilize Mixed-Linkage Beta-Glucans, Major Health-Promoting Cereal Polysaccharides. *Cell Rep* **21**, 417-430, doi:10.1016/j.celrep.2017.09.049 (2017).

46 Hehemann, J. H. *et al.* Biochemical and structural characterization of the complex agarolytic enzyme system from the marine bacterium Zobellia galactanivorans. *The Journal of biological chemistry* **287**, 30571-30584, doi:10.1074/jbc.M112.377184 (2012).

47 Briliute, J. *et al.* Complex N-glycan breakdown by gut Bacteroides involves an extensive enzymatic apparatus encoded by multiple co-regulated genetic loci. *Nature microbiology* **4**, 1571-1581, doi:10.1038/s41564-019-0466-x (2019).

**Supplementary Table 1│ Percentage identity between all members of the GH16 family included in this study**

| | BT2824$^{GH16}$ | BF4058$^{GH16}$ | BF4060$^{GH16}$ | Baccac_02679$^{GH16}$ | Baccac_02680$^{GH16}$ | Baccac_03717$^{GH16}$ | Amuc_0724$^{GH16}$ | Amuc_0875$^{GH16}$ | Amuc_2108$^{GH16}$ |
|---|---|---|---|---|---|---|---|---|---|
| BT2824$^{GH16}$ | 100 % | 38 % | 37 % | 38 % | 38 % | 35 % | 27 % | 25 % | 30 % |
| BF4058$^{GH16}$ | | 100 % | 28 % | 87 % | 27 % | 31 % | 29 % | 26 % | 29 % |
| BF4060$^{GH16}$ | | | 100 % | 29 % | 79 % | 31 % | 26 % | 24 % | 29 % |
| Baccac_02679$^{GH16}$ | | | | 100 % | 27 % | 31 % | 29 % | 27 % | 29 % |
| Baccac_02680$^{GH16}$ | | | | | 100 % | 31 % | 24 % | 22 % | 28 % |
| Baccac_03717$^{GH16}$ | | | | | | 100 % | 30 % | 28 % | 36 % |
| Amuc_0724$^{GH16}$ | | | | | | | 100 % | 34 % | 29 % |
| Amuc_0875$^{GH16}$ | | | | | | | | 100 % | 23 % |
| Amuc_2108$^{GH16}$ | | | | | | | | | 100 % |

**Supplementary Table 2│ Signal peptide predictions and likely cellular locations of the Cazymes characterised in this study.**

| Species | Locus Tag | Cazy Classification | Predicted signal peptide[a] | Experimentally determined location |
|---|---|---|---|---|
| *B. thetaiotaomicron* | BT2824 | GH16 | SPII | |
| *B. fragilis* | BF4058 | GH16 | SPII | |
| *B. fragilis* | BF4059 | GH20 | SPII | |
| *B. fragilis* | BF4060 | GH16 | SPI | |
| *B. fragilis* | BF4061 | GH35 | SPI | |
| *B. caccae* | Baccac_02679 | GH16 | SPII | |
| *B. caccae* | Baccac_02680 | GH16 | SPI | |
| *B. caccae* | Baccac_03717 | GH16 | SPII | |
| *A. muciniphila* | Amuc_0724 | GH16 | nd | |
| *A. muciniphila* | Amuc_0875 | GH16 | SPI | |
| *A. muciniphila* | Amuc_2108 | GH16 | SPI[b] | Outer membrane[20] |

[a]Predictions carried out using SigP5.0

[b]Has a very hydrophobic region at the C-terminus that could be a membrane anchor

**Supplementary Table 3│List of strains used in this study, the locus tag prefixes and also shortened versions used in Supplementary Figs. 5-7.**

| Species | Strain analysed | True locus tag prefix | Prefix used for brevity in Supplementary Fig. 20 |
|---|---|---|---|
| *A. muciniphila* | ATCC | Amuc | Amuc |
| *B. caecimuris* | I48 | Bcae | Bcae |
| *B. caccae* | ATCC43185 | BACCAC | BC |
| *B. cellulosilyticus* | DSM 14838 | BACCELL | BAC |
| *B. dorei* | DSM 17855 | BACDOR | BACDOR |
| *B. fragilis* | NCTC 9343 | BF | BF |
| *B. finegoldii* | DSM 17565 | BACFIN | BACFIN |
| *B. helocogenes* | DSM 20613 | Bache | Bache |
| *B. heparinolyticus* | DSM 23917 | Bhep | Bhep |
| *B. intestinalis* | 341, DSM 17393 | BACINT | BACINT |
| *B. plebius* | DSM 17135 | BACPLE | BACPLE |
| *B. thetaiotaomicron* | VPI-5482 | BT | BT |
| *B. ovatus* | ATCC 8482 | BACOVA | BO |
| *B. vulatus* | ATCC 8483 | BVU | BVU |
| *B. xylanisolvans* | XB1A | BXY | BXY |

**Supplementary Table 4│ Activity of GH16 family members against specific O-glycan substrates** This summarises the enzyme activities of the GH16 family members analysed in this study and characterises what sugars and linkages can be accommodated at different subsites. The information is derived from Supplementary Figs. 13-15. The blue text specifically refers to the results obtained from the substrate depletion data where substrate preferences were analysed in greater detail (Supplementary Fig. 13).

| | | $BT2824^{GH16}$ | $BF4058^{GH16}$ | $BF4060^{GH16}$ | $Baccac\_02679^{GH16}$ | $Baccac\_02680^{GH16}$ | $Baccac\_03717^{GH16}$ | $Amuc\_0724^{GH16}$ | $Amuc\_0875^{GH16}$ | $Amuc\_2108^{GH16}$ |
|---|---|---|---|---|---|---|---|---|---|---|
| Linear substrates | **TriLacNAc** spanning -3 to +3 subsites (Supplementary Fig Xa) | Yes | Yes | Yes | Yes | Yes | Yes | Yes | Partial | Yes |
| | **TriLacNAc product** GlcNAcβ1,3Galβ1,4GlcNAc spanning the -2 to +1 subsites (Supplementary Fig Xa) | Yes | Yes | Yes | Yes | Yes | Yes | Yes | Partial | Yes |
| | **Lacto-N-neotetraose** spanning the -3 to +1 subsites (Supplementary Fig Xb) | Yes Reduced rate relative to TriLacNAc **Preference over LNT** | Yes Very reduced rate relative to TriLacNAc | Yes Very reduced rate relative to TriLacNAc | Yes Very reduced rate relative to TriLacNAc | Yes Reduced rate relative to TriLacNAc | Yes Unaffected relative to TriLacNAc | Yes Reduced rate relative to TriLacNAc **Preference over LNT** | No | Yes Reduced rate relative to TriLacNAc |
| | **Lacto-N-neotetraose product** Galβ1,4GlcNAcβ1,3Gal spanning the -1 to +2 or -2 to +1 subsites (Supplementary Fig Xb) | Yes | No | No | No | No | Yes | Partial | No | Partial |

| | | | | | | | | | | |
|---|---|---|---|---|---|---|---|---|---|---|
| | **Lacto-N-tetraose** spanning the -3 to +1 subsites (Supplementary Fig Xc) | Yes Reduced rate relative to TriLacNAc | Yes Very reduced rate relative to TriLacNAc | Yes Very reduced rate relative to TriLacNAc | Yes Very reduced rate relative to TriLacNAc | Yes Reduced rate relative to TriLacNAc **Preference over LNnT** | Yes Unaffected relative to TriLacNAc | Yes Reduced rate relative to TriLacNAc | No | Yes Reduced rate relative to TriLacNAc |
| | **Lacto-N-neotetraose product** Galβ1,3GlcNAcβ1,3Gal spanning the -1 to +2 or -2 to +1 subsites (Supplementary Fig Xc) | No | No | Partial | No | Partial | No | Partial | No | No |
| | **Lacto-N-triose** spanning the -2 to +1 subsites (Supplementary Fig Xc) | Yes | Yes | Yes | Partial | Yes | Yes | Yes | No | Yes |
| | **LacNAc** | No | No | No | No | No | No | No | No | No |
| | **Galβ1,4Gal** | No | No | No | No | No | No | No | No | No |
| | **Lacto-N-biose** | No | No | No | No | No | No | No | No | No |
| | **Lactose** | No | No | No | No | No | No | No | No | No |
| α-linked sugar at the -2 subsite | **Blood group H trisaccharide** α1,2 fucose at the -2 subsite | No | No | No | No | No | No | No | No | No |
| | **3-Sialyllactose** α2,3 sialic acid at the -2 subsite | No | No | No | No | No | No | No | No | No |
| | **P1 antigen** α1,4 Galactose at the -2 subsite | No | No | No | No | No | No | No | No | No |
| | **Globotriose** α1,4 Galactose at the -2 subsite | No | No | No | No | No | No | No | No | No |
| | **Forssman antigen** α1,4 Galactose at the -2 subsite | No | No | No | No | No | No | No | No | No |

| | | | | | | | | | | |
|---|---|---|---|---|---|---|---|---|---|---|
| α1,4 fucose branch at the -3' subsite | **Lacto-N-fucopentaose II** spanning the -3' to +1 subsites | Yes | Yes | Yes | Partial | Yes | Yes | Partial | No | Yes |
| Blood groups α-linked sugars at the -4 and -4' subsites | **Blood Group A hexasaccharide II** | Yes Reduced rate relative to LNT and LNnT. | Yes Reduced rate relative to LNT and LNnT. | Partial Reduced rate relative to LNT and LNnT. | No | Partial | Yes Reduced rate relative to LNT and LNnT. | Yes Rate unaffected relative to LNT and LNnT | No | Partial Reduced rate relative to LNT and LNnT. |
| | **Blood Group B hexasaccharide II** | Yes Reduced rate relative to LNT and LNnT. **Preference for over A** | Yes Reduced rate relative to LNT and LNnT. | Yes | No | Partial | Yes Reduced rate relative to LNT and LNnT. | Yes Rate unaffected relative to LNT and LNnT | No | Partial Reduced rate relative to LNT and LNnT. |
| | **Blood Group H pentasaccharide II** | Yes Improved rate relative to BGA and B | Yes Same rate to BGA and B | Yes Improved rate relative to BGA and B | Partial Same rate to BGA and B | Yes Improved rate relative to BGA and B | Yes Improved rate relative to BGA and B | Yes Rate unaffected relative to all | No | Partial Rate unaffected relative to BGA and B |

**Supplementary Table 5│Data collection and refinement statistics**

| | Amuc_0724$^{GH16}$ | Baccac_02680$^{GH16}$ ligand | Baccac_03717$^{GH16}$ | BF4060$^{GH16}$ ligand |
|---|---|---|---|---|
| Date | 25/01/19 | 25/11/18 | 11/10/18 | 11/10/18 |
| Source | I24 | I03 | I03 | I03 |
| Wavelength (Å) | 0.9786 | 0.9793 | 0.9796 | 0.9796 |
| Space group | $P2_12_12_1$ | $P4_1$ | $C222_1$ | $P6_122$ |
| Cell dimensions | | | | |
| *a*, *b*, *c* (Å) | 87.5, 96.1, 128.8 | 82.92, 82.92, 121.31 | 46.9, 87.1, 156.2 | 156.6, 156.6, 197.0 |
| α, β, γ (°) | 90, 90, 90 | 90, 90, 90 | 90 ,90, 90 | 90.0, 90.0, 120.0 |
| No. of measured reflections | 214047 (21796) | 418545 (31263) | 136701 (11117) | 469646 (98047) |
| No. of independent reflections | 33993 (4096) | 55261 (4092) | 19202 (1554) | 22156 (4469) |
| Resolution (Å) | 19.90 – 2.70 (2.83 – 2.70) | 19.99 – 2.00 (2.05 – 2.00) | 43.57– 2.10 (2.16 – 2.10) | 197.05 – 3.30 ( 3.56 – 3.30) |
| $CC_{1/2}$ | 0.991 (0.485) | 0.999 (0.472) | 0.995 (0.743) | 0.992 (0.679) |
| $I/\sigma I$ | 7.7 (1.0) | 14.0 (1.2) | 7.6 (1.6) | 5.1 (1.7) |
| Completeness (%) | 99.7 (100.0) | 99.2 (98.4) | 100.0 (100.0) | 100.0 (100.0) |
| Redundancy | 6.4 (5.4) | 7.6 (7.6) | 7.1 (7.2) | 21.2 (21.9) |
| **Refinement** | | | | |
| $R_{work/}$ $R_{free}$ | 20.51 / 25.90 | 19.39 / 22.59 | 19.72 / 24.20 | 19.45 / 25.38 |
| No. atoms | | | | |
| Protein | 4450 | 3999 | 2035 | 5811 |
| Ligand/Ions | 2 | 16 | 10 | 111 |
| Water | 0 | 99 | 126 | 0 |
| B-factors | | | | |
| Protein | 75.2 | 47.4 | 34.1 | 66.1 |
| Ligand/Ions | 108.7 | 46.4 | 52.2 | 68.2 |
| Water | N.A. | 43.3 | 36.2 | N.A. |
| R.m.s deviations | | | | |
| Bond lengths (Å) | 0.011 | 0.011 | 0.010 | 0.014 |
| Bond angles (º) | 2.01 | 1.71 | 1.73 | 2.31 |
| Ramachandran plot (%) Favoured/Outliers | 87.1 / 3.6 | 97.5 / 0.0 | 92.6 / 2.0 | / |
| PDB | | | | |

Values in parenthesis are for the highest resolution shell. Rfree was calculated using a set (5%) of randomly selected reflections that were excluded from

**Supplementary Table 6│Crystallisation conditions for the GH16 enzymes**

| Enzyme (#) | Ligand | Condition | Cryoprotectant |
|---|---|---|---|
| Amuc_0724 (10 mg/ml) | apo | 1.6 M tri-sodium citrate pH 6.5 | Paratone-N |
| Baccac__02680$^{GH16}$ (8.1 mg/ml) | apo | 20 % PEG3350, 0.24 M sodium malonate pH7.0 | 25 % ethylene glycol |
| Baccac__02680$^{GH16E143Q}$ (8.1 mg/ml) | apo | 24 % PEG3350, 0.45 M sodium malonate pH7.0 | 25 % ethylene glycol |
| Baccac__02680$^{GH16\ E143Q}$ (8.1 mg/ml) | L404 | 24 % PEG3350, 0.45 M sodium malonate pH7.0 | 25 % ethylene glycol |
| Baccac__02680$^{GH16\ E143Q}$ (8.1 mg/ml) | TriLacNAc | 24 % PEG3350, 0.45 M sodium malonate pH7.0 | 25 % ethylene glycol |
| Baccac__03717$^{GH16}$ (10 mg/ml) | apo | 1.0 M Ammonium sulphate | 33 % ethylene glycol |
| BF4060$^{GH16}$ (9.6 mg/ml) | TriLacNAc | 20 % PEG6000, 1 M lithium chloride, 0.1 M Citric acid pH 4.0 | 25 % ethylene glycol |

### Initial concentration

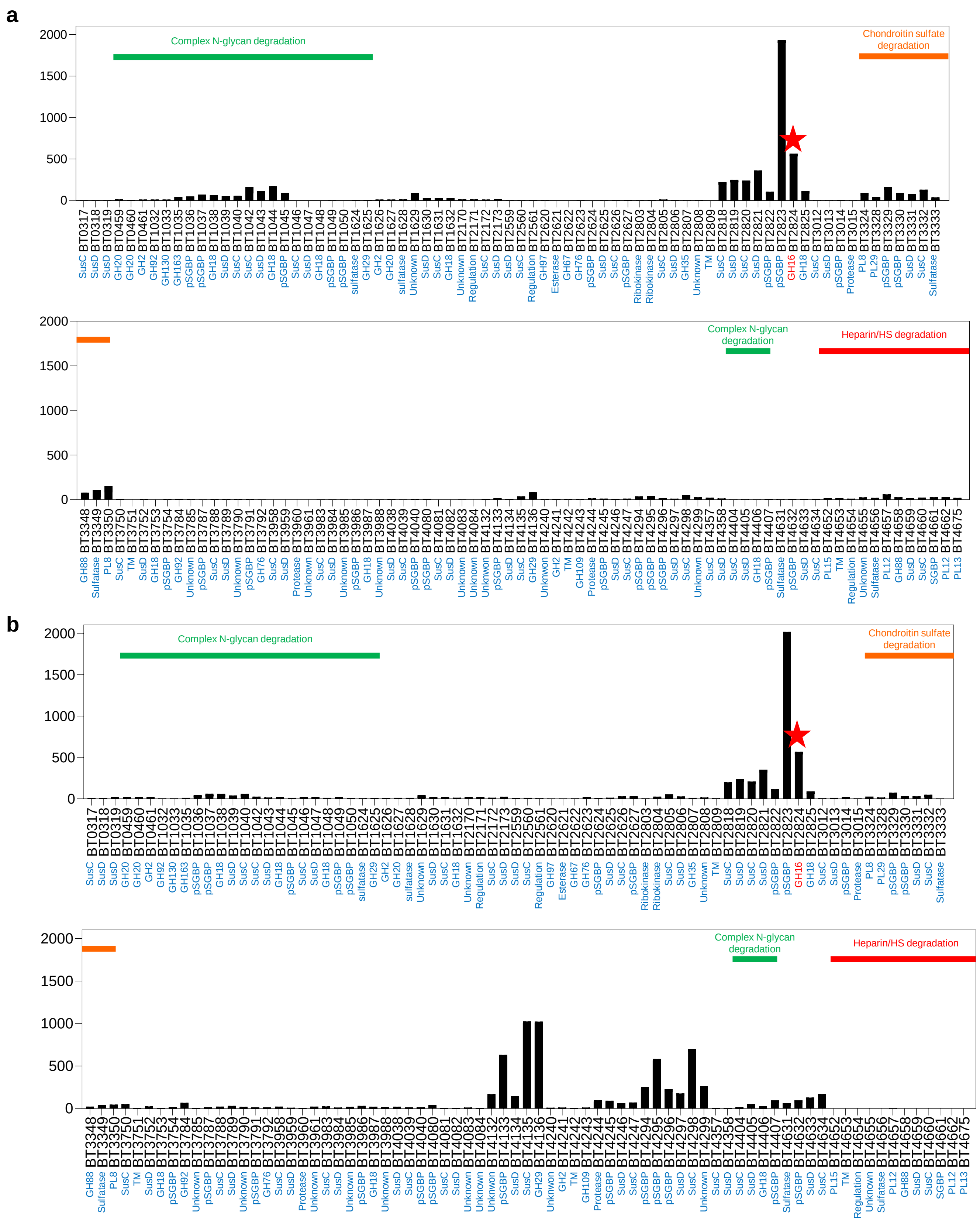


**Supplementary Figure 1│Upregulation of *Bacteroides thetaiotaomicron* genes on mucin O-glycans** Genes upregulated at least 10-fold *in vitro* using purified mucin O-glycans from PGM type III (Sigma) as the carbon source relative to glucose-grown cells (Martens *et al.* 2008) **a**, early in the growth curve. **b**, late in the growth curve. A number of PULs and non-PUL Cazyme containing loci were upregulated, including those involved in complex N-glycan and glycosaminoglycan degradation. The GH16 family member is indicated to by a red star

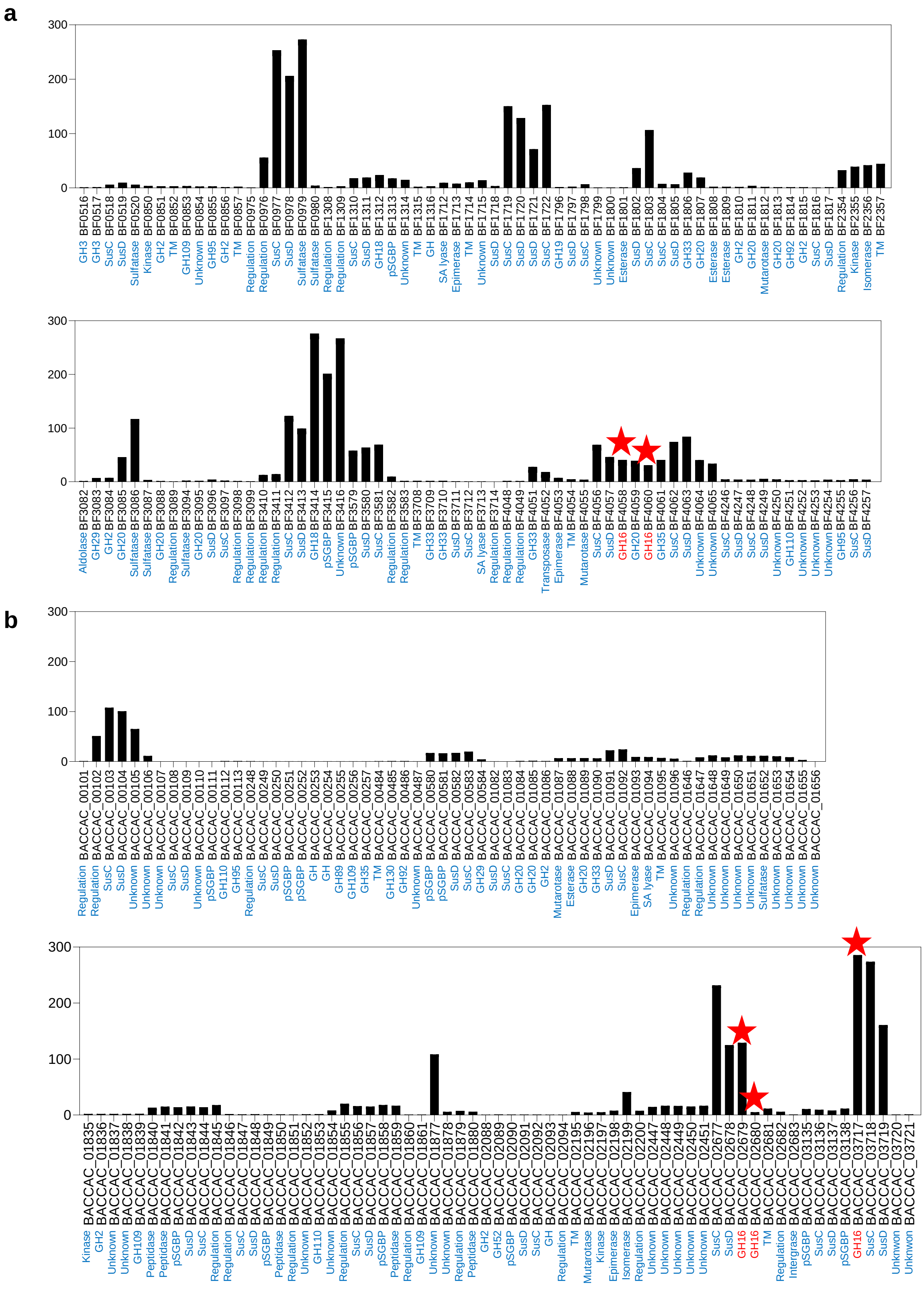


**Supplementary Figure 2 | Upregulation of *Bacteroides fragilis* and *Bacteroides caccae* genes on mucin O-glycans**
Genes upregulated at least 10-fold *in vitro* using purified mucin O-glycans from PGM III (Sigma) as the carbon source relative to glucose-grown cells **a**, *Bacteroides fragilis* (Pudlo *et al.* 2015). **b**, *Bacteroides caccae* (Desai *et al.* 2016). The GH16 family members are indicated to by a red star.

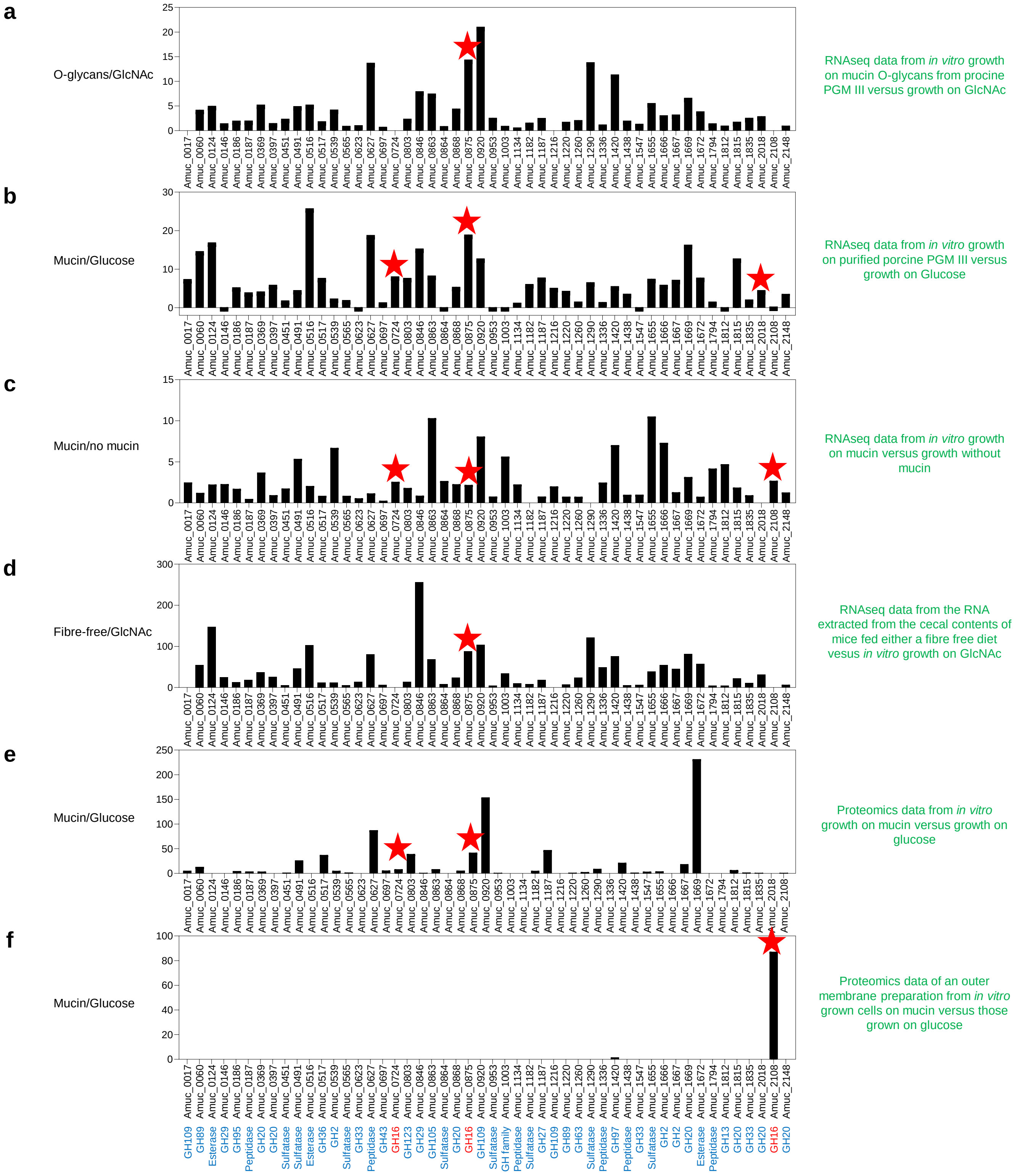


**Supplementary Figure 3│Upregulation of *A. muciniphila* genes and protein levels during growth on mucins in vitro and in vivo a**, Genes that are upregulated *in vitro* using purified mucin O-glycans as the carbon source relative to a culture grown on GlcNAc (Desai *et al.* 2016). **b**, Genes that are upregulated *in vitro* using purified porcine gastric mucin Type III (Sigma) as the carbon source relative to a culture grown on GlcNAc (Ottman *et al.* 2017). **c**, Genes that are upregulated *in vitro* using purified mucin O-glycans as the carbon source relative to a culture grown on GlcNAc (Shin *et al.* 2019). **d**, Genes that are upregulated when a mouse is fed a fibre-free diet compared to a simple sugar diet (Desai). **e**, Relative protein levels during *in vitro* growth on purified PGM III compared to growth on glucose (Ottman *et al.* 2016). **f**, Relative protein levels in the outer membrane fraction during growth on purified PGM III compared to growth on glucose (Ottman *et al.* 2016). It should be noted that the *in vitro* growths a-c were performed in different basal medias. The GH16 family members are indicated by a red star.

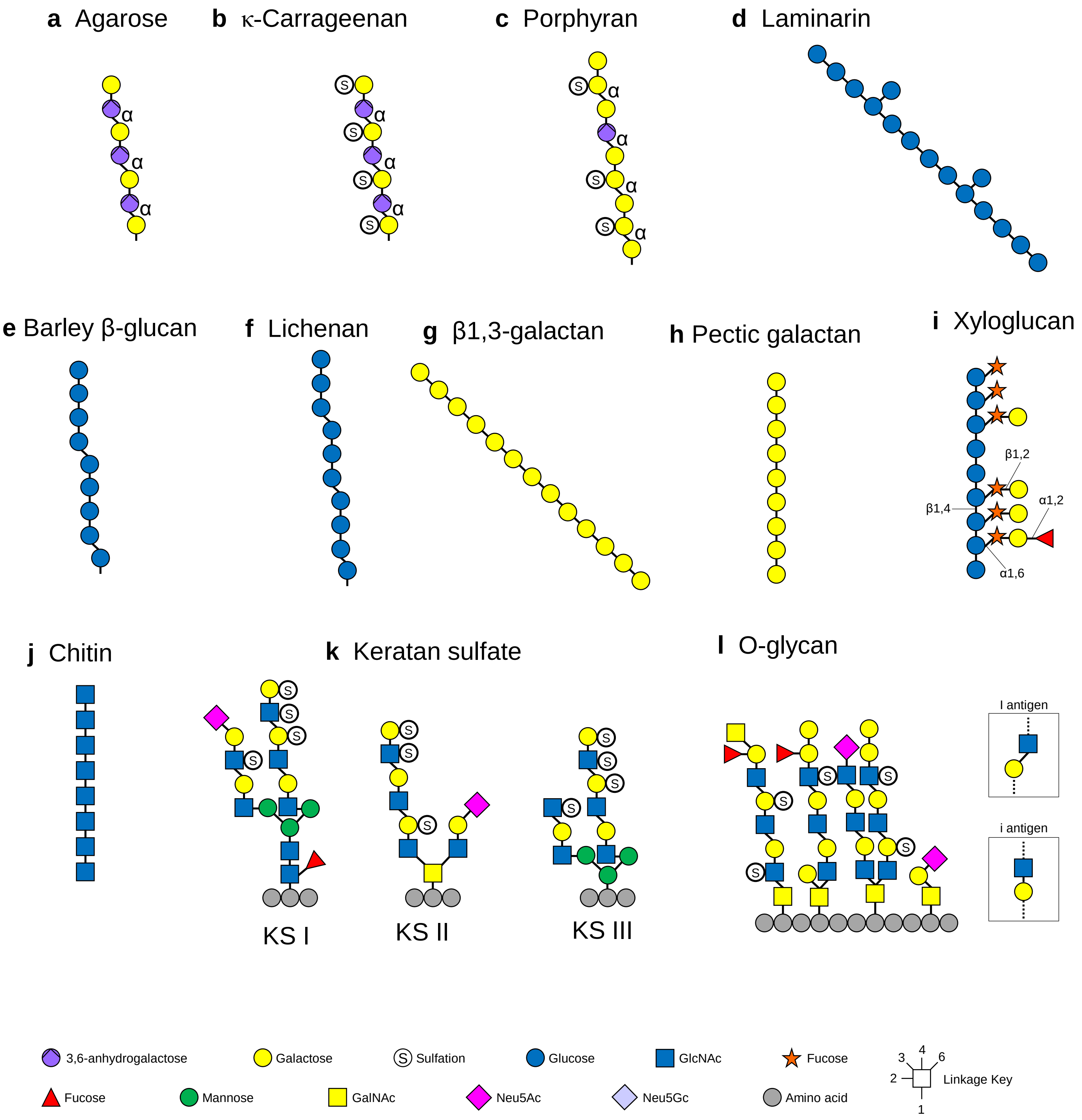


**Supplementary Figure 4│Classes of glycan targeted by GH16 enzymes. a**, Agarose (α1,3-3,6-anhydrogalactose-β1,4-galactose repeating units). **b**, κ-carrageenan (α1,3-3,6-anhydrogalactose-β1,4-galactose6S repeating units). **c**, Porphyran (α1,4-galactose6S-β1,3-galactose repeating units where the galactose6S is sometimes replaced with 3,6-anhydrogalactose). **d**, Laminarin (β1,4-glucan with occasional β1,6-glucose decoration). **e**, Barley β-glucan (β1,4-glucan with occasional β1,3-linkages). **f**, Lichenan (predominantly β1,4-glucan with ~25 % β1,3-linkages). **g**, β1,3-galactan (arabinogalactan backbone; β1,3-linkages). **h**, Pectic galactan (β1,4-linkages). **i**, Xyloglucan (linkages displayed do not follow the key, but are labelled). **j**, Chitin (β1,4-linkages). **k**, Keratan sulfate (polyLacNAc structures that can be O- or N-linked to protein, 6S decoration possible on galactose and GlcNAc and occasional sialylation and fucosylation). **l**, Mucin O-glycans. In keratan sulfate and O-glycan structures the GlcNAc sugars can also be linked through β1,4 and β1,6 linkages (boxes).

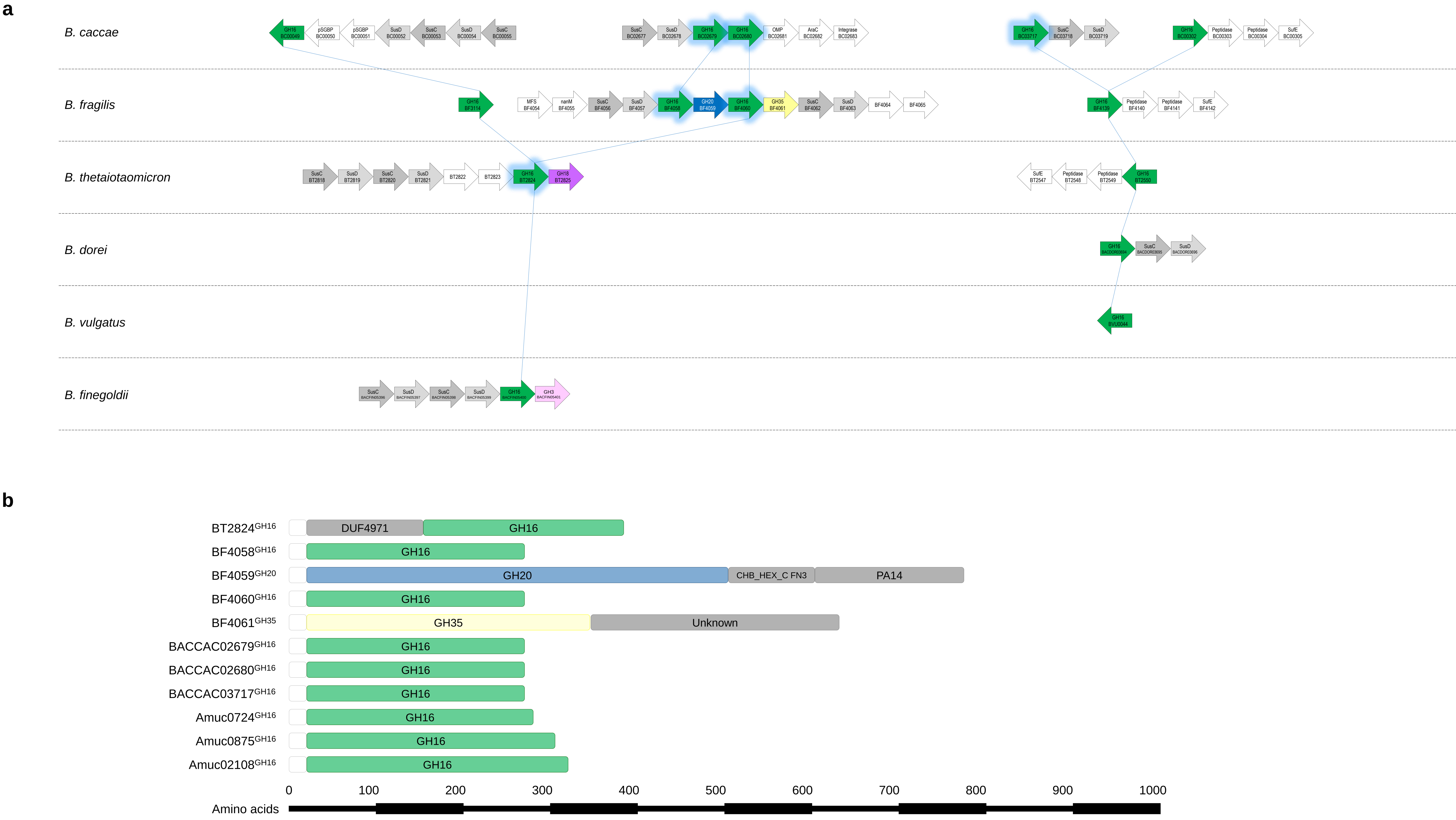


**Supplementary Figure 5│Genomic context of the O-glycanase GH16 genes from different *Bacteroides* species and the predicted protein domains of the CAZymes explored. a,** The GH16 enzymes upregulated on mucin were used to search for homologues in other *Bacteroides* spp. (see Materials and Methods). The GH16 enzymes upregulated on O-glycans are highlighted with a blue glow. The lines connect the homologues. GH and other protein families are colour-coded: GH20 (blue), GH35 (yellow), GH18 (purple), GH3 (pink) and SusC/D-like pairs (grey). Abbreviations include: MFS - major facilitator superfamily. **b,** The domain structure and approximate lengths, which were determined as described in Materials and Methods. The white boxes at the start indicate the signal sequence of the protein.

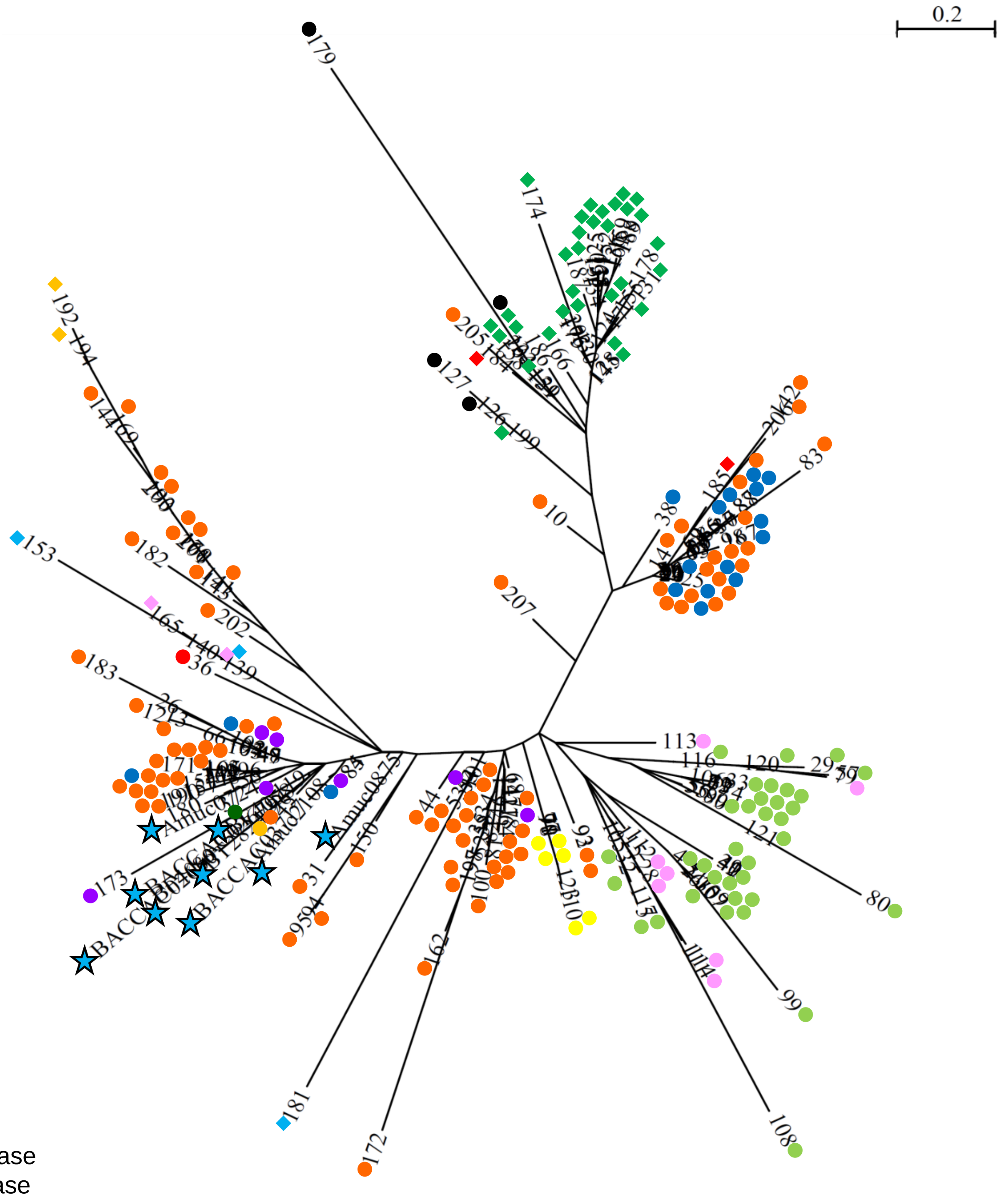


**Supplementary Figure 6│Phylogenetic tree of characterised GH16 family members.** The sequences of GH16 family members with reported activities (CAZy database) and the O-glycan active family members characterised in this report (blue stars) were compared as described in Materials and Methods. Each CAZy data base entry was given a number to simplify the tree. Sub-activities can be seen branching off together in many instances.

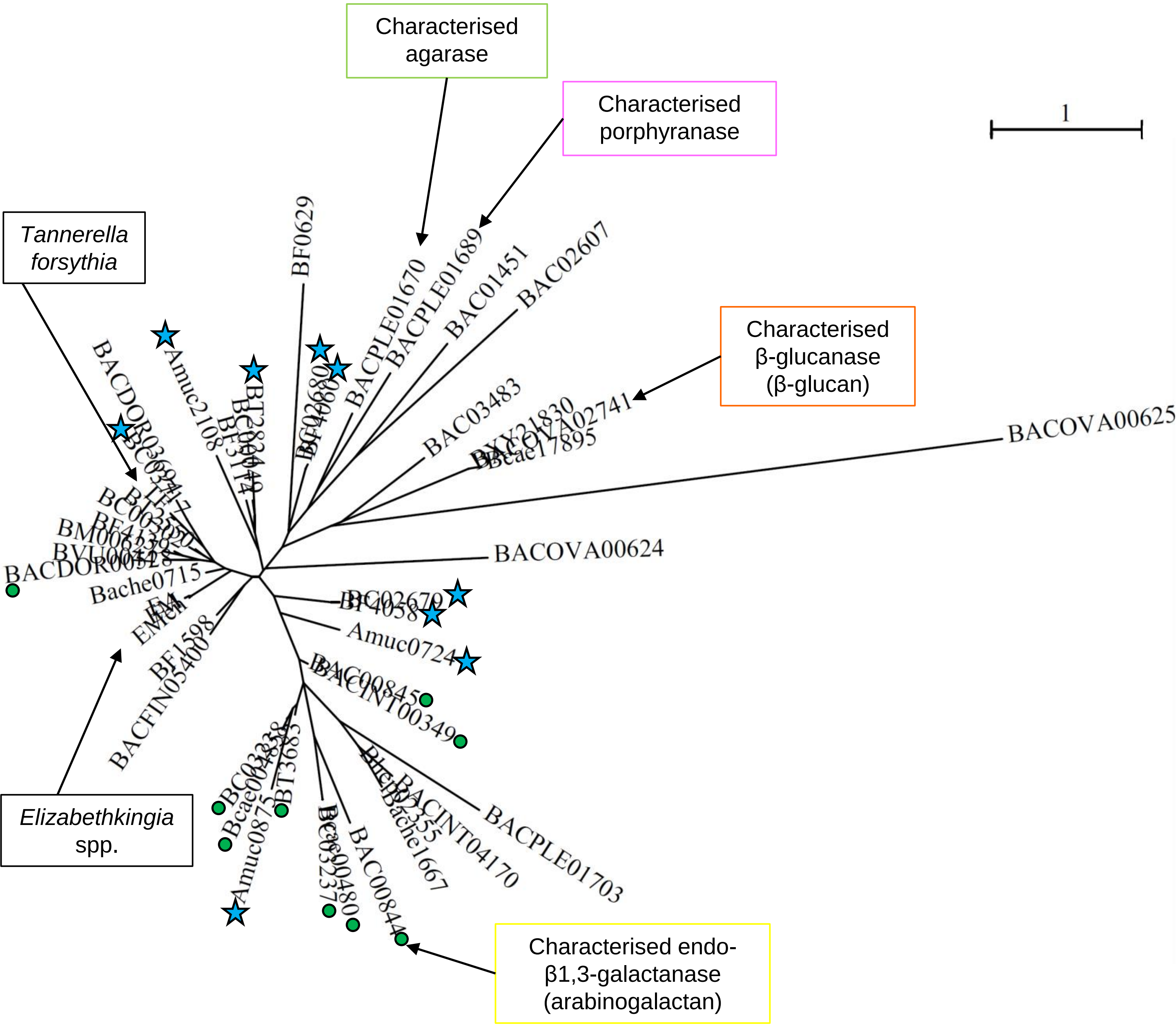


**Supplementary Figure 7│Phylogenetic tree of a selection of characterised and uncharacterised GH16 family members.** 47 characterised and uncharacterised GH16 family members were selected from 14 mutualistic *Bacteroides* spp., *A. muciniphila* and 4 pathogens from the Bacteroidetes phylum. A key to the abbreviations of the locus tags is provided in Supplemenatry Table X. The pathogens include *Tanneralla forsythia* and the *Elizabethkingia* species *E. meningoseptica*, *E. miricola* and *E. anophelis*. The O-glycan active GH16 family members described in this report are indicated by a blue star. The characterised GH16 family members include a β-glucanase involved in degrading mixed-linkage β-glucan from *B. ovatus* (Tamura *et al.* 2017), a porphyranase and agarose from *B. plebius* (Heheman *et al.* 2012) and a β1,3-galactanase from *B. cellulosilyticus* involved in the degradation of arabinogalactan (Cartmell *et al.* 2018). Those GH16 enzymes that are in PULs with a similar gene composition to the characterised arabinogalactan GH16 from *B. cellulosilyticus* are highlighted by a green dot.

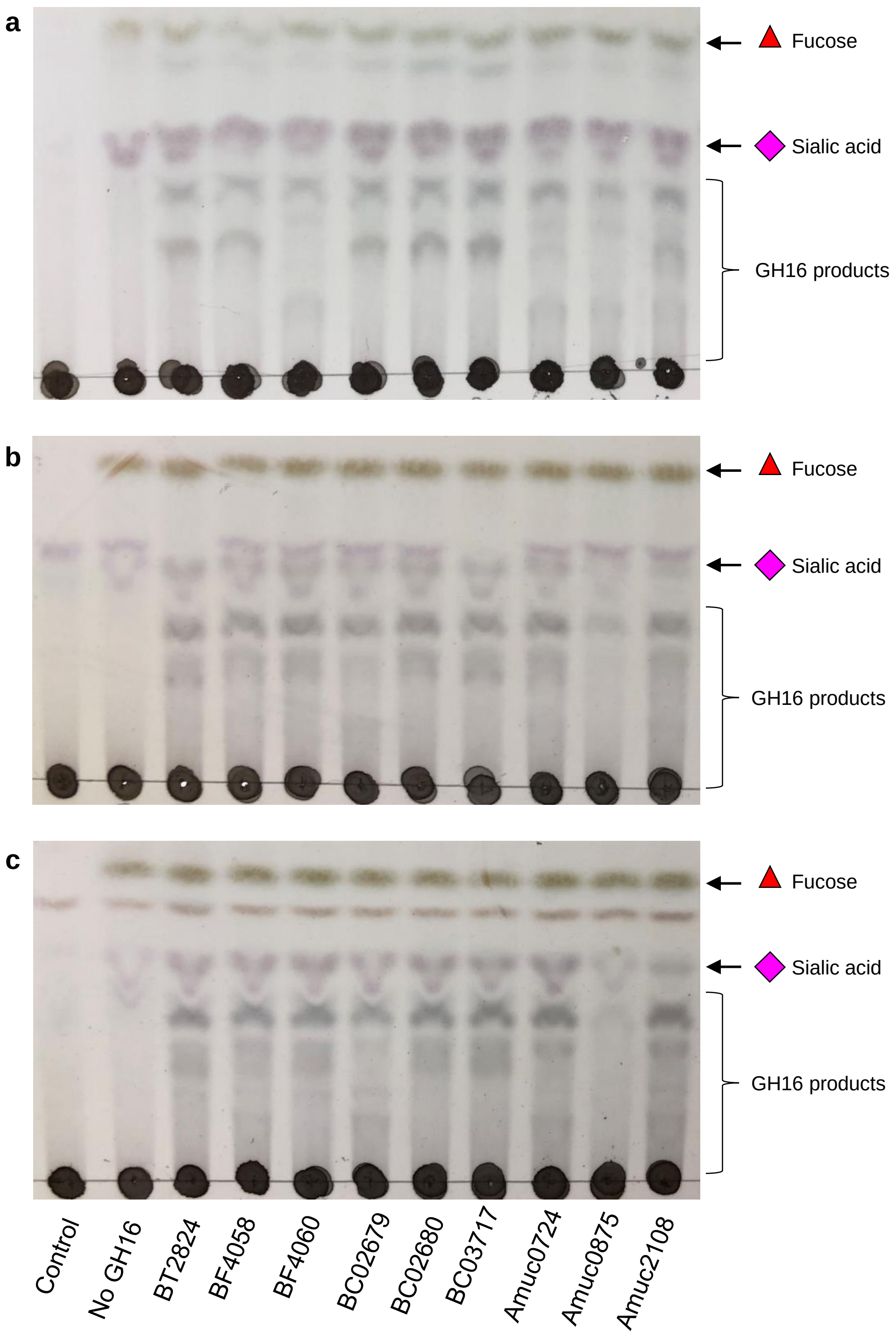


**Supplementary Figure 8│Activity of the GH16 enzymes against porcine small intestinal mucin and commercially available Porcine gastric mucin a**, Porcine small intestinal mucin. **b**, PGM type II. **c**, PGMtype III. All assays with a GH16 enzyme also included a sialidase and α1,2-fucosidase, BT0455$^{GH33}$ and a GH95 (from *Bifidobacterium bifidum*; see Supplementary Fig. 9).

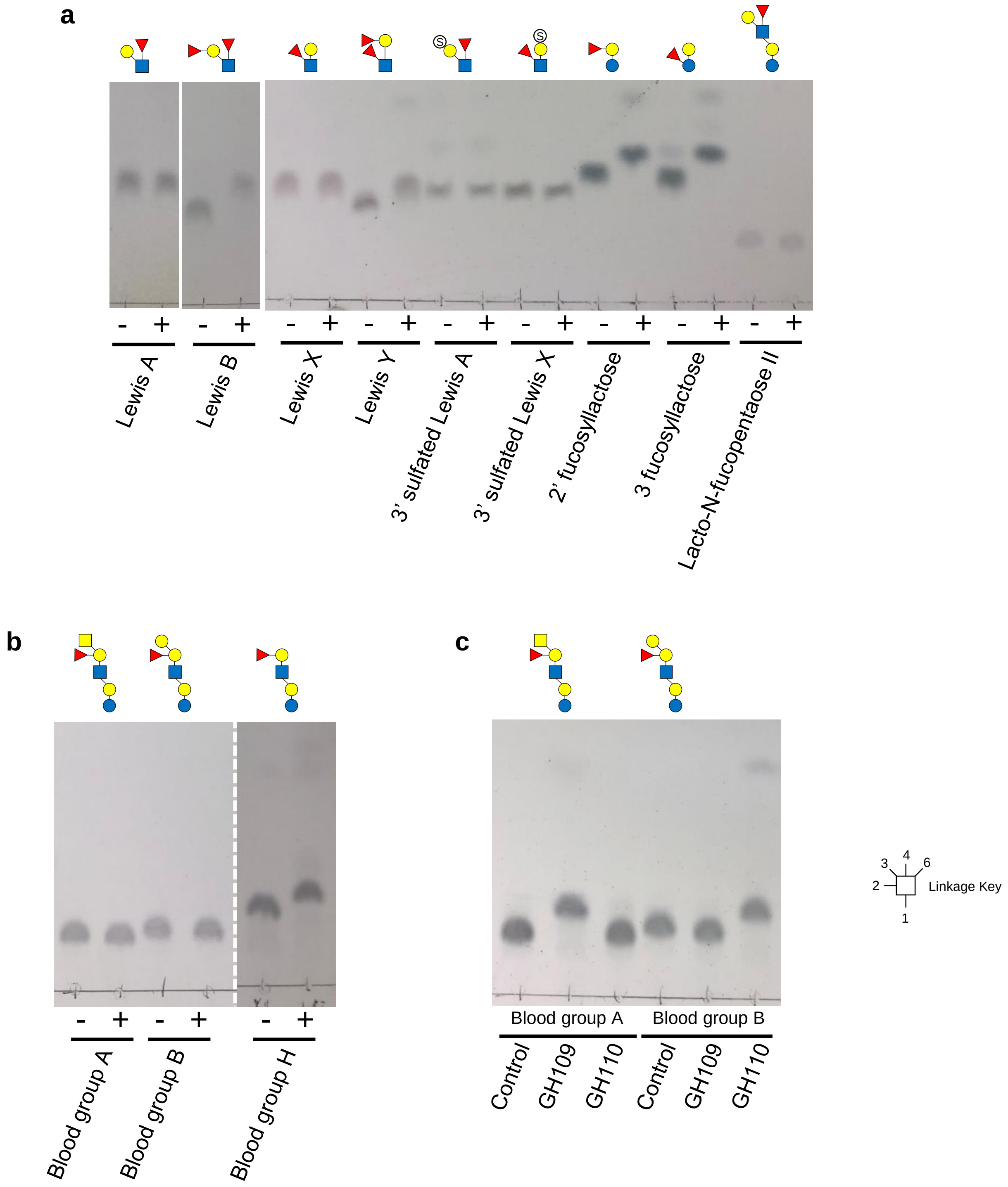


**Supplementary Figure 9│Activity of exo-acting enzymes against di-and oligo-saccharides a,** GH95 α-fucosidase from *Bifidobacterium bifidum* **b**, GH95 α-1,2-fucosidase from *Bifidobacterium bifidum* against blood group sugars. Activity can only be seen after removal of the α-linked GalNAc or Gal (blood group H). **c**, Activity of a GH109 α-GalNAc'ase and GH110 α-galactosidase against different blood group structures.

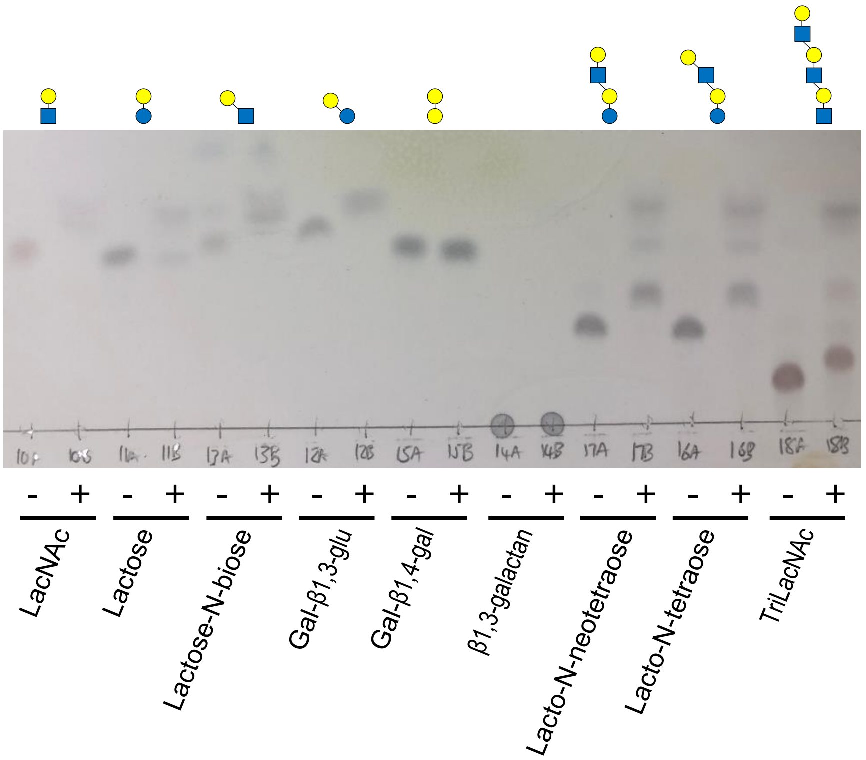


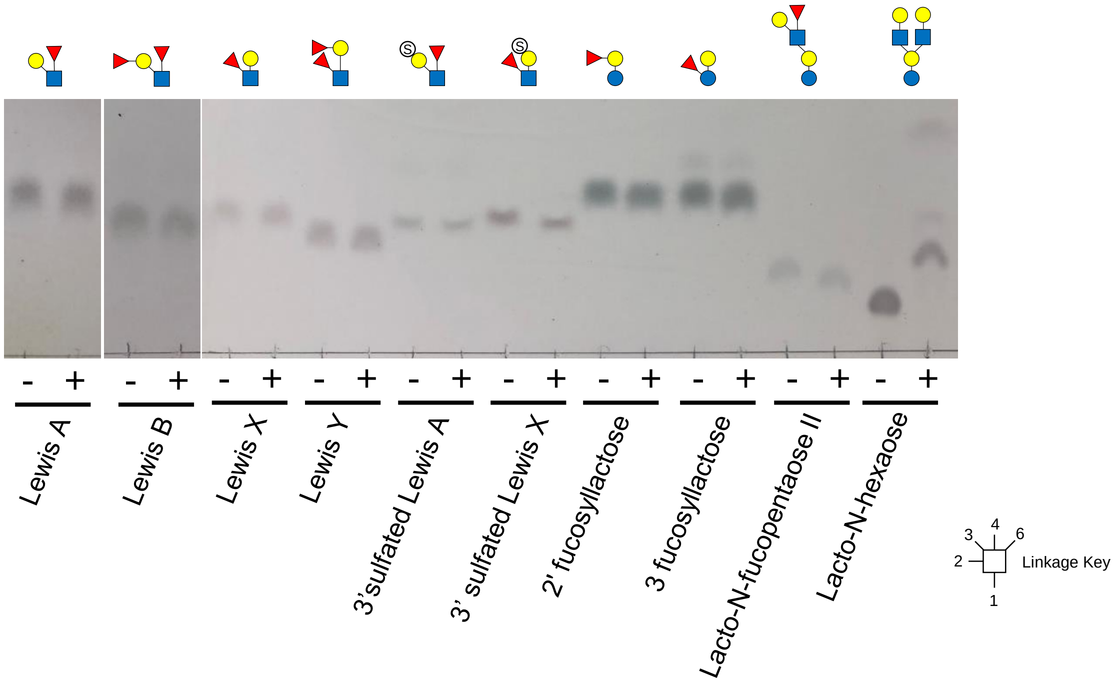


**Supplementary Figure 10 │ Activity of galactosidase BF4061$^{GH35}$ di- and oligosaccharides**
The polysaccharide utilisation loci in *Bacteroides fragilis* encoding the O-glycan active GH16 enzymes also encodes a putative GH35. The recombinant form was incubated with a variety of glycans to determine specificity.

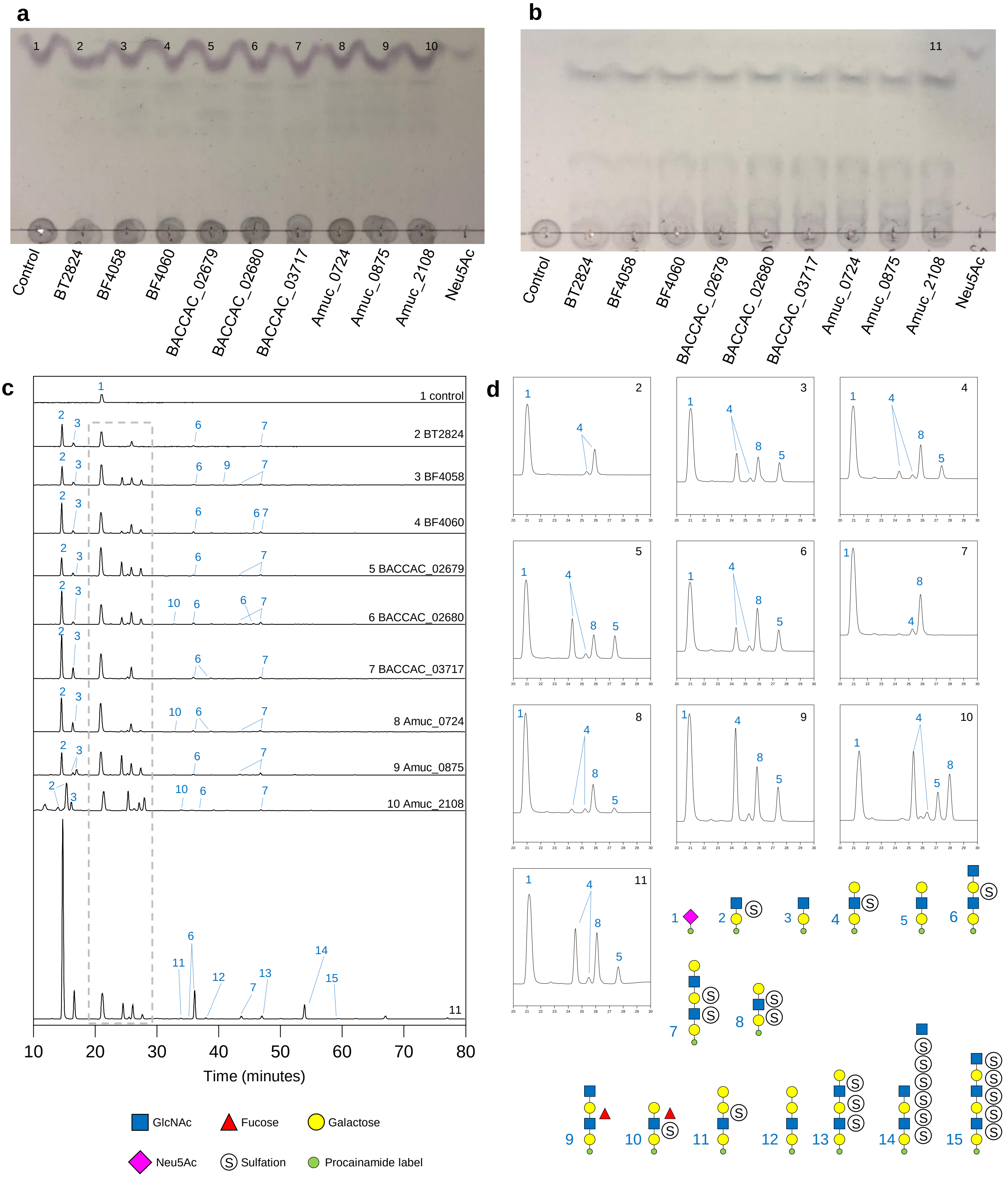


**Supplementary Figure 11│Products produced by O-glycan active GH16 family members on keratan a**, TLC of activity against N-linked egg keratan. **b**, TLC of activity against N-linked bovine cornea keratan. **c**, Chromatograms of procainamide-labelled keratan products, where the number correspond to those in a and b. The grey dotted box indicated the areas which are magnified in d. **d**, Magnifications of the chromatograms between 20-30 minutes to allow clearer annotation of the products. All assays also included the broad-acting sialidase BT0455$^{GH33}$. The composition and structure of the products are shown.

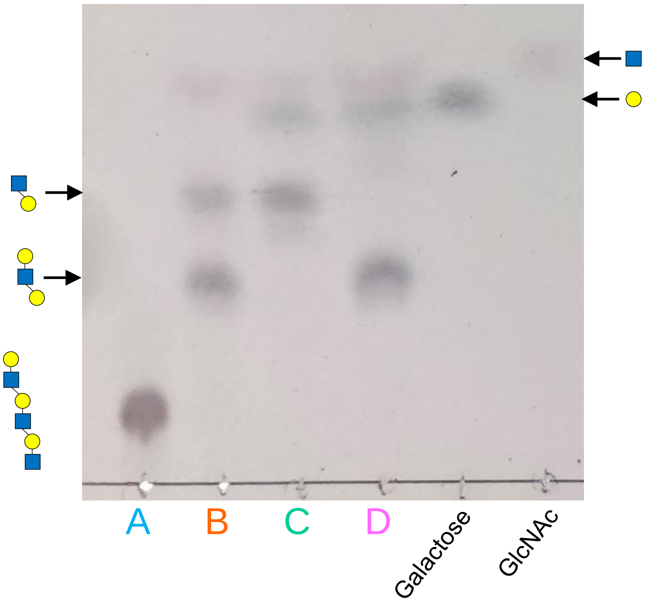


A. Control
B. Amuc_0724$^{GH16}$
C. Amuc_0724$^{GH16}$ post-treated with β1,4-galactosidase (BT0461$^{GH2}$)
D. Amuc_0724$^{GH16}$ post-treated with broad-acting GlcNAc'ase (BT0459$^{GH20}$)

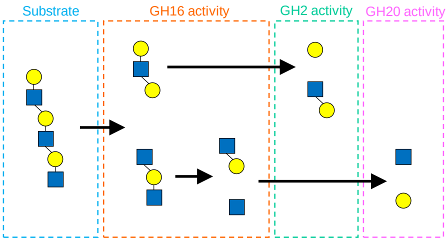


**Supplementary Figure 12│Activity of Amuc_0724$^{GH16}$ against triLacNAc** The products of triLacNAc digestion by Amuc_0724$^{GH16}$ (lane B) were incubated with either a β1,4-galactosidase (lane C) or an β-GlcNAc'ase (lane D) to confirm their identity.

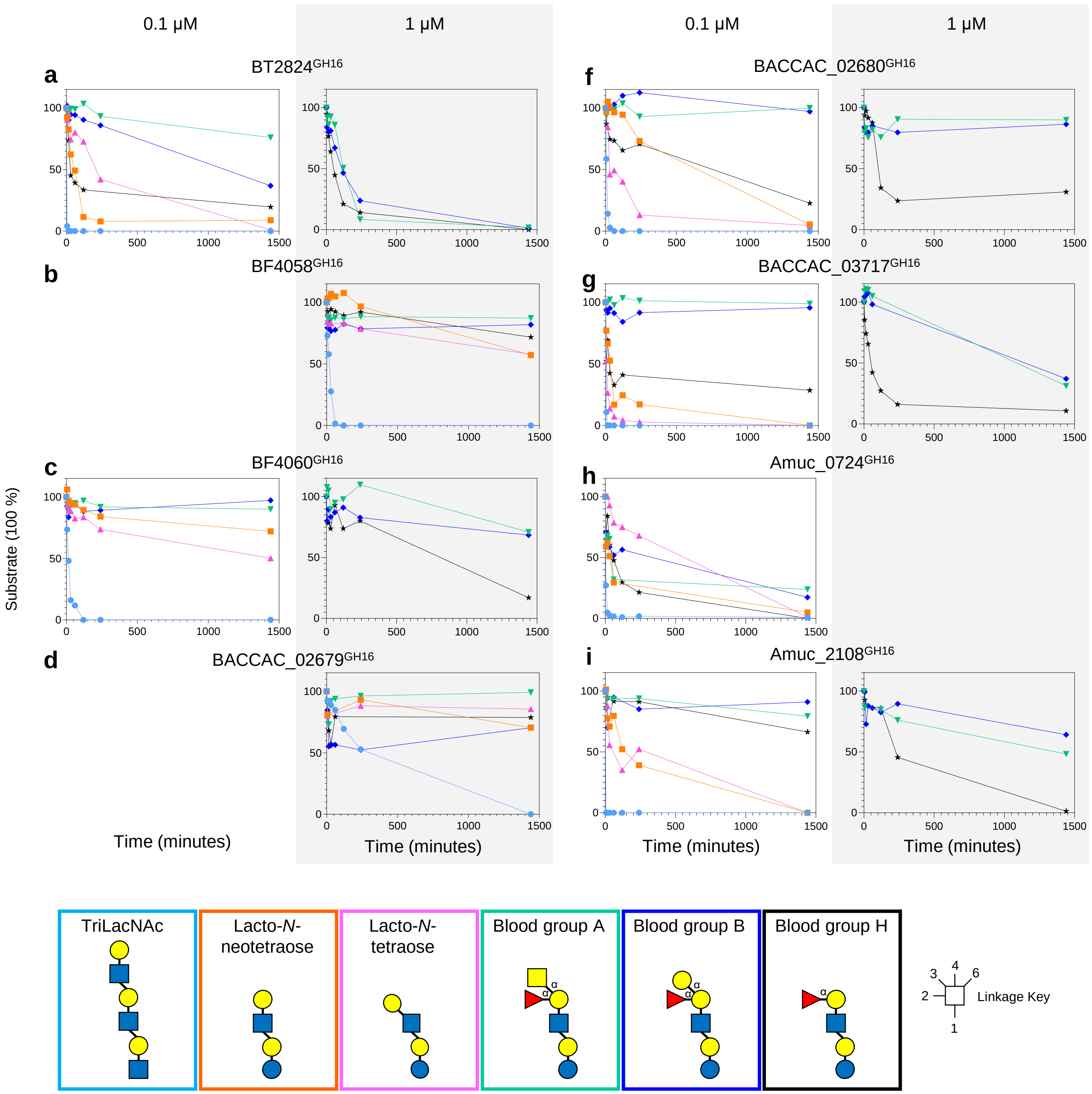


**Supplementary Figure 13│Substrate depletion assays with defined oligosaccharides to probe the importance of the different sub-sites of the GH16 enzymes** Assays were carried out with 1 mM substrate concentration and samples taken at different time points to monitor substrate depletion by HPAEC-PAD. Enzyme concentration was 0.1 µM (white columns) or 1 µM (grey columns). The defined oligosaccharides used were TriLacNAc (light blue circles), Lacto-*N*-neotetraose (orange squares), Lacto-*N*-teraose (pink triangles), blood group A (green inverted triangles), blood group B (dark blue diamonds) and blood group H (black stars). **a**, BT2824$^{GH16}$. **b**, BF4058$^{GH16}$. **c**, BF4060$^{GH16}$. **d**, BACCAC_02679$^{GH16}$. **e**, BACCAC_02680$^{GH16}$. **f**, BACCAC_03717$^{GH16}$. **g**, Amuc_0724$^{GH16}$. **h**, Amuc_0875$^{GH16}$. **i**, Amuc_2108$^{GH16}$.

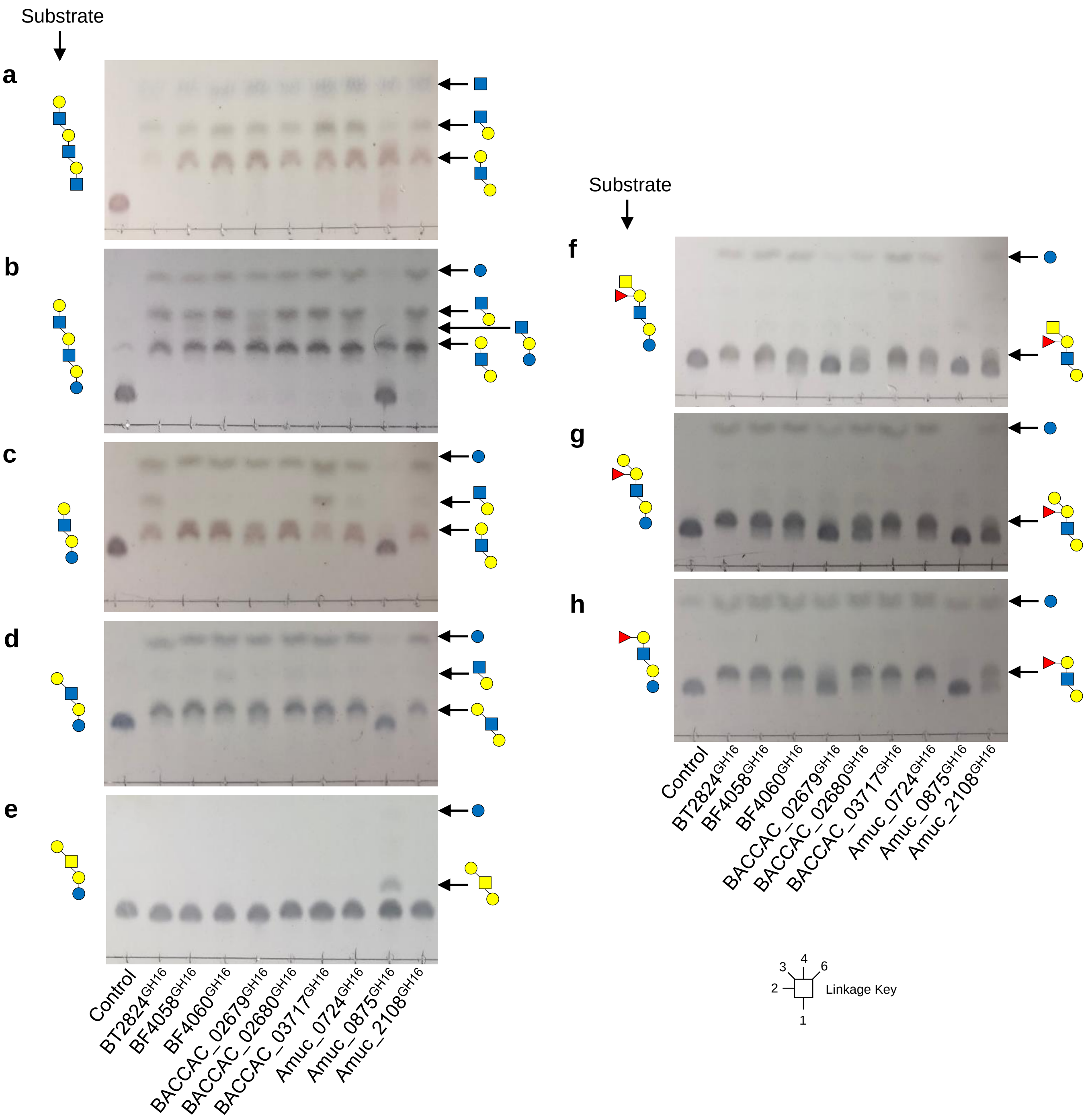


**Supplementary Figure 14│Activity of the GH16 enzymes against defined oligosaccharides.** Products of GH16 activity were visualised by TLC. **a** triLacNAc. **b**, paraLacto-N-neohexaose. **c**, Lacto-N-neotetraose. **d**, Lacto-N-tetraose. **e**, Galβ1,3GalNAc β1,3Galβ1,4Glc. **f**, Blood group A hexasaccharide II. **g**, Blood group B hexasaccharide II. **h**, Blood group H pentasaccharide generated from A or B. Control = no enzyme added.

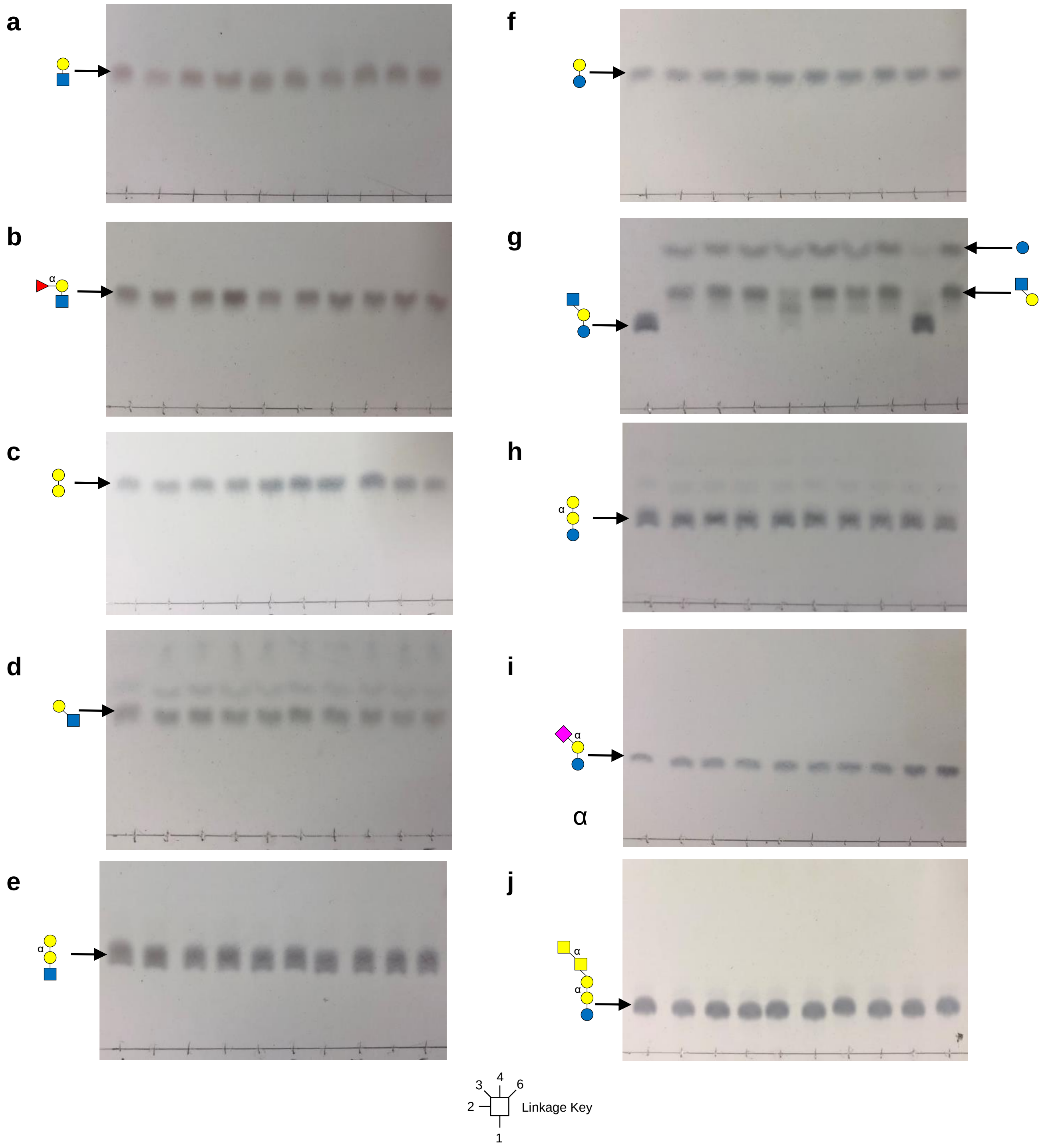


**Supplementary Figure 15│Activity of the GH16 enzymes defined di- and oligo-saccharides.** Products of GH16 activity were visualised by TLC. **a**, LacNAc. **b**, Blood group H trisaccharide II. **c**, Galβ1,4Gal. **d**, Lacto-N-biose. **e**, P1 antigen. **f**, Lactose. **g**, Lacto-N-triose. **h**, Globotriose. **i**, 3-Sialyllactose. **j**, Forssman antigen.

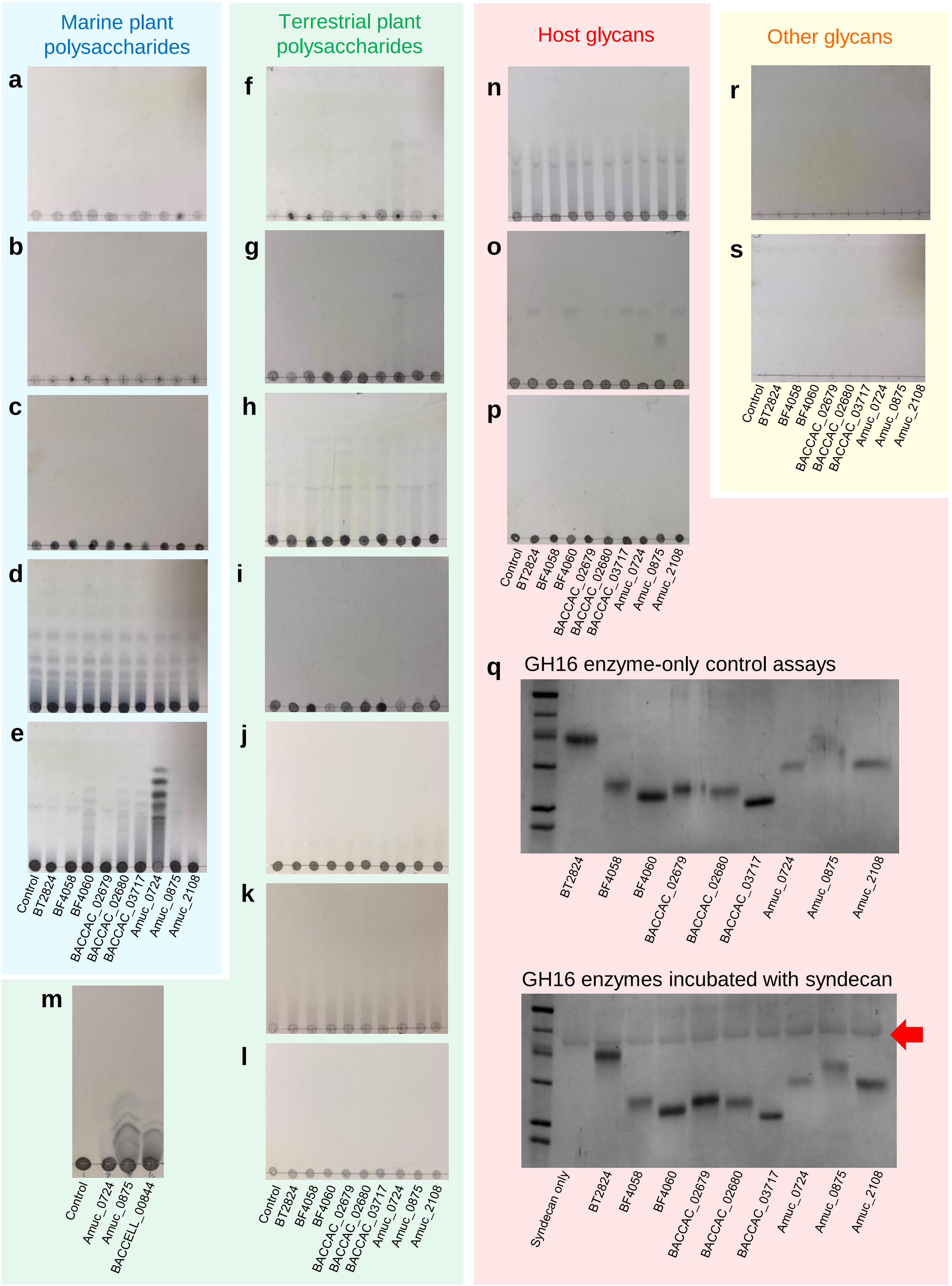


**Supplementary Figure 16│Activity of the GH16 enzymes characterised in this work against polysaccharides already associated with the family** Activity of GH16 enzymes against substrates associated with this family. All assays were carried out with **a**, Agar. **b**, Agarose. **c**, κ-carrageenan. **d**, Porphyran. **e**, Laminarin. **f**, Barley β-glucan. **g**, Lichenan. **h**, Pectic β1,4-galactan. **i**, Xyloglucan. **j**, Larch arabinogalactan. **k**, Wheat arabinogalactan. **l**, Gum arabinogalactan. **m**, β1,3-galactan **n**, Chondroitin sulfate. **o**, Heparan sulfate. **p**, Hyaluronic acid. **q**, Human syndecan. To assess if the GH16 enzymes could cleave heparan sulfate polysaccharides away from protein they were incubated with human syndecan (bottom gel). A control gel (top) shows GH16-only assays. The red arrow indicates where syndecan (50 μg) runs on the gel and shows that it does not decrease in mass with the addition of GH16. **r**, Shrimp chitin. **s**, Squid chitin. All reactions were carried out in phosphate buffer at pH 7 overnight with 3 μm enzyme.

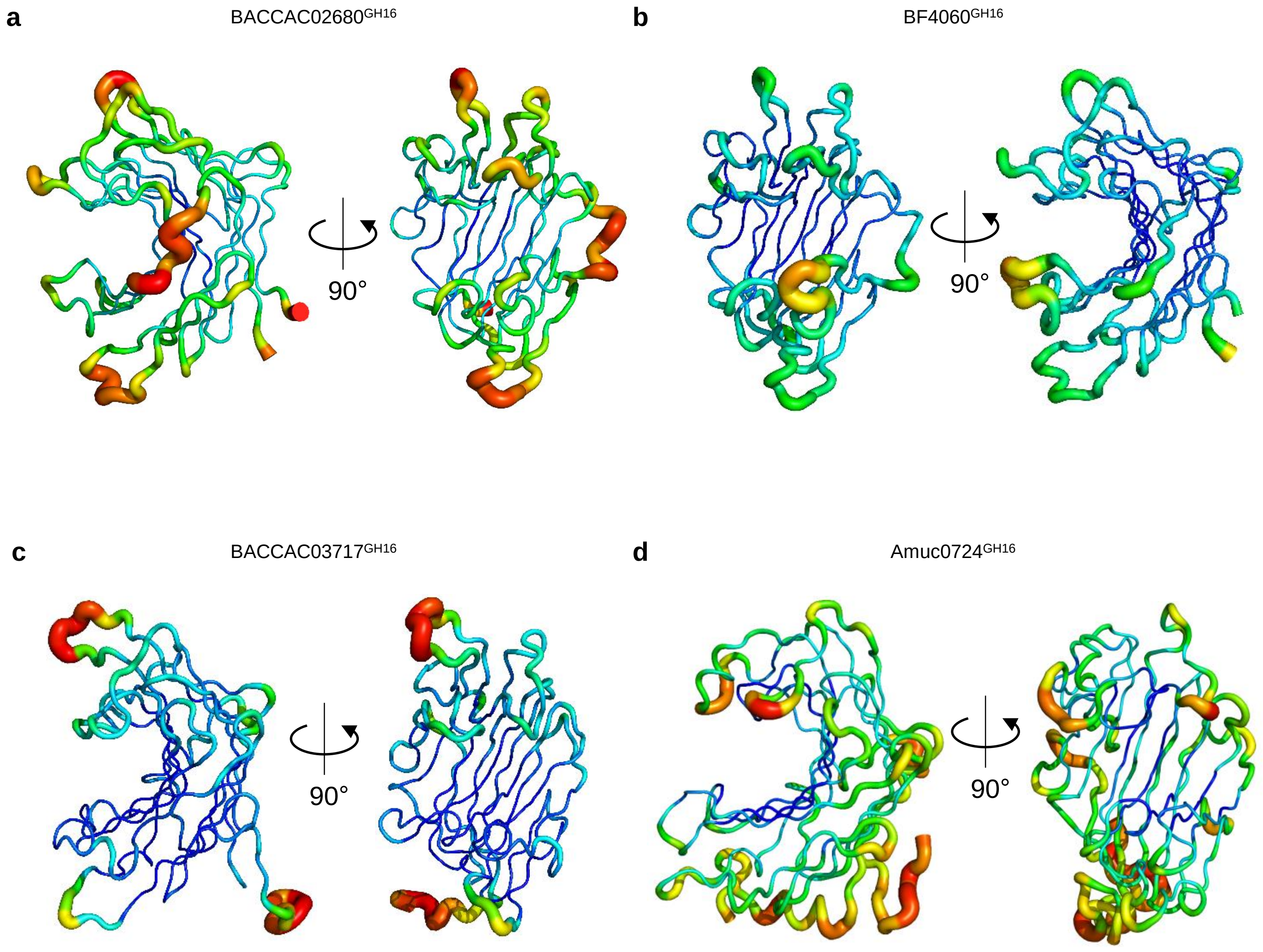


**Supplementary Figure 17│B factor putty projections of the O-glycan active GH16 crystal structures obtained in this study**. **a**, Baccac_02680$^{GH16E143Q}$. **b**, BF4060$^{GH16}$. **c**, Baccac_03717$^{GH16}$. **d**, Amuc_0724$^{GH16}$.

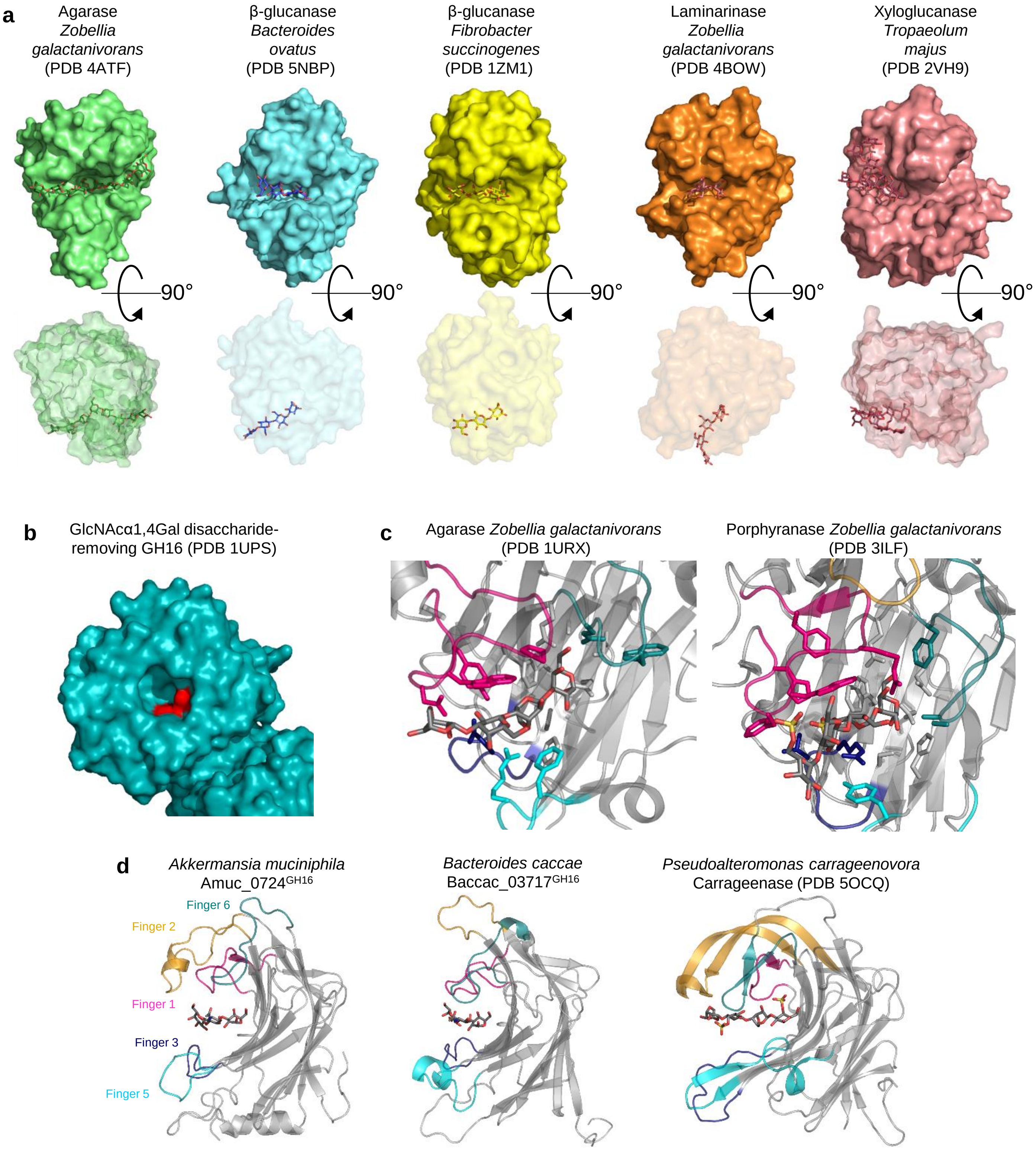


**Supplementary Fig. 18│Crystal structures differences within the GH16 family members a**, Crystal structures of a number of different GH16 family members to exemplify their variation in terms of cleft structure and orientation of substrate (Heheman *et al.* 2012, Tamura *et al.* 2017, Tsai *et al.* 2005, Labourel *et al.* 2014 and Mark *et al.* 2009). **b**, The substrate binding site of *Clostridium perfringens* GH16 family member that removes GlcNAcα1,4Gal disaccharide from stomach O-glycans. This pocket-like binding site is a unique example amongst the GH16 family structures so far. Catalytic residues are shown in red (Tempel *et al.* 2005). **c**, The active sites of two GH16 enzymes with activity towards marine plant polysaccharides. This demonstrates the differences in the negative subsites to GH16 β-glucanases. Both types of enzyme select for a β1,3 linkage between the -1 and -2 subsite, but the structural reasons driving this are different. **d**, Side views of Amuc_0724$^{GH16}$, BACCAC_03717$^{GH16}$ and a carrageenase (Matard-Mann *et al.* 2017) to demonstrate the different types of finger two (yellow) and their interactions with sugars occupying the -3 and -4 subsites.

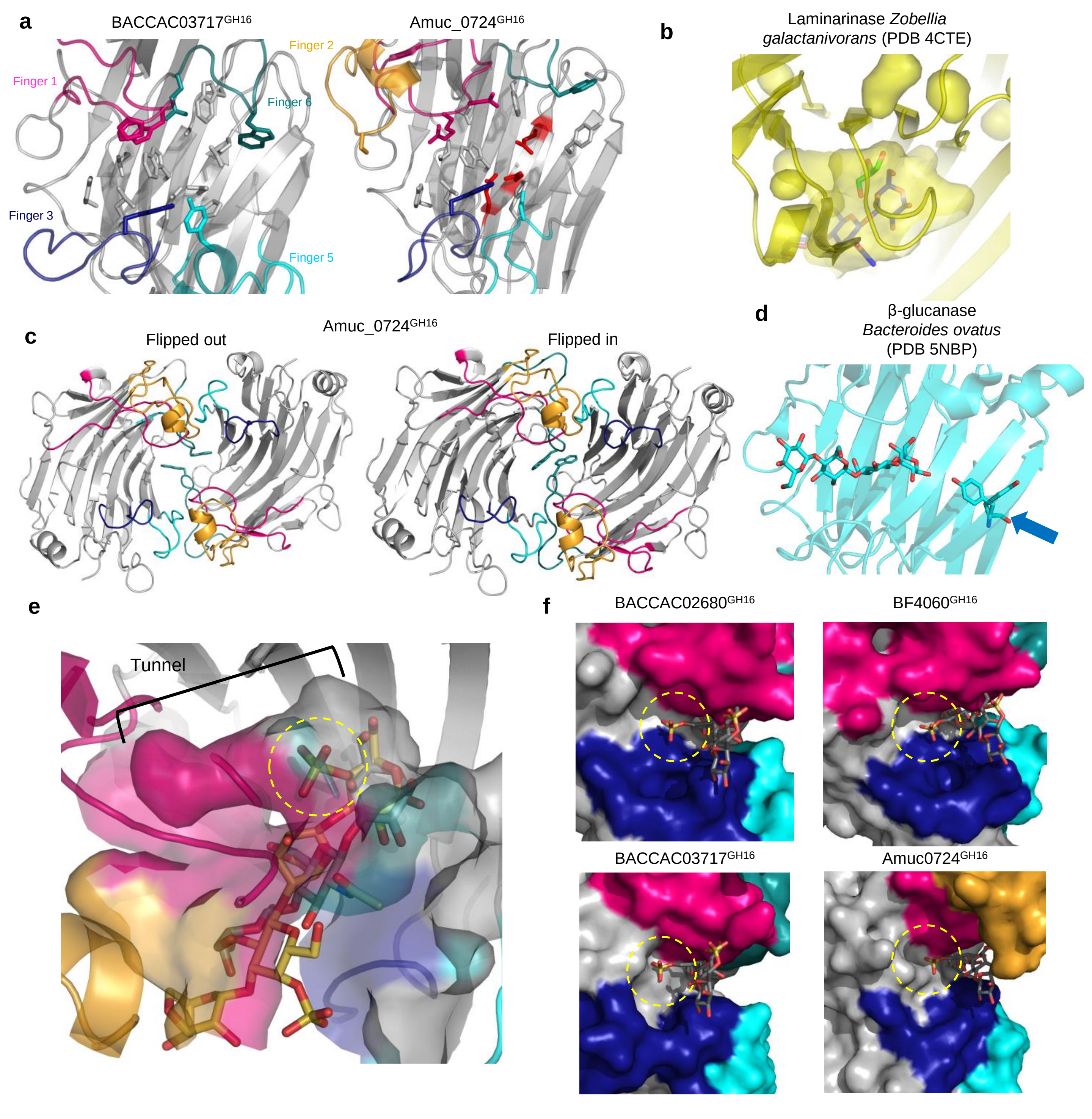


**Supplementary Fig. 19│Crystal structures comparisons within the GH16 family members a**, The detailed view of the active sites of Baccac_03717$^{GH16}$, and Amuc_0724$^{GH16}$. The fingers are colour coded. **b**, A surface representation of the cavities and pockets around the -1 subsite of a laminarinase (Labourel *et al.* 2015). A glycerol was crystallised within the pocket adjoining the -1 subsite and the product from Baccac_02680$^{GH16E143Q}$ is overlaid to show the -1 subsite. The presence of the glycerol, suggested that branching sugars may be accommodated at this position. **c**, The two different occupancies possible for W279 in Amuc_0724$^{GH16}$ in the unit cell to show the interactions between the two molecules. The tyrosines are show as sticks and come from finger 6 (teal). The flipped-out version may represent a crystallographic artefact rather than a biologically relevant observation. **d**, An example of a dynamic tyrosine (blue arrow) also observed in the positive subsites of a crystal structure of a β-glucanase GH16 family members from *B. ovatus*. **e**, A surface representation of the pockets and cavities around the negative subsites of the Amuc_0724$^{GH16}$ crystal structure. The different colours represent the different fingers of the enzyme. The trisaccharide product from Baccac_02680$^{GH16E143Q}$ is overlaid into the structure (grey) as well as the carrageenan product (yellow) from a κ-carrageenases from *Pseudoalteromonas carrageenovora* (PDB 5OCQ; Matard-Mann *et al.* 2017) to show how a sulfate could be accommodated in the tunnel of Amuc_0724$^{GH16}$ (yellow dashed circle). Accommodation of this sulfate group would not be possible with the O-glycan active GH16 enzymes from *Bacteroides* spp. **f**, The four O-glycan active GH16 enzymes from this study are overlaid with the porphyran product originally crystallised with a porphyranase from *Zobellia galactanivorans* (PDB 3ILF; Heheman *et al.* 2010). This demonstrates how a sulfate group could be accommodated in the cleft at the -2 subsite of these enzymes (yellow dashed circle).

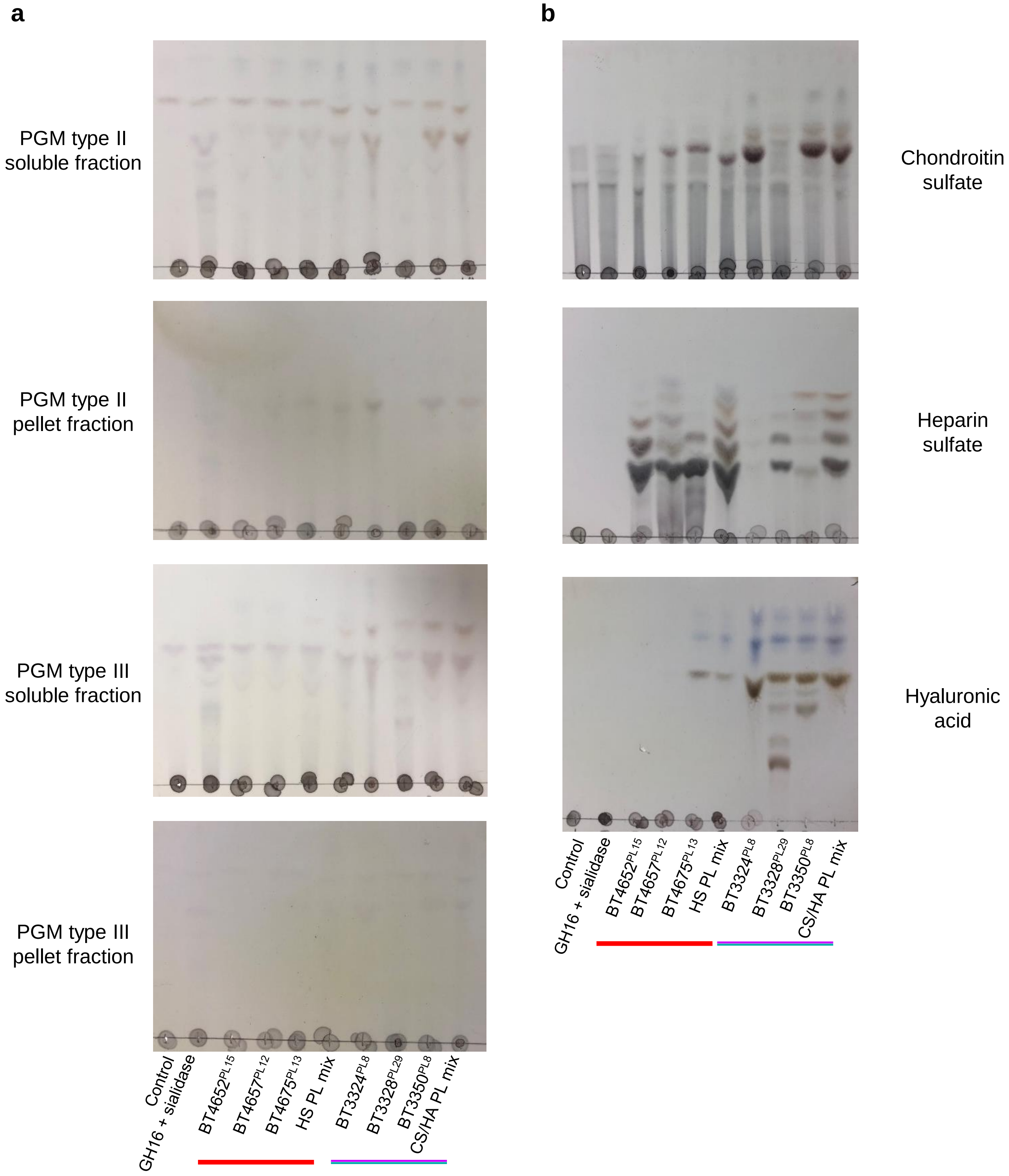


**Supplementary Figure 20│Analysis of the glycosaminoglycan (GAG) content of porcine gastric mucin (PGM) type II and III. a,** The polysaccharide lyases from *B. thetaiotaomicron* involved in or predicted to be involved in the degradation of HS, CS and HA were used to test for the presence of these GAGs in PGM types II and III. PGM was dissolved in DI $H_2O$ at 50 mg/ml then centrifuged at 16000 x g for 30 min. The supernatant (soluble) and pellet fractions were assayed separately. Each polysaccharide lyase was tested individually and then as a mix (Cartmell *et al.* 2017; Ndeh *et al.* 2018). The red line and purple/teal line represents the HS and CS/HA active lyases, respectively. **b,** The Polysaccharide lyases were tested against pure GAGs as controls.

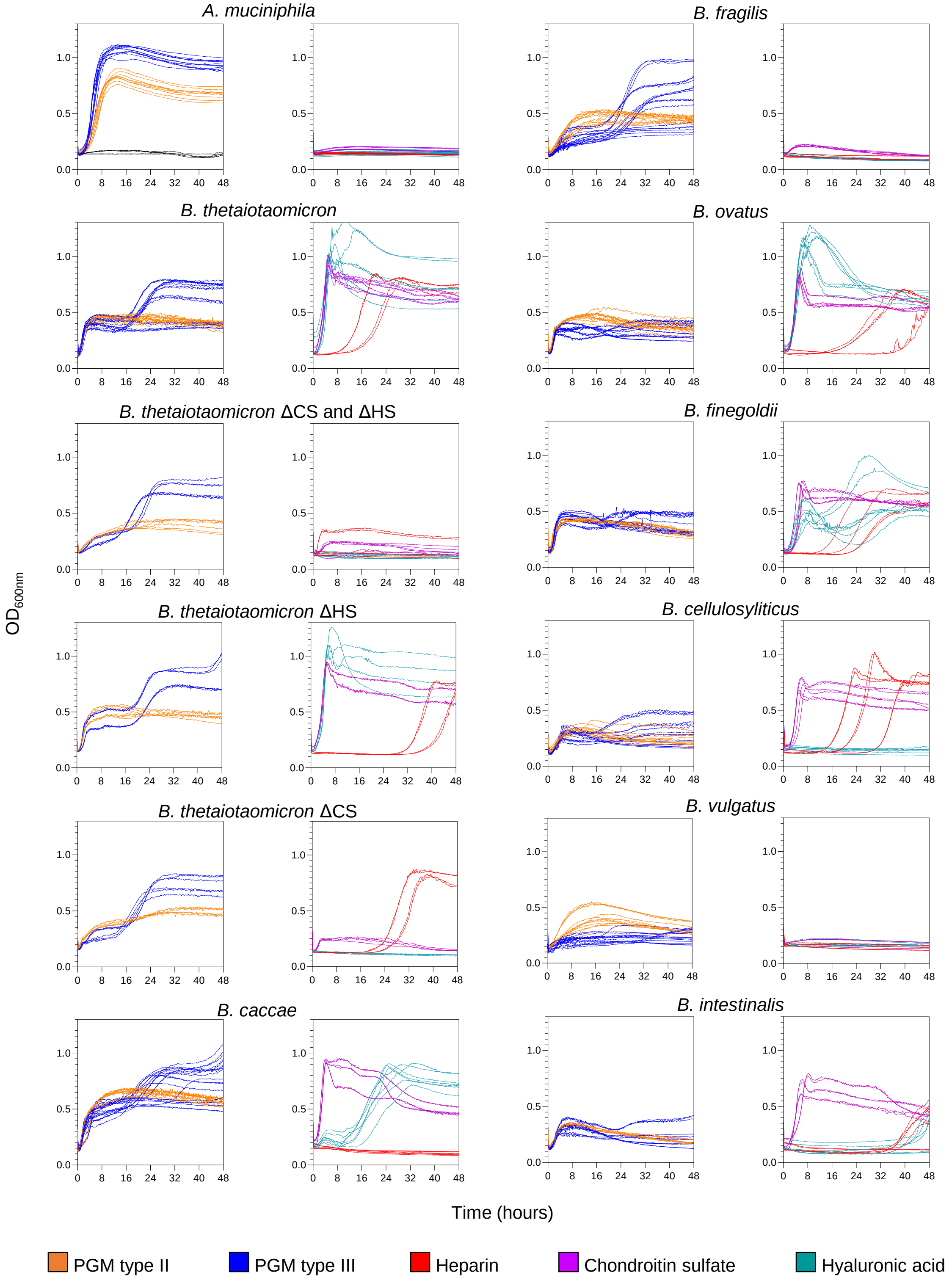


**Supplementary Figure 21│Growth of human mutualists of host glycans** *A. muciniphila* and different species of *Bacteroides* were grown on commercially available PGM type II, type III, Heparin sulfate, chondroitin sulfate and hyaluronic acid at 35, 40, 20. 20 and 10 mg/ml, respectively. Growth (OD600) was monitored continuously using a plate reader.

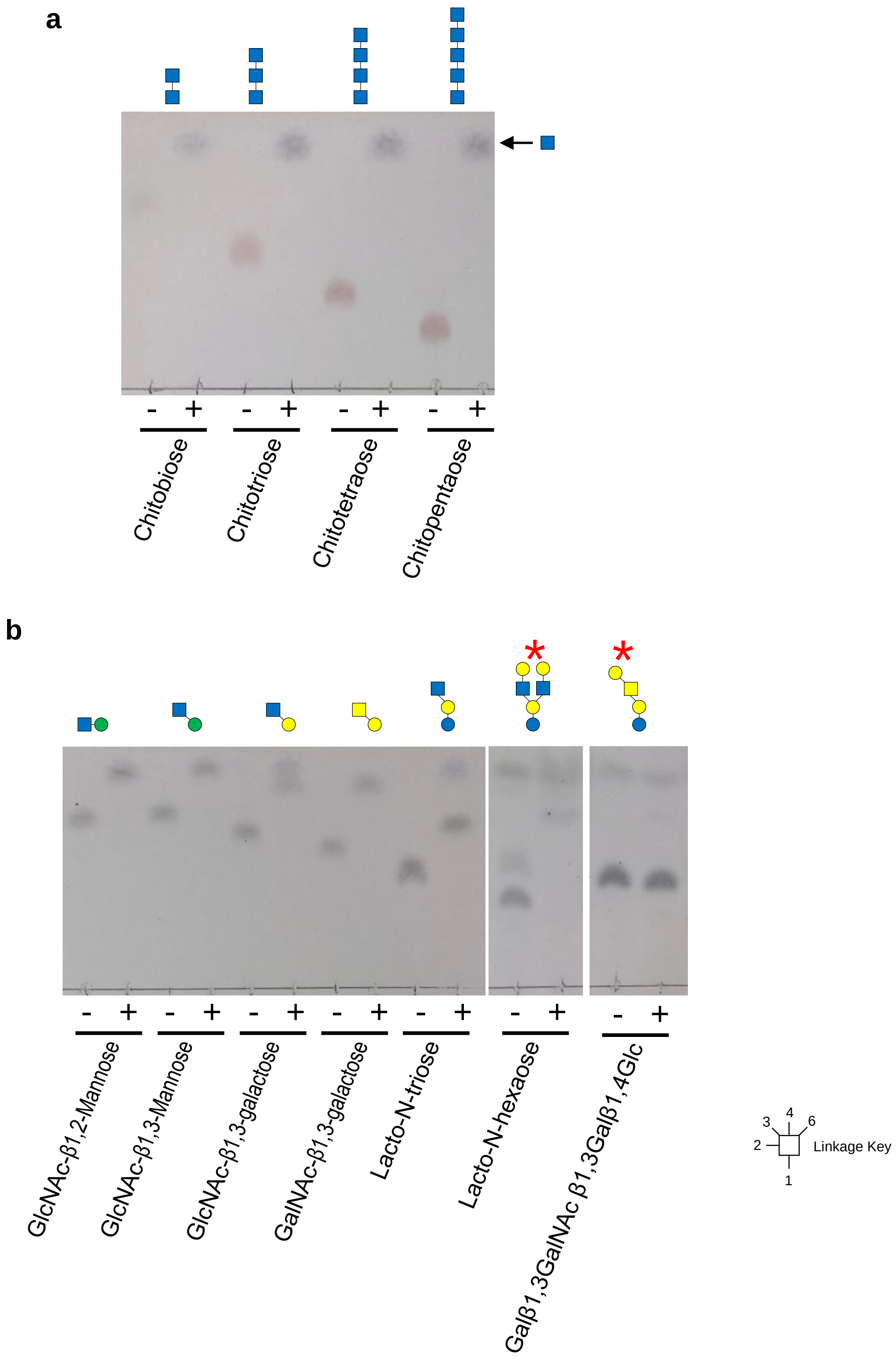


**Supplementary Figure 22│Activity of BF4059$^{GH20}$ against β-GlcNAc containing di- and oligosaccharides a,** Chitooligosaccharides. **b**, Other O- and N-glycan substrates were screened for activity. A red asterisk indicates lacto-N-hexaose and Galβ1,3GalNAc β1,3Galβ1,4Glc were pre-treated with BF4061$^{GH35}$ to remove the galactose before testing.
